# Unveiling shared genetic regulators for plant architectural and biomass yield traits in sorghum

**DOI:** 10.1101/2024.03.13.584802

**Authors:** Anuradha Singh, Linsey Newton, James C. Schnable, Addie M. Thompson

## Abstract

Sorghum is emerging as an ideal genetic model for designing high-biomass bioenergy crops. Biomass yield, a complex trait influenced by various plant architectural features, is typically regulated by numerous genes. This study aims to dissect the genetic mechanisms underlying fourteen plant architectural and ten biomass yield traits in a sorghum association panel (SAP) across two growing seasons. We identified 321 associated loci via genome-wide association studies involving 234,264 single nucleotide polymorphisms (SNPs). These loci encompass both genes with *a priori l*inks to biomass traits, such as ‘maturity’, ‘dwarfing (*Dw*)’, ‘*leafbladeless1’*, ‘cryptochrome’, and several loci not previously linked to roles in determining these traits. We identified 22 pleiotropic loci associated with variation in multiple phenotypes. Three of these loci, located on chromosomes 3 (S03_15463061), 6 (S06_42790178; *Dw2*), and 9 (S09_57005346; *Dw1*), exert significant and consistent effects on multiple traits. Additionally, we identified three genomic hotspots on chromosomes 6, 7, and 9, containing multiple SNPs associated with variation in plant architecture and biomass yield traits. Positive correlations were observed among linked SNPs close to or within the same genomic regions. Thirteen haplotypes were identified from these positively correlated SNPs on chr 6, with haplotypes 8 and 11 emerging as optimal combinations, exhibiting pronounced effects on the traits. Lastly, network analysis revealed that loci associated with flowering, plant heights, leaf characteristics, plant number, and tiller number per plant were highly interconnected with other genetic loci linked to plant architecture and biomass yield traits. The pyramiding of favorable alleles related to these traits holds promise for enhancing the future development of bioenergy sorghum crops.

## Introduction

The inescapable depletion of fossil fuels creates an urgent need to replace them with cellulosic biofuels, which have the potential to mitigate several undesirable aspects of fossil fuel utilization, such as greenhouse gas emissions and reliance on unstable foreign suppliers (USEPA, 2023). Optimized bioenergy crops represent one of the most sustainable and renewable resources for plant-based biomass production (Saleem, 2022). The cultivation of bioenergy crops is further accelerated by ethical considerations (Rosegrant and Msangi, 2014) and environmental concerns (Ain *et al*., 2022). This is particularly crucial when considering estimates that global energy demand is projected to rise due to population growth between now and 2050 (FAO, 2017).

A member of the Andropogoneae family, sorghum [*Sorghum bicolor* (L.) Moench], constitutes the world’s fifth most important cereal crop. It is grown for both grain and forage, serving as a vital food source in many food-insecure regions around the world (Silva *et al*., 2022). Because it requires less water and nitrogen than maize, sorghum can be grown with less associated greenhouse gas emissions and on marginal land not otherwise being used for food crops, creating great potential to meet the requirements of lignocellulosic biomass production (Damay *et al*., 2018; Ain *et al*., 2022; Mathur *et al*., 2017). Sorghum exhibits adaptability to a variety of soil types and environments, and lignocellulosic ethanol production from sorghum is predicted to yield five times more net energy per unit land area than ethanol production from grain starch and sugar alone, while emitting only a quarter of the greenhouse gases per unit energy (Mullet *et al*., 2014). As a result, improving high-biomass sorghum could significantly boost biofuel efficiency and renewable chemical production (Baloch *et al*., 2023).

The dry mass fraction of plant parts, such as stems, leaves, tillers, and panicles, is commonly called biomass and production of lignocellulosic biofuel increases with increasing biomass (Habyarimana *et al*., 2020; Habyarimana *et al*., 2022). Of this, stems and leaves account for 80-85% of harvested biomass from sorghum plants (Mullet *et al*., 2014). Plant traits weight, number, width, and length, can have a significant impact on the plant’s capacity to convert captured energy into biomass, and maintain physiological activity (Anami *et al*., 2015). The period from germination to flowering plays a critical role in determining the overall biomass productivity of different sorghum varieties. Later flowering, including delays produced in temperate environments as a result of photoperiod sensitivity, results in a longer vegetative growth period for sorghum accessions, and the production of more leading to increased stem and leaf biomass (Casto *et al*., 2019). Stem diameter, specifically thicker stems, is preferred for biomass yield (Kong *et al*., 2020). The physiological control of vegetative branching or tillers is essential in the deterministic breeding of optimized genotypes for sustainable cellulosic biomass production in both optimal and marginal conditions (Anami *et al*., 2015). Kong *et al*. (2014) reported that both the number of tillers with mature panicles and the number of immature secondary branches consistently showed positive correlations with total dry biomass production.

Biomass yield is a composite trait influenced by multiple plant architectural traits. These plant architectural traits are, in turn, frequently controlled by multiple quantitative trait loci (QTLs) with small effects. In many cases hundreds or thousands of genes each explain small portions of the contributing to variation in biomass production among different sorghum genotypes and the size and direction of the effects of individual genes can be different in different environments (Lasky *et al*., 2015; Miao *et al*., 2019; de Souza *et al*., 2021). To date, four major genes controlling plant height have been described in sorghum (*Dw1*, *Dw2*, *Dw3*, and to a less detailed extent *Dw4*) across various environments (Quinby and Karper, 1954; Hilley *et al*., 2017; Chen *et al*., 2019). Similarly, six Maturity (*Ma*) loci (*Ma1*, *Ma2*, *Ma3*, *Ma4*, *Ma5*, and *Ma6*) have been shown to control sorghum flowering time (Quinby and Karper, 1945; Murphy *et al*., 2011; Casto *et al*., 2019; Grant *et al*., 2023). The sorghum ortholog of the maize domestication gene *tb1*, a C2H2 zinc finger protein, and Dormancy Associated Protein 1 (DRM1, a well-known marker of bud dormancy) are known to regulate sorghum tiller number (Kebrom *et al*., 2006; Govindarajulu *et al*., 2021). Additionally, several growth regulators, such as phytohormones, play an important role in plant growth and development, contributing to biomass. For instance, auxin and brassinosteroid are associated with height (Hirano *et al*., 2017; Upadhyaya *et al*., 2013; Mu *et al*., 2022), while gibberellin and cytokinin are linked to stem biomass and diameter (Wang *et al*., 2020; Wang *et al*., 2022; Yu *et al*., 2022).

Understanding the genetic basis of complex quantitative and polygenic traits that contribute to variation in biomass productivity is crucial for advancing breeding of biomass sorghum cultivars. Genome-wide associated studies (GWAS), which rely on genetic linkage disequilibrium (LD), offer an opportunity to identify genes that affect the natural variation of quantitative traits by associating markers across the genome with phenotypic variation within diverse panels (Morris *et al*., 2013). Genomic analysis of diverse sorghum populations has been used to characterize the genetic determinants of traits including height (Brown *et al*., 2008; Murray *et al*., 2009), flowering time (Mace *et al*., 2013), panicle architecture (Brown *et al*., 2006; Wang *et al*., 2021), seed size (Tao *et al*., 2020; Zhang *et al*., 2023), photosynthesis (Ortiz *et al*., 2017), and carbon allocation between structural and non-structural carbohydrate (Murray *et al*., 2008; Brenton *et al*., 2016). In this study, leveraging a high-quality genomic sequence (McCormick *et al*., 2018) and GBS SNP data (Miao *et al*., 2020) for a sorghum diversity panel (Casa *et al*., 2008), loci contributing to variation in a wide range of plant architecture and biomass related traits were identified using genome-wide association studies. QTLs explaining significant variability across multiple biomass yield relevant traits represent suitable targets for marker-assisted selection to develop desirable and productive biomass sorghum cultivars.

## Material and Methods

### Plant material Site description and growth conditions

In this study, we utilized accessions from the Sorghum Association Panel (SAP), encompassing individuals from all five major sorghum botanical clades (bicolor, caudatum, durra, guinea, and kafir) and sourced from varieties originally collected across African, Asia, and America (Casa *et al*., 2008) (Data S1). The SAP panel accessions were grown in a randomized block design with two replications in each of two growing seasons, spanning from June to October in the year 2020 (391 accessions planted June 1^st^) and 2021 (396 accessions planted June 6^th^). The field design in each year also incorporated multiple replicates of the reference genotype BTx623 (PI 564163) (Paterson *et al*., 2009) within each block. Field experiments were conducted at the Michigan State University research field (42°42′53.5′′N, 84°27′46.5′′W; elevation of ∼265 m above sea level). The soil at the research site is classified as loamy with high available water capacity and moderate or moderately slow permeability. In each year, the experimental unit was a plot of two parallel 10-foot (approximately 3 meter) long rows separated by 30 inches (approximately 0.75 meters) with 3-foot (approximately 1 meter) alleyways separating sequential plots. Once the seeds were planted in the experimental field, weather data were checked every other day throughout the growing season (https://www.wunderground.com/history/monthly/us/mi/lansing/KLAN/date/2022-9). The incidence/amount of precipitation and temperature variations of each month in the experimental field from June to September are shown in Figure S1.

### Architectural- and biomass yield trait phenotyping

In both 2020 and 2021, data on twenty-four traits, comprising fourteen plant architectural traits and ten biomass yield traits (eight hand-harvested and two machine-harvested) were scored for each plot. The plant architectural traits included measurements related to plant height up to flag leaf (PH_FL; cm), plant height up to panicle (PH_Pani; cm), panicle length (PaniL; cm), days to 50% flowering (DOF; days), several leaf-related traits, such as largest leaf length (LLL; cm), width (LLW; cm), and area (LLA; cm^2^), the number of final green leaves (FGL; n), and the largest leaf number from the top (LLN_top; n). Additionally, stem-related traits, encompassing stem diameter (StD; mm) and volume (StV; cm^3^), along with counts of the plant (PlantN; n), panicle (PaniN; n), and tiller number per plant (TillN; n) within a specified area were measured (Figure 1).

**Figure 1.**
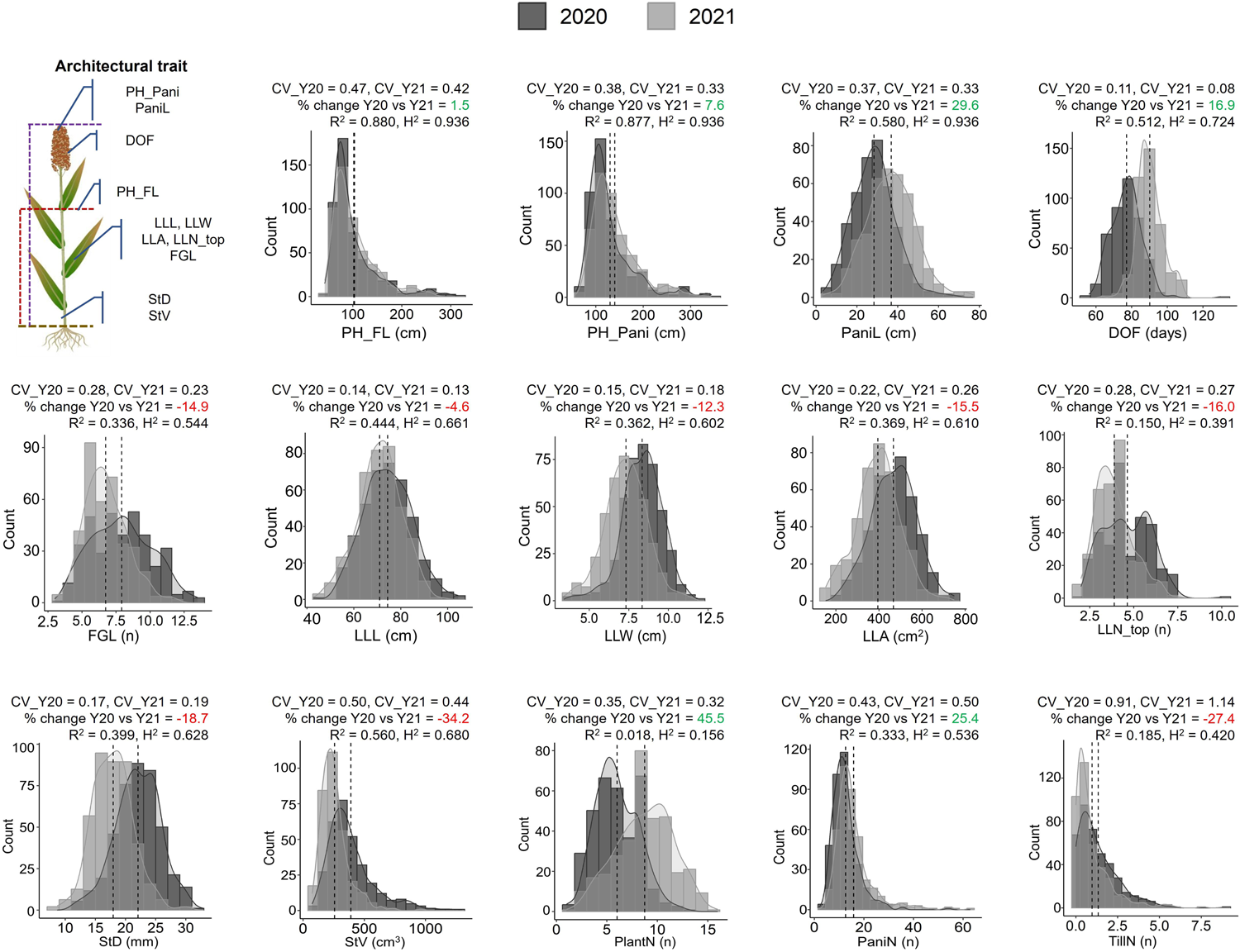
Phenotypic distribution of plant architectural traits in the Sorghum Association Panel (SAP) over two growing seasons (2020 and 2021). The mean value of each trait in each growing season was depicted as dotted lines on the respective plots. The coefficient of variations for each trait in each year was denoted as CV_Y20 and CV_Y21. The percentage (%) change in data acquisition between the years 2020 and 2021 (Y20 vs. Y21) was indicated, with gains shown in green and losses in red. The repeatability of data, measured through regression coefficient (R^2^, *P < 0.05*) and broad-sense heritability (H^2^) for each trait across growing seasons was presented on each distribution plot. The traits were abbreviated as follows: Plant height up to flag leaf (PF_FL), Plant height up to panicle (PH_Pani), Panicle length (PaniL), Days to 50% flowering (DOF), Final green leaves (FGL), Largest leaf length (LLL), Largest leaf width (LLW), Largest leaf area (LLA), Largest leaf number from top (LLN_top), Stem Diameter (StD), Stem Volume (StV), Plant number (PlantN), Panicle number (PaniN), and Tiller number per plant (TillN).

Days of flowering (DOF) for each plot was defined as the number of days between planting and the first date at which anthesis had occurred for at least half of the anthers of the primary panicles of at least 50% of the plants in that plot. When the plant reached the soft-dough stage, leaf-related traits, such as the number of final green leaves (those with 50% or more green surface area) on the main stem (FGL; n) and the largest leaf number from the top (LLN_top; n) were counted and marked. Then the largest leaf length (LLL; cm) and largest leaf width (LLW; cm) were measured using a ruler with a precision of 0.1 cm. The length was measured from the apex to the base of the leaf, while the width was measured at the widest part of the leaf. Once LLL and LLW were obtained, the largest leaf area (LLA; cm2) was calculated using the formula (LLL×LLW× 0.75) as described (Stickler *et al*., 1961).

Upon reaching physiological maturity, defined as when 90% of the grain had attained a black layer at the base of the grain (Eastin *et al*., 1973), plant height measurements were taken from the base of the stalk to the apex of the panicle (PH_Pani; cm), and the panicle length, from the node where the flag leaf joins the stem to the apex of the panicle (PaniL; cm). Plant height up to flag leaf (PH_FL; cm) was calculated using the difference between PH_Pani and PaniL. Stem diameter (StD; mm) was measured manually using a digital caliper at the thickest point of the stem in the region between 5 to 10 cm above the soil surface. Stem volume (StV, cm^3^) was calculated modeling the sorghum stem as a cylinder (V = πr^2^h) where r = (StD/2) and h = PH_FL. We also collected realized plant density as plant number (PlantN; n), counted panicle number (PaniN; n), and calculated tiller number per plant (TillN; n) all within a unit of area using a yardstick (91.5 cm) strategically placed in the center of each plot. PlantN was counted based on the number of main stems, while PaniN counted any panicles within the plot. TillN, a derived estimate of productive tiller number, was calculated using the formula (PaniN-PlantN)/(PlantN).

We also assessed ten biomass yield traits, eight of which were hand-harvested and two were machine-harvested. The hand-harvested biomass traits include the fresh weight (FW; g) and dry weight (DW; g) of both an entire single plant (SP_FW and SP_DW) as well as individual plant parts, specifically the stem (SPSt_FW and SPSt_DW), leaves (SPL_FW and SPL_DW), and panicle (SPPani_FW, and SPPani_DW) (Figure 2a). To collect these eight measurements, from each plot a single individual plant was selected and cut at the base of the stem. The fresh weight of the whole plant was recorded on a digital scale (accuracy, 0.1g). The same plant was then cut and divided into stems, leaves, and panicles. The fresh weights of each of these components were recorded. The samples were then over-dried at 60 °C for 5 days until reaching a consistent weight, and their dry weights were recorded. Single plant dry weight was defined as the total of SPSt_DW + SPL_DW + SPPani_DW.

**Figure 2a.**
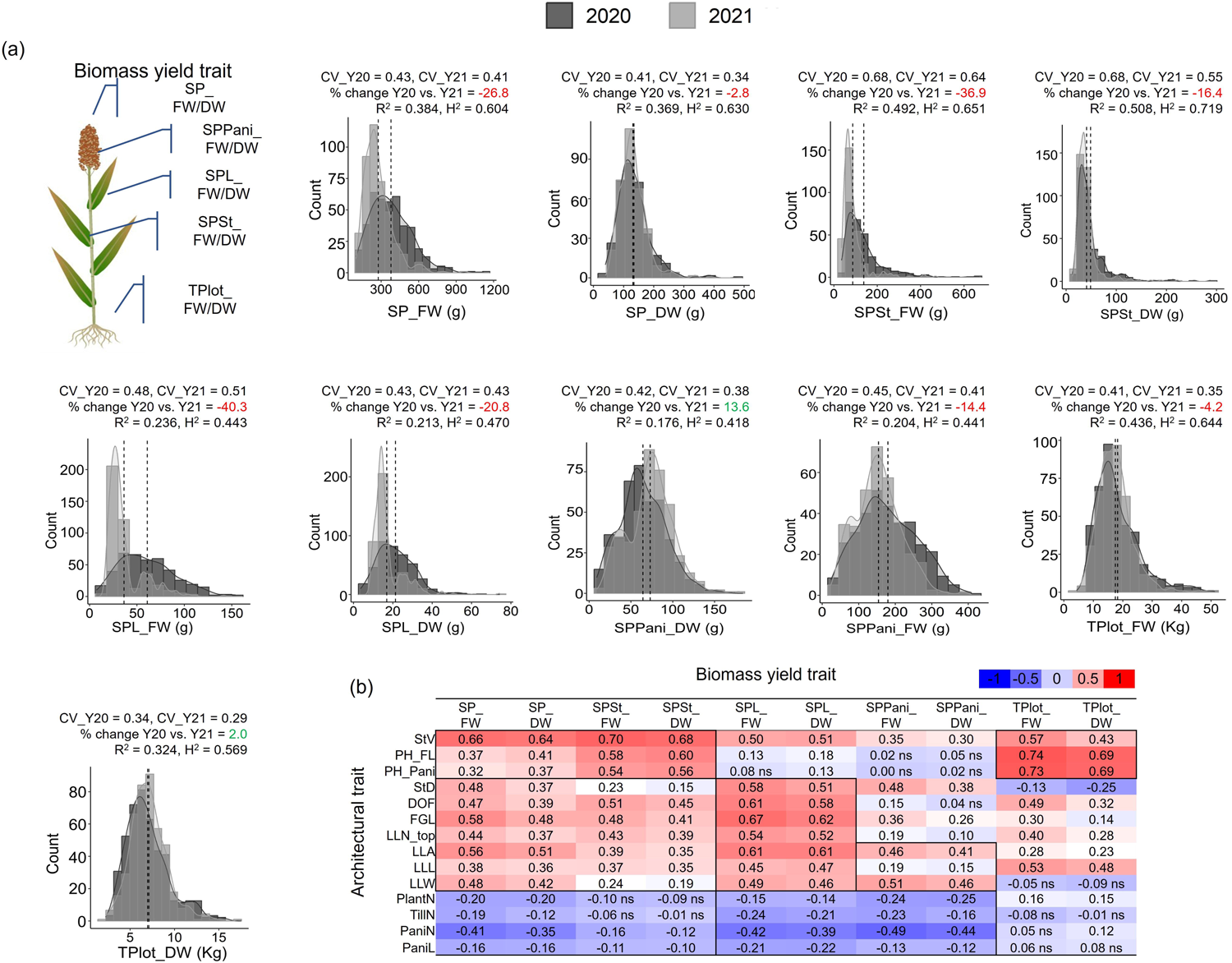
Phenotypic distribution of plant biomass yield traits in the Sorghum Association Panel (SAP) over two growing seasons (2020 and 2021). The mean value of each trait in each growing season was depicted as dotted lines on the respective plots. The coefficient of variations for each trait in each year was denoted as CV_Y20 and CV_Y21. The percentage (%) change in data acquisition between the years 2020 and 2021 (Y20 vs. Y21) was indicated, with gains shown in green and losses in red. The repeatability of data, measured through regression coefficient (R^2^, *P < 0.05*) and broad-sense heritability (H^2^) for each trait across growing seasons was presented on each distribution plot. **(b)** Pearson correlation analysis between plant architectural- and biomass yield traits in the year 2020. Positive correlations were indicated in red, while negative correlations were shown in blue. All correlations, whether positive or negative, were considered significant at *P < 0.05*, with non-significant correlations denoted as ‘ns’. The traits were abbreviated as follows: Single Plant_fresh weight and dry weight (SP_FW and SP_DW), Single plant-stem_fresh weight and dry weight (SPSt_FW and SPSt_DW), Single plant-leaves_fresh weight and dry weight (SPL_FW and SPL_DW), Single plant-panicle_fresh weight and dry weight (SPPani_FW and SPPani_DW), and Total plot_fresh weight and dry weight (TPlot_FW and TPlot_DW)

For machine harvesting, a biomass harvester was used to cut the entire plot of each accession. A portion of the chopped plot was immediately weighed as the sub-sample weight (SubS_FW; g). The remainder of the plot was also weighted and termed as the bucket weight. SubS_FW and bucket weight were then combined to obtain the total weight for the plot (TPlot_FW; kg). The sub-sample was subsequently dried at 60 °C for at least 5 days and the dry weight of sub-samples (SubS_DW; g) was recorded. Then with the help of available moisture calculated from SubS_FW and SubS_DW, TPlot_DW was determined and expressed in kg.

### Phenotypic data distribution, regression, and heritability analysis

Most statistical analyses of phenotypic data were carried out using the R statistical software (R Core Team, 2018). For instance, two-year data with individual year mean were visualized through skewed histograms using the “ggplot2” and “moments” packages. The coefficient of variation (CV) was measured through the ratio of standard deviation (σ) in the individual year trait data to its mean (μ) (CV = σ/μ). The repeatability of the data between years was evaluated using the regression coefficient (R^2^) value. Broad-sense heritability (H^2^) for all the captured phenotypic data was calculated based on the impact of genotype (G), environment (E), and genotype by environment interaction (G × E) using the “variability” package on the R. Different years (2020 and 2021) were treated as different environments. Genotype, environment, and their interactions (genotype × environment) were all considered as random factors. To analyze the effects of genotypes and two growing seasons on phenotypic traits, a two-way ANOVA was performed using the “res.aov()” R function. To determine the relationship between plant architectural and biomass-related traits, correlation analysis was performed using the phenotypic mean of the two replicates per year. Pearson correlations and the subsequent p-values were calculated using R statistical software with the “corr.test” function and plotted using Microsoft Excel.

### GWAS analyses

Genome-wide association studies were conducted using a previously published set of 569,305 high-confidence cleaned and imputed SNPs genotyped across the SAP using a modified tGBS (tuneable-genotyping-by-sequencing) protocol (Miao *et al*., 2020; Ott *et al*., 2017). SNPs were scored relative to the sorghum BTx623 reference genome v3.1.1 (McCormick *et al*., 2018), and filtering and imputation were previously described in Miao *et al*., 2020. Next, markers with less than 3% minor allele frequency were removed, yielding 234,264 SNPs scored across 358 sorghum accessions. For each trait, prior to GWAS analysis, outlier values were identified via the interquartile range (IQR; 25-75 percentile) method and values more than 1.5 times IQR below the 25 percentile or above the 75 percentiles were removed. The “rMVP” package (Yin *et al*., 2021) in R was then used to run GWAS using the Fixed and Random Model with Circulating Probability Unification (FarmCPU) algorithm (Liu *et al*., 2016). The first three principal components were fit as covariates to control for population structure and the kinship matrix computed internally by the FarmCPU algorithm were fit as random effects (Mural *et al*., 2021). Each GWAS model fit was assessed by examining the quantile-quantile (Q-Q) plots, and multiple trait Manhattan plots were created using the “CMplot” R package (v3.6.2) (Yin, 2020). To reduce the chance of false positives, significance levels were determined using a false discovery rate (FDR)-adjusted p-value threshold. The phenotypic variance explained (PVE) by each SNP was calculated as described (Kumar *et al*. 2021).

### Candidate gene identification and gene-set enrichment analysis in significantly associated regions controlling plant functional and biomass-related traits

Candidate genes were defined as those within 75 kb up- and downstream of a significant marker identified in the genome-wide association study. Gene ontology (GO) enrichment analysis for the set of all identified candidate genes was performed using PlantRegMap (plantregmap.gao-lab.org/). A GO term was considered significantly enriched in a gene set if the Fisher’s exact p-value < 0.05.

### Correlation and network visualization of the SNPs

Chromosome-wise Pearson correlation analysis was conducted on the identified SNPs. In brief, the allelic variations at each SNP were transformed into numeric values [REF (1), ALT (2), and heterozygous allele (1.5)]. Then, correlation analysis and its visualization were carried out using the R package “ggcorrplot”. Haplotype network analysis of selected SNPs and the visualization of the subpopulations were performed using the median-joining network interface in POPART (Population Analysis with Reticulate Tree) software. For network analysis among the SNPs, all possible pairwise correlation analyses and basic partial correlations were performed. The correlation network was then projected on Cytoscape (v 3.10.0), and the highest connectivity among SNPs was identified using the clustering coefficient algorithm on cytoHubba.

## Results

### Overview of plant architecture and biomass yield traits variability, heritability, and correlation

In this study, we measured fourteen plant architectural traits and ten biomass yield traits over two growing seasons, utilizing diverse sorghum accessions collected from various regions worldwide (Data S1). The sorghum accessions exhibited substantial natural variations in the quantified phenotypic data, presenting a positive skewed distribution (Figure 1, 2a, Data S2, Data S3). We also observed significant variations in temperature and precipitation during two growing seasons of sorghum in the university research field. Specifically, temperature and precipitation recorded a notable increase in 2021, rising by almost 4.5% and 52%, respectively, resulting in a warm and humid year (Figure S1). These variations in the growing season led to observable differences in the acquisition of plant architectural and biomass yield traits. For instance, plant height-related traits (PH_Pani), DOF, PlantN, and PaniN were overestimated by 7.6% to 45.4% in the year 2021 compared to 2020, while all leaf- and stem-related traits were underestimated by 4.6% to 34.2% (Figure 1). The variation in the quantification of architectural traits further corroborated their biomass yield traits, with most of them being underestimated in the following year than the previous year (Figure 2a).

The effect of accession and growing season on the acquisition of plant architectural and yield traits was tested using a two-way analysis of variance (ANOVA). We found a significant effect of accession on the phenotyped traits (Table S1). Furthermore, we observed a significant impact of growing season on phenotypic traits, with exceptions for three specific traits (PH_FL with a 1.45% change, SP_DW with a −2.79% change, and Tplot_DW with a 2.04% change), where the percentage variations in data collections were relatively modest and not significant at all (Figure 1, 2a, Table S1).

We then used the coefficient of variation (CV) to compare the phenotypic plasticity of traits across growing seasons. Notably, among plant architectural traits, DOF exhibited the lowest phenotypic plasticity, while TillN showed the highest in both growing seasons, which was supported by previous findings (Kim *et al*., 2010). Biomass yield traits, on the other hand, exhibited modest plasticity, ranging from 0.29 (Tplot_DW) to 0.68 (SPSt_FW/DW), indicating that plant architectural traits underlying biomass yield traits remained stable in different environments.

Furthermore, the two-year plant architectural and biomass yield data exhibited strong repeatability, with the maximum regression coefficient (R^2^) observed for PH_FL (0.88), while PlantN displayed the lowest (0.018). Similarly, the R^2^ values for biomass yield traits were moderate, ranging from 0.51 to 0.18 for SPSt_DW and SPPani_DW, respectively. Broad sense heritability (H^2^) for the plant architectural traits ranges from the lowest (PlantN; 0.156) to the highest (PH_FL; 0.936), while biomass yield traits showed moderate heritability with a range of 0.418 (SPPani_DW) to 0.630 (SP_DW).

We also conducted Pearson correlation analysis between plant architectural and biomass traits, independently in the two-year growing season data (Figure 2b, Figure S2). A similar correlation pattern was observed in both year’s phenotypic data; however, the magnitude of correlation values was slightly reduced in the year 2021 (Figure 2b, Figure S2). Most plant architectural traits (excluding PlantN, TillN, PaniN, and PaniL) exhibited significant positive correlations to biomass yield traits. For instance, strong positive correlations were observed between biomass traits (SP_FW/DW and SPSt_FW/DW, Tplot_FW/DW), primarily determined by plant height and stem volume. Leaf biomass was mostly associated with leaf architectural traits and DOF, influencing corresponding biomass yield by extending longer vegetative growth duration. In contrast, a negative correlation was observed between all the biomass and PlantN, TillN, PaniN, and PaniL, suggesting competition between the traits for resource utilization (Yang *et al*., 2019; Burgess and Cardoso, 2023). Taken together, it was concluded that these plant architectural traits serve as key drivers that affect biomass accumulation in sorghum.

### Genetic basis of plant architecture and biomass related trait in sorghum

We then performed independent Genome-Wide Association Studies (GWAS) analyses on fourteen architectural and ten biomass yield traits, captured from two growing seasons. The FarmCPU method was employed to detect significant SNP-trait associations, as it offers a favorable trade-off between power and FDR for moderately complex traits, with an increased likelihood of identifying rare causal variants (Miao *et al*., 2019). Manhattan plots and Q-Q plots (where observed p-values exceeded the expected p-values) corresponding to all plant architectural- and biomass yield traits, are presented in Figure S3.

A total of 321 significant SNPs were detected from the two-year phenotypic dataset (Figure S4a, Data S4). Nearly 38.0% of these SNPs were located on chr 6, 4, and 9, while the remaining 62.0% were distributed across the rest of the chromosomes (Figure S4b). Among these 321 SNPs, 101 fell within the cutoff range of 1.44 × 10^-40^ to 9.75 × 10^-9^ (p-value), while the remaining 220 were in the cutoff range of 1.10 × 10^-8^ to 2.4 × 10^-6^ (Figure S4c, Data S4).

On an individual year-wise, a total of 161 SNPs (p ≤ 2.4 × 10^-6^) were detected for a total of 23 traits, except for PlantN, which had no significant SNPs found in 2020-year phenotypic data (Figure S4d). In that year, leaf-related traits (LLL and LLW) exhibited the highest number of detected SNPs, while SP_FW has only one significant associated SNP. Similarly, excluding TPlot_FW/DW traits, the year 2021 discovered 160 SNPs for a total of 22 traits, with StV having the greater number of SNPs and Plant_N and SP_DW having the fewest (Figure S4d).

The minimum percentage of phenotypic variance explained by SNPs (PVP_min) across the traits ranged from 2.5% (TPlot_DW by S06_36711969) to 6.5% (PlantN by S10_50508696). Conversely, PVP_max ranged from 6.9% (TPlot_DW by S06_30488539) to 38.8% (PH_Pani, S09_57005346). It was interesting to note that SNP (S09_57005346) not only exhibited the greatest PVP_max for PH_Pani, but also for PH_FL (32.05%), StV (20.95%), and SPSt_DW (12.25%) (Data S4).

We then extracted candidate genes centered on each significant SNP, considering the LD decay around 150 kb. In total, 2773 candidate genes were identified (Data S5). These genes were predominantly associated with biological pathways related to single organism processes (GO:0044699), response to stimulus (GO:0050896), anatomical structure development (GO:0048856), gene expression (GO:0010467), macromolecule biosynthetic process (GO:0009059) (Figure S4e). The detailed description of genes underlying these pathways and their connections to plant architectural- and biomass yield traits is described below.

### Genetic analysis of plant architectural- and biomass yield traits revealed both known and novel regulators

#### Days of flowering (DOF)

Days of flowering (DOF) represents a pivotal agronomic trait, exerting a positive influence on both plant size and biomass at maturity (Habyarimana *et al*., 2020). A total of 20 SNPs associated with DOF were identified, colocalized with 199 genes (Figure 3, Data S4, Data S5). Remarkably, 50% of these SNPs were located on chr 4 and 6. Sorghum, being a short-day flowering plant, exhibits flowering in response to longer dark periods and correspondingly shorter days above a critical threshold (Rooney *et al*., 2007). Notably, six maturity loci (*Ma1-Ma6*) have been identified as key regulators of flowering time in sorghum (Quinby, 1966; Ge *et al*., 2023). Specifically, *Ma1*, located on chr 6, encodes a pseudo-response regulator (*SbPRR37;* Sobic.006G057866, Sobic.006G057900) and has the greatest influence on flowering time photoperiod sensitivity (Murphy *et al*., 2011). *Ma2* represents the Sobic.002G302700 gene on chr 2, encoding a SET and MYND (SYMD) domain-containing lysine methyl transferase (Casto *et al*., 2019). *Ma3* and *Ma5*, both located on chr 1, are encoded as phytochrome B (phyB, Sobic.001G394400) (Childs *et al*., 1997) and phytochrome C (phyC; Sobic.001G087100) (Rooney and Aydin, 1999), respectively. A fourth maturity locus (*Ma4*) was discovered in crosses of Milo (*Ma4*) and Hegari (*ma4*), but the corresponding underlying gene has not been identified (Quinby, 1966). *Ma6*, located on chr 6, encodes Ghd7 (Sobic.006G004400), a repressor of flowering in long days (a CONSTANS, CO-like and TOC1 (CCT)-domain protein) (Murphy *et al*., 2014).

**Figure 3.**
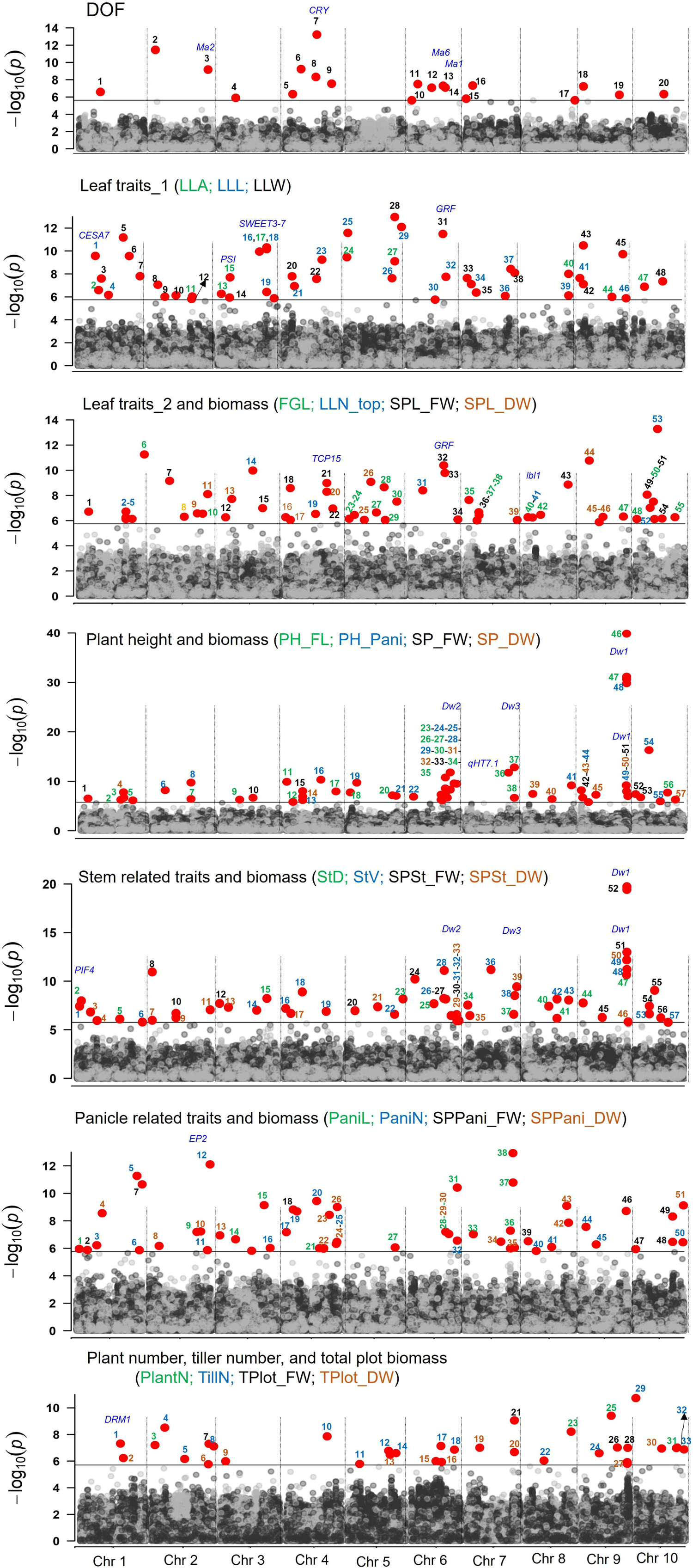
The Manhattan plot of significant SNPs identified from Genome-wide association studies (GWAS) for plant architectural and biomass yield traits in sorghum over two growing seasons. A total of 321 SNPs were identified from FarmCPU-based GWAS method. The plot shows -log10(p-values) of SNPs on y-axis, while SNPs were ordered by chromosomal location on x-axis. A grey line indicates the significance threshold (2.4 ×10^-6^). For a total of 24 traits, multiple manhattan plots are presented. Known genes are highlighted in blue. SNPs identified from different traits are color-coded and sequentially numbered.

In this study, we identified three SNPs located near previously reported loci, namely *Ma1*, *Ma2*, and *Ma6*. For instance, the SNP markers S06_40580184, S06_26501314, and S02_71011504 were found in proximity to Sobic.006G057866 (*Ma1* loci; 275.3kb upstream), Sobic.006G004400 (*Ma6* loci; 1253.68 kb downstream), and Sobic.002G302700 (*Ma2* loci; 3128.89 kb upstream), potentially representing the same QTL. The co-occurrence of *Ma1* and *Ma6* suggests an additive repressing action to enhance photoperiod sensitivity and delay flowering. This, in turn, results in a significant increase in leaf and stem biomass, as indicated by the positive correlation among the traits (Murphy *et al*., 2014).

We further identified the top two SNPs, one on chr 2 (S02_6231392) and another on chr 4 (S04_39191115), explaining the highest PVE ranging from 11.0% to 12.3%. The SNP (S04_39191115) does not colocalize with any known annotated gene within the LD region, however, the closest gene (Sobic.004G188400) is present nearly 14.8M downstream from this SNP. This gene encodes cryptochrome (CRY) and shares nearly 65.7% identity at protein with CRY of *Brassica napus*. This gene displayed early flowering at maturity in *BnCRY2* overexpressed transgenic plants (Sharma *et al*., 2022). On the other hand, SNP (S02_6231392) was present in the genomic region of the zinc ion-binding protein (Sobic.002G063900).

We also identified an additional five SNP markers, located in the genomic sequence region of glyoxalase, erythronate-4-phosphate dehydrogenase, RNA-dependent RNA polymerase, zinc ion binding protein, and ARF GTPase-activating domain-containing protein (Table S2). These genes may be considered as potential candidates for DOF colocalized SNP markers, which need further investigation to elucidate their specific impact and mechanism.

### Leaf-related traits (LLL, LLW, LLA, LLN_top, and FGL) and corresponding biomass (SPL_FW and SPL_DW)

Leaf-related traits, which include size, number, and stay-green characteristics, play a crucial role in determining plant fitness and their adaptation to environmental conditions (Yin *et al*., 2022). The shape and size of leaves are particularly important for plant architecture traits in cereals, as they affect planting density and light energy utilization through photosynthesis (Long *et al*., 2006). Our study identified 103 SNPs associated with these seven leaf-related traits, distributed across all chromosomes. Specifically, 18, 20, and 10 SNPs were linked to LLL, LLW, and LLA, respectively, while LLN_top, FGL, SPL_FW, and SPL_DW were associated with 10, 17, 15, and 13 SNPs, respectively. Notably, SNP related to LLW (S05_61965692; 13.5%), LLN_top (S10_27438731; 13.2%), and LLL (S05_70343006; 11.8%) explained the most PVE (Figure 3, Data S4, Data S5). Six out of these 103 SNPs stand out as pleiotropic loci that influence multiple leaf-related traits, as mentioned in Table 1.

**Table 1.** List of pleiotropic SNPs identified from two years of GWAS data.

| Trait | SNP | REF/<br>ALT | FarmCPU<br>FDR | PVE<br>(SNP) | Candidate genes |
| --- | --- | --- | --- | --- | --- |

| Year 2020 (12 SNPs) |  |  |  |  |  |
| --- | --- | --- | --- | --- | --- |
| SPSt_FW | S02_33697971 | T/C | 2.07E-07 | 6.38 | NA |
| SPSt_DW |  |  | 5.69E-07 | 5.52 |  |
| PH_FL | S02_48320641 | C/T | 1.87E-10 | 8.58 | Sodium/solute symporter |
| PH_Pani |  |  | 3.89E-07 | 5.53 |  |
| PH_FL | S04_20578520 | G/A | 9.75E-09 | 7.31 | Hydrolase |
| PH_Pani |  |  | 1.24E-07 | 6.42 |  |
| PaniN | S04_41002930 | G/A | 3.67E-10 | 9.58 | exonuclease |
| LLW |  |  | 2.74E-08 | 6.78 |  |
| SPL_FW | S04_6235481 | C/T | 2.64E-09 | 8.00 | CSLA1 - cellulose synthase-like family A |
| SPL_DW |  |  | 8.65E-07 | 5.28 |  |
| PH_Pani | S05_11983378 | C/A | 1.94E-10 | 9.50 | Glutathione S-transferase |
| PH_FL |  |  | 2.21E-06 | 4.32 |  |
| PH_Pani | S06_37640451 | G/A | 5.09E-08 | 7.07 | Zinc finger protein |
| PH_FL |  |  | 5.35E-07 | 5.31 |  |
| SP_DW | S06_48856570 | C/G | 4.98E-09 | 6.55 | Wax synthase, ABC transporter, glycosyl hydrolase |
| SP_FW |  |  | 1.60E-12 | 11.46 |  |
| Tplot_FW | S07_59146937 | C/T | 8.90E-10 | 8.69 | Phosphoglycerate mutase, oxidoreductase, Alpha-AMY |
| Tplot_DW |  |  | 2.11E-07 | 5.67 |  |
| StD | S08_40664971 | C/T | 6.90E-09 | 4.58 | NA |
| StV |  |  | 6.70E-07 | 3.40 |  |
| LLA | S08_57438310 | C/T | 1.01E-08 | 7.13 | Cycloartenol synthase 1 (BR |
| LLL |  |  | 7.57E-07 | 5.01 |  |
| PH_FL | S10_41292160 | A/C | 1.94E-08 | 7.24 | CSLD5 - cellulose synthase-like family D |
| StV |  |  | 1.72E-06 | 5.22 |  |
| Year 2021 (7 SNPs) |  |  |  |  |  |
| SPSt_FW | S02_4477084 | G/A | 1.15E-11 | 11.32 | Galactose-1-phosphate uridyl transferase |
| SPSt_DW |  |  | 1.08E-06 | 5.18 |  |
| LLL | S03_60633978 | G/T | 4.91E-11 | 10.33 | Polyprenyl synthetase SWEET2b |
| LLA |  |  | 3.77E-07 | 5.58 |  |
| SPL_DW | S04_51152508 | C/T | 1.04E-09 | 8.46 |  |
| SPL_FW |  |  | 5.16E-09 | 7.52 |  |
| LLW | S05_61965692 | A/C | 1.10E-13 | 13.51 | Glutamate synthase |
| LLA |  |  | 7.93E-10 | 8.75 |  |
| SPSt_DW | S09_56542041 | G/A | 2.23E-11 | 10.73 | Lipid phosphatase protein |
| SP_DW |  |  | 6.40E-10 | 9.28 |  |
| SP_FW | S09_58458241 | C/G | 1.04E-07 | 6.50 | Indole-3-acetic acid-amido synthetase |
| SPSt_FW |  |  | 1.53E-06 | 5.13 |  |
| PH_Pani | S10_18255972 | G/A | 4.90E-17 | 16.96 | □-glucosidase Aarabinogalactan |
| SPSt_FW |  |  | 3.72E-08 | 6.79 |  |
| StV |  |  | 2.35E-07 | 5.96 |  |
| SP_FW | S10_1887712 | C/A | 4.02E-08 | 7.15 | Glycosyl hydrolases Starch synthase |
| SPPani_FW |  |  | 1.15E-06 | 5.27 |  |
| Both year (3 SNPs) |  |  |  |  |  |
| SPL_DW (Y20) | S03_15463061 | T/C | 1.94E-08 | 7.01 | Endoglucanase, pectinesterase |
| LLA (Y21) |  |  | 2.04E-08 | 7.39 |  |
| PH_FL (Y20) | S06_42790178 | T/C | 2.75E-09 | 7.95 | Protein kinase (*Dw2*) |
| PH_Pani (Y20) |  |  | 3.28E-08 | 6.16 |  |
| PH_Pani (Y21) |  |  | 1.76E-11 | 10.72 |  |
| PH_FL (Y21) |  |  | 4.20E-08 | 5.96 |  |
| SPSt_FW (Y20) | S09_57005346 | G/A | 6.32E-12 | 10.87 | Aux/IAA (*Dw1*) |
| SPSt_DW (Y20) |  |  | 1.07E-13 | 12.25 |  |
| SP_DW (Y20) |  |  | 1.06E-08 | 6.94 |  |
| PH_FL (Y20) |  |  | 1.59E-30 | 29.98 |  |
| PH_FL (Y21) |  |  | 6.93E-32 | 32.06 |  |
| PH_Pani (Y20) |  |  | 2.29E-31 | 31.54 |  |
| PH_Pani (Y21) |  |  | 1.44E-40 | 38.85 |  |
| StV (Y20) |  |  | 3.76E-20 | 20.14 |  |
| StV (Y21) |  |  | 1.94E-20 | 20.95 |  |

The identified 103 SNPs were found to colocalize with 851 genes, representing those associated with phytohormone biosynthetic/degradation, transcription factors, photosynthesis, and metabolite biosynthesis. Among them, 14 genes showed the presence of SNPs within their genomic sequence. These genes were associated with various functions, including hormone degradation (e.g., Sobic.007G151400, which encodes cytokinin dehydrogenase), transcription regulation (e.g., Sobic.004G237300, representing TCP15), photosynthesis-related processes (e.g., Sobic.010G132100, encoding thioredoxin), and structural polysaccharide biosynthesis (Sobic.001G224300, CESA7, responsible for cellulose synthesis) (Table S2). Furthermore, we identified *a priori* candidate gene, Sobic.008G070600, the orthologue of maize *leafbladeless1* (lbl1, Zm00001eb264310) through colocalized SNP related to LLN_top (S08_9353971). The *lbl1* gene is responsible for a variety of leaf and plant phenotypes, including the specification of adaxial cell identity within leaves and leaf-like lateral organs (Nogueira *et al*., 2007).

Sorghum, being a C_4_ plant, possesses an efficient carbon concentrating mechanism, resulting in a higher carboxylation efficiency of rubisco (Ermakova *et al*., 2023). In this study, we observed the enrichment of photosystem I reaction center subunit XI (Sobic.003G052500) and thioredoxin (Sobic.010G132100) through S03_4722073 (LLA) and S10_18278152 (FGL) SNPs, respectively. This suggests that these genes could be potential candidates to improve leaf biomass. Once photosynthate, primarily sucrose is synthesized, it needs to be transported between various plant tissues through the phloem. The coordinated action of sucrose transporter (SWEET; Sugars Will Eventually be Exported Transporters and SUT; Sucrose Transporters) together with sucrose metabolic enzymes, regulates sugar content in the different tissues (Mizuno *et al*., 2016), and provides raw material for structural polysaccharide like cellulose (Persson *et al*., 2007). Cellulose, being the predominant component of plant cell walls, is produced by cellulose synthase complexes (CSCs), comprising numerous CESA proteins that generate individual glucan chains (Kumar and Turner, 2015). A total of 21-23 SWEET and 12 CESA genes were identified in *S. bicolor*. Among them, *SbSWEET3-7* (Sobic.003G269300) associated with pleiotropic loci S03_60633978, influencing both LLL and LLA, while CESA7 (Sobic.001G224300) was identified due to an SNP located within the gene and linked to LLL (S01_21486114).

Recent studies have demonstrated the significant role of both cytokinin and auxin in stay-green phenotypes in sorghum (Markovich *et al*., 2017; Borrell *et al*., 2022). Stay-green traits enable plants to retain green leaves and stems, thus maintaining longer growth during the end of the drought season (Borrell *et al*., 2000). Four major QTLs, namely *Stg1-4* derived from BTx642, have been identified. Detailed analysis revealed the presence of multiple PIN FORMED family of auxin efflux carrier genes within these QTLs, such as the *Stg1* (*SbPIN4*), Stg2 (*SbPIN2*), and *Stg3b* (*SbPIN1*) (Borrell *et al*., 2022). Similarly, cytokinin biosynthetic genes delay senescence in Arabidopsis and sorghum, probably due to increased activity of cell wall invertase (*CWINV*), a gene that affects the leaf source/sink balance (Markovich *et al*., 2017). In this study, the quantification of stay-green phenotype through FGL identified SNPs that were colocalized with flavin monooxygenase (Sobic.001G495850), auxin response factor 14 (ARF14; Sobic.009G196900), and cytokinin dehydrogenase (Sobic.007G151400.1) genes, indicating the possible involvement of these genes to regulating the stay-green phenotype through FGL in sorghum.

Among plant-specific transcription factors, members of the TCP family, known as growth suppressors and named after TEOSINTE BRANCHED1 (TB1) from *Zea mays*, CYCLOIDEA (CYC) from *Antirrhinum majus*, and PROLIFERATING CELL FACTORS (PCFs) from *Oryza sativa*, have been shown to play key roles in evolution of plant form and structure (Li, 2015). In sorghum, nearly 20 TCP genes have been identified, and all the TCP proteins are characterized by a conserved basic helix–loop–helix (bHLH) motif (Francis *et al*., 2016). In Arabidopsis, loss of function of *AtTCP* resulted in enlarged leaves and wrinkled leaf margins due to excessive cell division (Hervé *et al*., 2009). In this study, a SNP related to SPL_FW (S04_58517971) was identified in proximity to *SbTCP15* (Sobic.004G237300; 3.977 kb), suggesting an association with rapid regulation during cell division and cell differentiation, which ultimately affect the leaf biomass (Francis *et al*., 2016).

Furthermore, members of the GROWTH-REGULATING FACTOR (GRF) family require the transcription cofactors known as GIF to regulate downstream target genes (Kim, 2019). This interaction leads to a synergistic impact on various developmental processes in plants, including leaf size and longevity (Debernardi *et al*., 2014). In Arabidopsis, both the *gif* and *grf* mutant, as well as their combination, reduced the leaf size and cell numbers (Kim and Kende, 2004; Debernardi *et al*., 2014). On the contrary, overexpression of GIF1 and GRF3 promoted leaf growth (Debernardi *et al*., 2014). Eight putative GRF-encoding genes were identified in sorghum and designated as SbGRF1-SbGRF8 (Shi *et al*., 2022). In our study, pleiotropic SNP for LLA and LLW (S05_61965692) colocalized with Sobic.005G150900 (SbGRF6), located 19.28 kb upstream from the SNP. This suggests that this gene may play a vital role in regulating leaf growth and corresponding traits.

### Plant height (PH_FL and PH_Pani), stem diameter (StD), stem volume (StV) and corresponding biomass (SP_FW/DW and SPSt_FW/DW)

Plant height (PH) is an important agronomic trait, impacting not only whole plant biomass yield but also bolstering lodging resistance. PH is affected by the length of each internode (measuring the height of the stem from the ground to the flag leaf), the rate of internode production, and the duration of vegetative growth (Hilley *et al*., 2016). Further, the elongated internode, together with stem diameter, contributes to stem volume and stem biomass yield.

A total of 114 SNPs associated with these eight traits were identified. Remarkably, approximately 36.8% of these SNPs were exclusively located on chr 6 and 9. There were 22, 17, 11, and 19 SNPs associated with PH_FL, PH_Pani, StD, and StV, respectively. The remaining 8, 10, 13, and 14 SNPs were linked to SP_FW, SP_DW, SPSt_DW, and SPSt_FW, respectively (Figure 3, Data S4). It is also important to note that among these 114 SNPs, 14 exhibited pleiotropic loci, regulating more than one trait, as shown in Table 1.

These 114 SNPs were found to colocalized with 979 genes, representing the identification of a *priori* genes (Figure 3, Data S5). Plant height is known to be regulated by four dwarf loci, denoted as *Dw1*-*Dw4*, which control internode length (Quinby and Karper, 1954; Chen *et al*., 2019). These dwarf loci, *Dw1*, *Dw2*, *Dw3*, and *Dw4*, have been mapped to chr 9, 6, 7, and 4, respectively. The *Dw1* locus encodes a protein possibly involved in brassinosteroid signaling (Sobic.009G229800) (Yamaguchi *et al*., 2016). *Dw2,* encodes a protein kinase (Sobic.006G067700) (Hilley *et al*., 2017). *Dw3* is associated with an auxin transporter (ABCB1 auxin efflux transporter; Sobic.007G163800) (Multani *et al.,* 2003), while the causal gene for the *Dw4* locus has yet to be identified. Recently, another *dw5* mutant has been isolated which has a single recessive mutation in a single nuclear gene (Chen *et al*., 2019).

In the present study, we identified *Dw1* locus, which remained conserved across the year. This locus was detected through SNP (S09_57005346), intriguingly associated with multiple traits and explained the highest range of PVE (12.3% - 38.9%) for different traits. We also observed the presence of *Dw2* and *Dw3* loci, through S06_42790178 and S07_59944757 SNPs, explaining 10.7% and 12.7% PVE, respectively. Li et al. (2015) previously identified another plant height locus, *qHT7.1*, localized near the *Dw3* region on chr 7, which exerts a strong effect on plant height. In the present study, indeed we found another SNP (S07_52737620) for PH_FL which was in the close vicinity of *Dw3* loci and explained 11.0% PVE. Although genes underlying *qHT7.1* have not been identified, however considering the above SNP, the possible candidate genes could be *tasselseed-2* (Sobic.007G123000), which encodes a monocot-specific short-chain alcohol dehydrogenase known to affect plant height (Lunde *et al*., 2019).

We then identified other candidate SNPs that were associated with the highest PVP, which were found for StD (S09_56843188; 12.1%), StV (S07_32152737; 11.9%), SPSt_FW (S02_4477084; 11.3%), and SP_DW/SPSt_DW (S09_56542041; 9.3-10.7%). The colocalized genes around these SNPs were related to macromolecule biosynthesis. In sorghum, most stem biomass is in the form of primary metabolism (soluble sugars), cell wall composition (cellulose, and hemicellulose), and secondary metabolism (isoprenoid, flavonoid, phenol, and phenylpropanoids) (Hennet *et al*., 2020). Gene related to starch synthesis (Sobic.001G239500), starch branching enzyme (Sobic.006G066800), starch degrading enzyme (Sobic.006G063600), anthocyanin biosynthetic gene (Sobic.010G022700), and CSLD5-cellulose synthase-like family D (Sobic.010G146000) maintain the primary and secondary metabolism together with cell wall material in sorghum stem.

Hormones and transcription factors also play an essential role in stem development and corresponding biomass yield. For instance, SNP linked with StD (S01_5238883), colocalized with phytochrome-interacting factor 4 (PIF4), a bHLH transcription factor. PIF4 not only promotes the expression of auxin synthesis but also helpful in plant growth and development (Franklin *et al*., 2011). We also identified 30 genes that showed the presence of these SNPs in their genomic regions. Some of these genes were annotated as AAA-type ATPase family protein (Sobic.010G091400), phytochrome A (Sobic.001G111500.1), histone deacetylase (Sobic.006G067600.1, gene very close to *Dw2*), protein kinase (Sobic.006G067700; *Dw2*), dehydration-responsive element-binding protein (Sobic.006G082100), mitochondrial carrier protein (Sobic.009G249500.1; very close to *Dw1*), and many more (Table S2). These genes are believed to control both whole plant and stem biomass yield in sorghum, warranting further investigation.

### Tiller number per plant (TillN), plant number (PlantN), panicle number (PaniN), panicle length (PaniL), and corresponding biomass (SPPani_FW/DW and TPlot_FW/DW)

A total of 84 SNPs were identified across these eight traits, distributed throughout the chromosomes, with a higher abundance (∼26.19%) on chr 2 and 4. Specifically, 15, 4, 4, and 10 SNPs were linked to TillN, PlantN, TPlot_FW, and TPlot_DW, respectively. While 12, 16, 8, and 15 SNPs were found to be linked with panicle-related traits, such as PaniL, PaniN, SPPani_FW and SPPani_DW, respectively (Figure 3, Data S4). Furthermore, one SNP (S07_59146937) turned out to be pleiotropic, associated with both TPlot_FW and TPlot_DW (Table 1). These 84 SNPs colocalized with 1040 genes.

Tillering or vegetative branching is one of the most plastic traits that usually affect the biomass yield and grain yield in many crop plants (Kim *et al*., 2010). The interplay of hormones with transcription factors is known to be involved in promoting bud dormancy (Kebrom and Mullet, 2016). Two important transcription factor genes, tb1 (*SbTb1*, Sobic.001G121600) and gt1 (*SbGt1*, Sobic.001G468400) and Dormancy Associated Protein 1 (DRM1, Sobic.001G191200) are known to regulate axillary branching (Kebrom *et al*., 2006; Govindarajulu *et al*., 2021). In the present study, SNP related to TillN (S10_562834) had the greatest PVE (10.1%) and was colocalized with auxin-induced protein 5NG4 and several metabolic genes. However, SNP on chr 1 (S01_49505944), with a lower PVE (6.6%), may possibly colocalized with known DRM1 at roughly 32Mb upstream. Tiller numbers are generally influenced by factors such as the availability of water, light quality, and planting density (Yang *et al*., 2019). In our study, SNP related to PlantN (S08_56617080, 7.4%), TPlot_FW (S07_59146937, 8.7%), and TPlot_DW (S06_30488539, 6.9%), exhibited the highest PVP and colocalized with large number of gibberellin receptor genes (Sobic.007G156600, Sobic.007G157400, Sobic.007G157500 etc.) and primary metabolic genes (Sobic.007G157700, Sobic.007G158300). This is consistent with previous studies where overexpression of GA20-oxidases in Arabidopsis improves biomass production and alters the lignification of cell wall (Biemelt *et al*., 2004).

PaniN and PaniL are other important agronomic traits that determine the number of grains a panicle can hold, consequently affect the panicle biomass yield (Zhang *et al*., 2023). Remarkably, SNP related to PaniN (S02_75893499), PaniL (S07_59452509), SPPani_FW (S01_80437942), and SPPani_DW (S08_53008834) exhibited the highest PVP, which were 13.0%, 11.7%, 10.5% and 8.6%, respectively. These SNPs harbor the previously identified closet genomic region on chr 1 (59803397-19808620), 2 (73190000-73247000), 7 (8189476-8208789), and 8 (53337842-53434526) colocalized with pm1-2 and pm2-2, pr7-1, and pm8-1 QTL, respectively (Wang *et al*., 2021).

These genomic regions further identified the Sobic.001G311050, Sobic.002G374400, Sobic.007G072600, and Sobic.008G120200 genes, respectively (Wang *et al*., 2021). Among them, the Sobic.002G374400 gene, highly expressed in panicles and shares 77% similarity with *Erect Panicle2* (*EP2*) in indica rice, suggests a potential regulatory role in sorghum. EP2 encodes a novel plant-specific protein localized to the endoplasmic reticulum with unknown function (Zhu *et al*., 2010). The EP2 mutants have shorter panicle length, more vascular bundles, and a thicker stem than that of wild-type plants, creating an erect panicle phenotype in sorghum, likely due to panicle morphology regulation in both sorghum and rice may have similar mechanisms (Chen *et al*., 2015).

### Plant architecture and biomass yield traits regulated through a shared hub genomic regions and pleiotropic loci

Multiple plant architectural traits exhibited significant correlations to biomass yield traits at the phenotypic level in both growing seasons. As a result, we speculated that these traits might share common molecular regulators at the genetic level. To illustrate this, we mapped 321 significant SNPs onto the sorghum chr at a 150-kb interval, defining the mapped interval as the genomic region. In total, 158 genomic regions spanning across all the chromosomes were identified. Among them, 50 and 58 genomic regions were exclusively present in Y2020 (highlighted in light green) and Y2021 (highlighted in light blue), respectively, while the remaining 50 regions (highlighted in pink) were shared between both year’s SNP data derived from different architectural and biomass yield traits (Figure S5, Table S3). Among these regions, 19 were associated with no fewer than 4 traits (either plant architecture or biomass or both). Notably, three of these 19 regions, i.e., 93 (42227964 – 42444423, on chr 6), 113 (58344078 – 61456355, on chr 7), and 143 (56113408 – 58458241, on chr 9), were considered hotspot regions that colocalized almost all the plant architecture and biomass related traits (Figure 4a).

**Figure 4.**
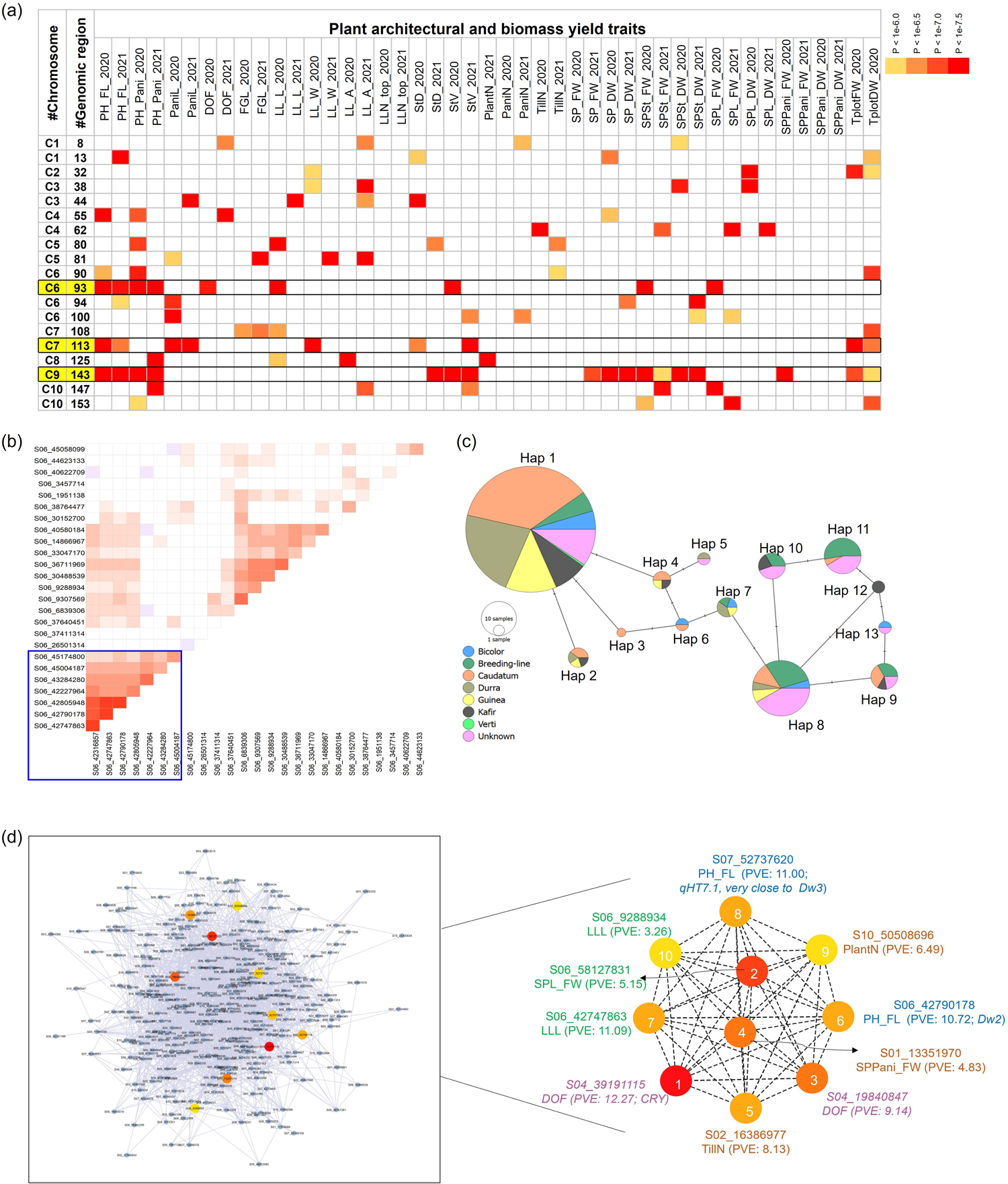
Genomic regions visualization, correlations analysis, haplotype analysis, and identification of highly connected SNPs for multiple plant architectural- and biomass yield traits in Sorghum Association Panel. **(a)** Distribution of significant SNPs in 19 genomic regions associated with no less than 4 traits (either plant architecture or biomass or both). These genomic regions were numerically coded, and details of these genomic regions were mentioned in Table S3. The significance of each SNP was color-coded based on P-value of the association, with white boxes representing no significant association. Solid boxes with yellow highlighting around specific genomic regions indicate hotspot areas exhibiting the presence of multiple SNPs related to plant architecture and biomass related traits. **(b)** Chromosome-wise correlation analysis among SNPs. The figure showed a correlation analysis specifically focusing on chromosome 6 SNPs. Positive correlations were indicated in red, while negative correlations were shown in blue. All correlations, whether positive or negative, were considered significant at P < 0.05, while non-significant correlations denoted as ‘white’. The purple box showed the SNP blocks with positive associations. **(c)** Haplotype analysis of six SNPs from the genomic region 93 and 94 on chromosome 6. A total of thirteen distinct haplotype variants were identified from the six SNPs. These haplotype variants were further explored at the level of their distribution among different sorghum races and their impact on phenotypic values. **(d)** Identification of the top ten highly connected SNPs through global correlation analysis among all Possible SNP Pairs. The global network was visualized using Cystoscope and the highest connectivity among SNPs was determined through clustering coefficient algorithm in cytoHubba. Node color from yellow to red, illustrating increasing connectivity.

Genomic region 93 contains six SNPs related to SPL_FW (S06_42227964), SPSt_FW (S06_42316857), LLL (S06_42747863), StV (S06_42805948), DOF (S06_43284280), with S06_42790178 acting as a pleiotropic regulator of plant height (PH_FL and PH_Pani) in both years. This genomic region was associated with the 1,4-alpha-glucan-branching enzyme, glycosyl hydrolases, together with protein kinases (*Dw2*), previously reported to regulate stem-related traits and plant height (Hilley *et al*., 2017).

Genomic region 113 has nine SNPs related to LLW (S07_58344078), PaniL (S07_59452509, S07_59524576, S07_61456355), PH_FL (S07_59944757, S07_59944889), StD (S07_60064545), StV (S07_61234352), together with S07_59146937 pleiotropic loci that regulate TPlotFW/DW. This region was also associated with another plant height regulating gene, *Dw3,* together with TCP, cyclins, etc., contributing to plant height, leaf- and panicle related traits, respectively.

Lastly, genomic region 143 contains seven SNPs that regulate 17 traits together. Of them, four SNPs were related to TPlot_FW (S09_56848978), TPlot_DW (S09_56113408), SPPani_FW (S09_56482255), StD (S09_56843188), while three SNPs were pleiotropic, including S09_58458241 (SP_FW, SPSt_FW), S09_56542041 (SP_DW, SPSt_DW), S09_57005346 (PH_FL, PH_Pani, StV, SP_DW, SPSt_FW/DW). This region was linked with genes related to photosynthesis and related metabolic processes (2Fe-2S iron-sulfur cluster, C4-dicarboxylate transporter, glycosyltransferase, oxidoreductase) and brassinosteroid signaling gene (*Dw1*), known to regulate height and biomass in sorghum (Hirano *et al*., 2017).

We also identified several pleiotropic loci that regulate multiple plant architectural and biomass yield traits (Table 1). Specifically, 13 and 8 pleiotropic loci were detected in the years 2020 and 2021, respectively. Of them, 12 and 6 pleiotropic loci predominantly colocalized with two different traits, while locus (S10_18255972) located on chr 10 from 2021-year data regulated three traits together (SPSt, PH_Pani, and StV). It was also interesting to note that from the two-year dataset, three pleiotropic loci were highly conserved, detected in both years. For instance, S03_15463061 regulating leaf related trait in both years and was linked with endoglucanase and pectinesterase genes, indicating cell wall metabolism for effective biofuel production. While genetic loci S06_42790178 and S09_57005346 regulate multiple plant height, stem volume, and related biomass yield traits, being linked to *Dw*2 and *Dw*1 genes, respectively (Hilley *et al*., 2016; Hilley *et al*., 2017).

### Pairwise correlation analysis among significant SNPs identified blocks of highly correlated markers

We next performed chromosome-wise Pearson correlation analysis among the identified significant SNPs. Within each chromosome, multiple pairs of highly correlated markers were identified (Figure S6). For instance, on chr 1, three SNPs related to LLN_top (S01_57545826, S01_57552937, and S01_57563841), a pair of SNPs on chr 4 associated with the trait PH_FL and PH_Pani (S04_20578520 and S04_20578555), another pair of SNPs regulating FGL and StD on chr 7 (S07_3517563 and S07_3522065) showed higher correlation with each other (r ≥ 0.85) (Figure S6). Since these highly correlated SNP were in the same genomic region, they can therefore be substituted as an alternative haplotype at the same locus.

We also identified several SNP blocks on chr 3, 6, 7, 8, 9, and 10, which although belonging to different genomic regions, exhibited moderate to high correlation (r > 0.50) (Figure S6). For instance, blocks of 8 SNPs on chr 6 from genomic regions 93 and 94 were associated with SPSt_FW, plant heights, StV, LLL, and DOF, leading to the identification of tightly linked *Ma1*/*Dw2* loci (Higgins *et al.,* 2014) (Figure 4b). Similarly, another block on chr 6 which contained 16 SNPs from genomic regions 83-92, was associated with DOF, StD, and StV, together with *Ma6* locus that regulate flowering. On chr 7, a block comprising 17 SNPs from genomic regions 108 −113 was identified that regulates multiple plant architectural traits together with *Dw3* locus that regulate plant height and derived biomass yield traits in sorghum (Li *et al*., 2015). Another block on chr 9 containing 15 SNPs from 131-143 genomic region was mostly associated with multiple plant architectural traits together with *Dw1* locus that regulate plant height, stem, and leaf biomass.

As correlation analysis identified highly correlated SNP blocks, we performed haplotype analysis on six SNPs [S06_42227964, S06_42316857, S06_42747863, S06_42790178 (*Dw2*), S06_42805948, and S06_43284280 (*Ma1*)] from the genomic region 93 and 94 to visualize the impact of allelic combinations on the traits (Figure 4c). A total of 13 haplotype combinations were identified. Among them, haplotype 1 emerged as the largest, comprising the REF alleles for all the selected SNPs, represented by caudatum, durra, and guinea race (Figure 4c). Haplotype 8 was the second largest, carrying four ALT alleles and being predominantly represented by breeding lines. Haplotype 11 stood as the third largest, carried ATL alleles for all the selected SNPs, and was mostly represented by breeding lines. We also observed the noticeable differences in the quantified data of SPL_FW, SPSt_FW, LLL, PH_FL, StV, and DOF as transitioned from haplotype 1 to haplotype 11 (Table 2). Taken together, the evidence suggests that the classical flowering (*Ma1*) and dwarfing alleles (*Dw2*) were likely introgressed into breeding lines that collectively affect biomass yield traits (Higgins *et al*., 2014).

**Table 2.** Haplotype analysis and visualization of traits across all haplotypes. Haplotype analysis was conducted using SNPs within the genomic region 93 and 94 on chromosome 6.

| Haplotype number | Haplotype sequence | # Accession | SPL_FW (g) | SPSt_FW (g) | LLL (cm) | PH_FL (cm) | StV (cm<sup>3</sup>) | DOF (days) |
| --- | --- | --- | --- | --- | --- | --- | --- | --- |
| Hap 1 | GCATTT | 211 | 47.36 | 97.53 | 70.76 | 89.30 | 292.64 | 83.16 |
| Hap 2 | GCACTT | 5 | 48.37 | 158.62 | 76.27 | 140.03 | 456.95 | 87.35 |
| Hap 3 | GCTTTT | 1 | 62.38 | 129.81 | 88.13 | 79.88 | 354.64 | 92.00 |
| Hap 4 | GTATTT | 4 | 34.20 | 65.71 | 70.00 | 87.56 | 241.07 | 81.06 |
| Hap 5 | TTATTT | 2 | 52.13 | 91.43 | 75.13 | 62.56 | 285.79 | 86.63 |
| Hap 6 | GTTTTT | 2 | 38.53 | 98.45 | 74.38 | 78.50 | 228.62 | 82.88 |
| Hap 7 | GTTCTT | 5 | 60.96 | 177.48 | 68.28 | 106.35 | 238.71 | 78.40 |
| Hap 8 | G TTCCT | 41 | 56.45 | 140.36 | 77.40 | 127.60 | 401.91 | 85.05 |
| Hap 9 | GCTCCT | 9 | 44.80 | 89.61 | 75.29 | 92.28 | 317.51 | 82.22 |
| Hap 10 | TTTCCT | 9 | 45.78 | 107.99 | 71.79 | 114.54 | 320.11 | 82.39 |
| Hap 11 | TTTCCC | 17 | 59.34 | 118.59 | 79.18 | 94.03 | 337.55 | 85.90 |
| Hap 12 | GTTCCC | 2 | 35.62 | 95.40 | 83.88 | 136.38 | 370.42 | 88.75 |
| Hap 13 | GCTCCC | 2 | 32.46 | 113.29 | 69.31 | 120.88 | 277.99 | 83.88 |

### Network analysis and overrepresented SNPs identified from GWAS of multiple plant architectural- and biomass yield traits

We also conducted a network analysis to identify the top 10 highly connected SNPs from our GWAS analyses. Before this, we performed pairwise correlation analysis and basic partial correlation among all possible SNP pairs. Subsequently, a correlation network was projected onto Cytoscape (v 3.10.0). The highest connectivity among SNPs was identified using the clustering coefficient algorithm on cytoHubba (Figure 4d). The correlation coefficient cutoff at *p* < 0.05 was *r = 0.231*. Notably, 60% of highly connected SNPs were located on chr 4 and 6, with the remaining 40% distributed across chr 1, 2, and 10. Among the SNPs, those related to DOF with the highest PVE [S04_39191115 (CRY) and S04_19840847] ranked 1^st^ and 3^rd^, implying that DOF is a crucial contributor to biomass (Casto *et al.,* 2019). Another large-effect SNPs related to plant height (S06_42790178; *Dw2,* S07_52737620; *qHT7.1,* very close to *Dw3*), and LLL (S06_42747863) ranked 6,7, and 8^th^ respectively. This finding aligns with a previous report where genes regulating plant height were shown to enhance biomass in sorghum (Hilley *et al*., 2016). Besides large-effect SNP, plant architectural traits and biomass yield traits are usually contributed by small-effect SNPs. Indeed, our analysis identified the SNP with a smaller effect related to SPL_FW (S06_58127831; 2^nd^), SPPani_FW (S01_13351970; 4^th^), TillN (S02_16386977; 5^th^), PlantN (S10_50508696; 9^th^), and LLL (S06_9288934; 10^th^). These small-effect SNPs work in concert with other SNPs underpinning complex traits like biomass.

## Discussion

Biomass yield is a complex trait comprising multiple plant architectural traits of various organs. It is typically regulated by many genes, although many of them have relatively small effects, with environmental factors often play a significant role (Miao *et al*., 2019; de Souza *et al*., 2021). Sorghum, a member of the Andropogoneae family, stands out as a preferred bioenergy crop. This is attributed to its diploid nature, extensive breeding history, substantial natural diversity, and a small genome of 750 Mb, making it a functional model for other Andropogoneae (Paterson *et al*., 2009). Identifying genomic regions contributing to biomass yield not only aids in the discovery of key genes but also facilitates the design of bioenergy crops that can adapt to changing climate conditions (Ain *et al*., 2022).

We employed a genome-wide association study (GWAS) to identify genomic regions and genes significantly associated with plant architectural traits and biomass yield traits over two growing seasons in sorghum. We utilized the power of Sorghum Association Panel (SAP) which was selected to maximize the genetic and phenotypic diversity of the panel (Casa *et al*., 2008). The SAP panel comprises temperate-adapted breeding lines and converted (photoperiod-insensitive) tropical accessions from the Sorghum Conversion Program (Klein *et al*., 2008). Notably, the SAP panel differ from other panels, such as the Bioenergy Association Panel, which was restricted to tall, photoperiod-sensitive, late-maturing accessions (Brenton *et al.,* 2016), or any of the multi-parent populations (Bouchet *et al*., 2017; Boatwright *et al*., 2021). The SAP was originally genotyped using simple sequence repeat markers (Casa *et al*., 2008) and later sequenced using a modified tGBS (tuneable-genotyping-by-sequencing) protocol (Ott *et al*., 2017; Miao *et al*., 2020), resulting 569,305 high-density genome-wide single-nucleotide polymorphism (SNP) markers. In this study, we utilized these SNP markers to expand our knowledge of genes that regulate biomass yield and plant architectural traits in sorghum, which may be useful for selecting genotypes with desirable traits for use in sorghum breeding programs.

Both plant architectural and biomass yield traits showed significant variations across accession, as indicated by their CV, which was highest for TillN and lowest for DOF in both years data collection. This indicates that plastic response to seasonal growth conditions strongly influences TillN, while DOF remains stable (Figure 1, 2a). A substantial variation across two growing seasons notably influences all traits, except PH_FL, SP_DW, and TPlot_DW, as evidenced by the lowest percentage change in the two-year data acquisitions. Furthermore, most of the plant architectural and biomass yield traits demonstrated high heritability, specifically plant heights and DOF, suggesting a strong genetically controlled regulation and amenability to selection (Naoura *et al*., 2019). Overall, the SAP panel demonstrates substantial phenotypic diversity in terms of both plant architecture traits and biomass yield traits, providing a robust foundation for GWAS analysis.

Our GWAS identified a total of 161 and 160 significant SNPs associated with plant architectural and biomass yield traits across two growing seasons. These SNPs were primarily distributed on chr 4, 6, and 9. Biomass, being a complex trait, showed high polygenicity, predominantly influenced by numerous small-effect polymorphic loci. In our study, only 11.1% of SNPs had large effects (ranging from 10% to 38.4%) on the phenotypic traits (especially plant height and StV). The remaining 88.9% had small effects that ranged from 2.53% to 9.99% (Data S4), which was consistent with previous finding (Habyarimana *et al*., 2020).

Correlation analysis between plant architectural and biomass yield traits revealed consistent patterns, indicating the stability of these traits across growing season (Figure 2b, Figure S2). A positive correlation between plant architectural and biomass yield suggests causal relationships between traits, shared genomic regions, and underlying SNPs/genes with pleiotropic effects. Indeed, several pleiotropic loci were identified, regulating at least two phenotypic traits together, suggesting a shared genetics mechanism between closely related traits from various organs that contribute to biomass yield (Mural *et al*., 2021).

Although dwarf varieties of rice and wheat made a great contribution to feeding people worldwide during the green revolution (Ferrero-Serrano *et al*., 2019). Additionally, the timing of flowering is a crucial agricultural trait, not only for successful reproduction, but also for the appropriate balance between vegetative growth and reproductive growth duration. Sorghum, being a promising feedstock for biomass, presents a challenge in designing an ideal sorghum ideotype. Tall varieties with photoperiod-sensitive sorghum can produce substantial amounts of aerial lignocellulosic biomass, serving as a sustainable and economically feasible feedstock.

The genetic basis of DOF and plant height in sorghum has been extensively studied, regulated through many maturity genes (*Ma1-Ma6*) and four dwarfing loci (*Dw1*-*Dw4)* (Casto *et al*., 2019; Grant *et al*., 2023). Our GWAS analysis identified at least three known regulators (*Ma1*, *Ma2*, and *Ma6*), along with the identification of novel loci (S04_39191115; CRY) that regulate flowering (Sharma *et al*., 2022). Furthermore, all three *Dw1*, *Dw2*, *Dw3* and *qHT7.1* which were known true positive height genes were identified that regulate plant height through elongated stem internodes in sorghum. *Dw1* (S09_57005346), considered as a conserved large effect pleiotropic locus, not only regulating plant height but also stem volume and biomass yield traits (Breitzman *et al*., 2019; Mural *et al*., 2021). We also observed a strong linkage between *Ma1* (Sobic.006G057866) and *Dw2* (Sobic.006G067700), both exerting a significant effect on sorghum flowering and final height (Figure 6B). In many cases, both loci are introgressed together during the backcrossing and conversion process of adapting tropical sorghum germplasm to grow in temperate latitudes (Quinby and Karper, 1954; Higgins *et al*., 2014).

The two-year SNP data showed substantial overlap in genomic regions and loci, regulating multiple plant architectural and biomass yield traits (Figure 4a, Figure S5, Table S3). Delayed flowering or prolonged vegetative growth in forage crops leads to increased leaf and stem biomass production (Rooney *et al*., 2007). In our study, genomic region 93 colocalized with multiple SNPs related to leaf feature and flowering *Ma1* loci (PRR37). The Ma1 loci not only enhance photoperiod sensitivity and delay flowering until daylengths are <12.3 h under field conditions, but also maintain photosynthetic capacity for longer, and greatly increasing leaf and stem biomass yield (Murphy *et al*., 2011)

Some plant architectural traits (PaniL, PaniN, TillN, PlantN) exhibited a negative correlation with all the biomass yield traits (Figure 2b, Figure S2), indicating a competition and antagonistic selection with biomass traits (Yang *et al*., 2019; Burgess and Cardoso, 2023). This observation was substantiated at the genetic level where SNP related to TillN on chr 5 (S05_15271748) showed trade-off with other biomass yield traits. In various cereals, a wide range of tiller numbers has been observed, spanning from high tiller counts in crops like wheat, rice, and barley, to lower tiller numbers, sometimes as few as zero to four fertile tillers in sorghum. This observation contrasts with many high-tillering cereals such as wheat, rice, and barley, which significantly contributed to improved grain yield during the green revolution (Sakamoto and Matsuoka, 2004). Planting density is another important factor that contributes to biomass in sorghum. SNPs related to PlantN on chr 9 (S09_19570441 with lowest PVP) and 10 (S10_50508696) consistently exhibited a negative correlation with several biomass yield traits.

The ability to identify both large- and small-effect loci associated with a phenotype of interest is paramount for breeders. Network analysis has highlighted both larger- and smaller-effect SNPs related to flowering (CRY), plant heights (*Dw2* and *qHT7.1*), leaf related traits, plant number and tiller number per plant, serving as major contributors that exhibit connections with other genetic loci related to plant architecture and biomass yield traits (Figure 4d). These identified loci are crucial for gaining a comprehensive understanding of the genetic regulation of complex traits of economic relevance, such as biomass and can be effectively utilized in the development of specialized plant feedstocks for bioenergy in sorghum.

## Conclusion

Our work offers insight into the genetic basis of multiple plant architectural- and biomass yield traits in sorghum, serving as a foundation for further functional investigation. We observed natural variation in individual traits and identified the well-characterized and novel loci that play a role in regulating these traits. Additionally, we identified conserved pleiotropic loci, shared genomic regions, highly correlated SNP blocks, and top connected SNPs that influence multiple plant architectural- and biomass yield traits in sorghum. Particularly, loci and significant SNP markers related to plant height, days of flowering and leaf related traits are of great interest. The strategic combination of favorable allele related to these traits through pyramiding will be beneficial for designing bioenergy sorghum crops.

## Data Availability Statement

All datasets will be available upon request.

## Acknowledgements

We acknowledge financial support from the U.S. Department of Energy, Grant no. DE-SC0020355 to JCS and AMT, as well as startup funds and project support from Michigan State University and the Plant Resilience Institute. We acknowledge and thank the many undergraduate students from the Thompson Laboratory at Michigan State University who assisted with field maintenance and data collection.

## Author contributions

AS performed the research, data collection, data analysis and interpretation. LN performed research and data collection. JCS provided initial seed stock and genotypic data and assisted in data analyses and interpretation. AT planned and designed the research study, and assisted in data collection, data analysis, and interpretation. All the authors contributed to the writing and revision of the manuscript.

