## Supplemental Figures for "Unveiling shared genetic regulators for plant architectural and biomass yield traits in sorghum"

**Supplementary files**

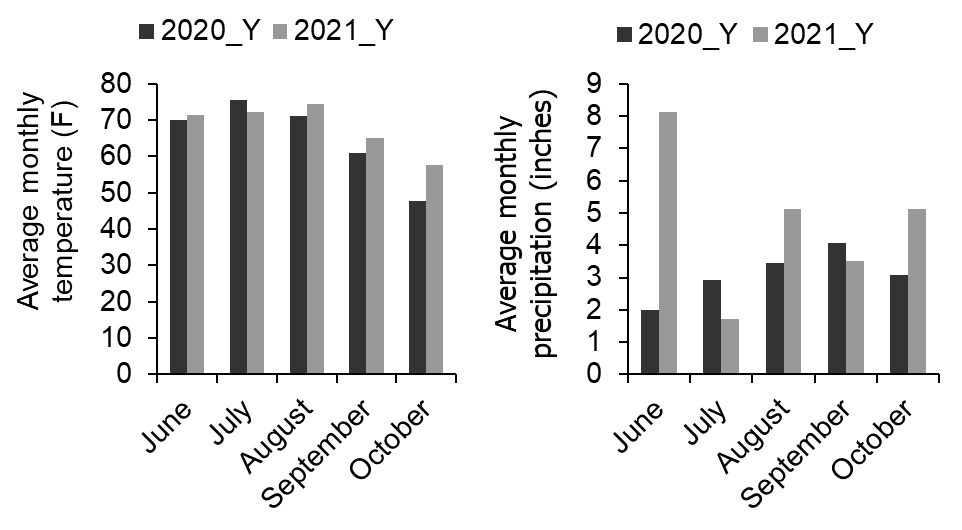

**Figure S1.** **Graphical representation of average monthly temperature (°F) and average monthly precipitation (inches) over two growing seasons of sorghum at the research farm of Michigan State University.** The data was obtained from this site. <https://www.wunderground.com/history/monthly/us/mi/lansing/KLAN/date/2022-9>.

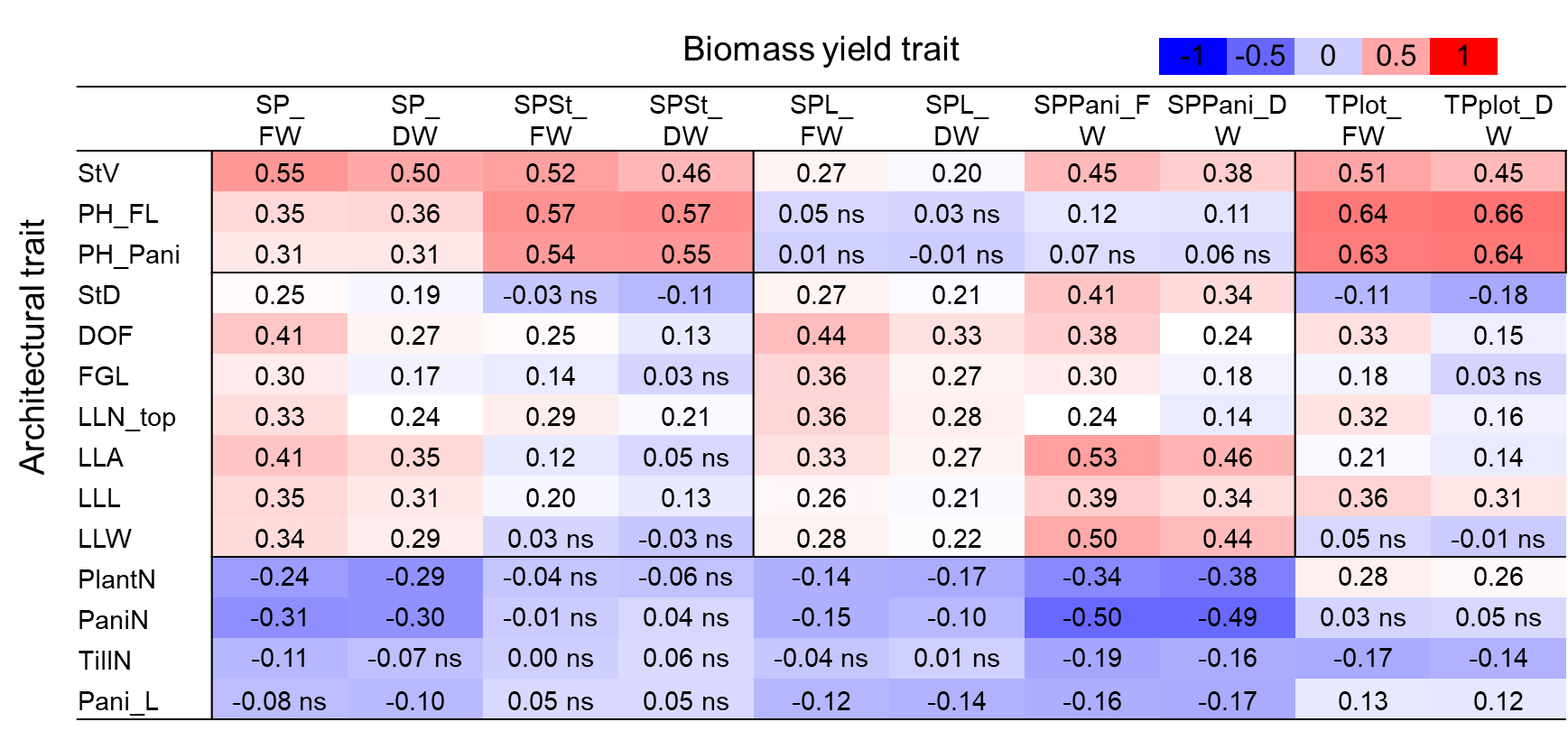

**Figure S2. Pearson correlation analysis between plant architectural- and biomass yield traits in the year 2021.**

Positive correlations were indicated in red, while negative correlations were shown in blue. All correlations, whether positive or negative, were considered significant at P < 0.05, with non-significant correlations denoted as 'ns’. The traits were abbreviated as follows: Single Plant_fresh weight and dry weight (SP_FW and SP_DW), Single plant-stem_fresh weight and dry weight (SPSt_FW and SPSt_DW), Single plant-leaves_fresh weight and dry weight (SPL_FW and SPL_DW), Single plant-panicle_fresh weight and dry weight (SPPani_FW and SPPani_DW), and Total plot_fresh weight and dry weight (TPlot_FW and TPlot_DW).

| 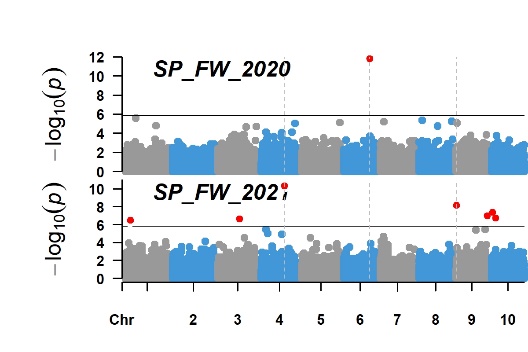 | 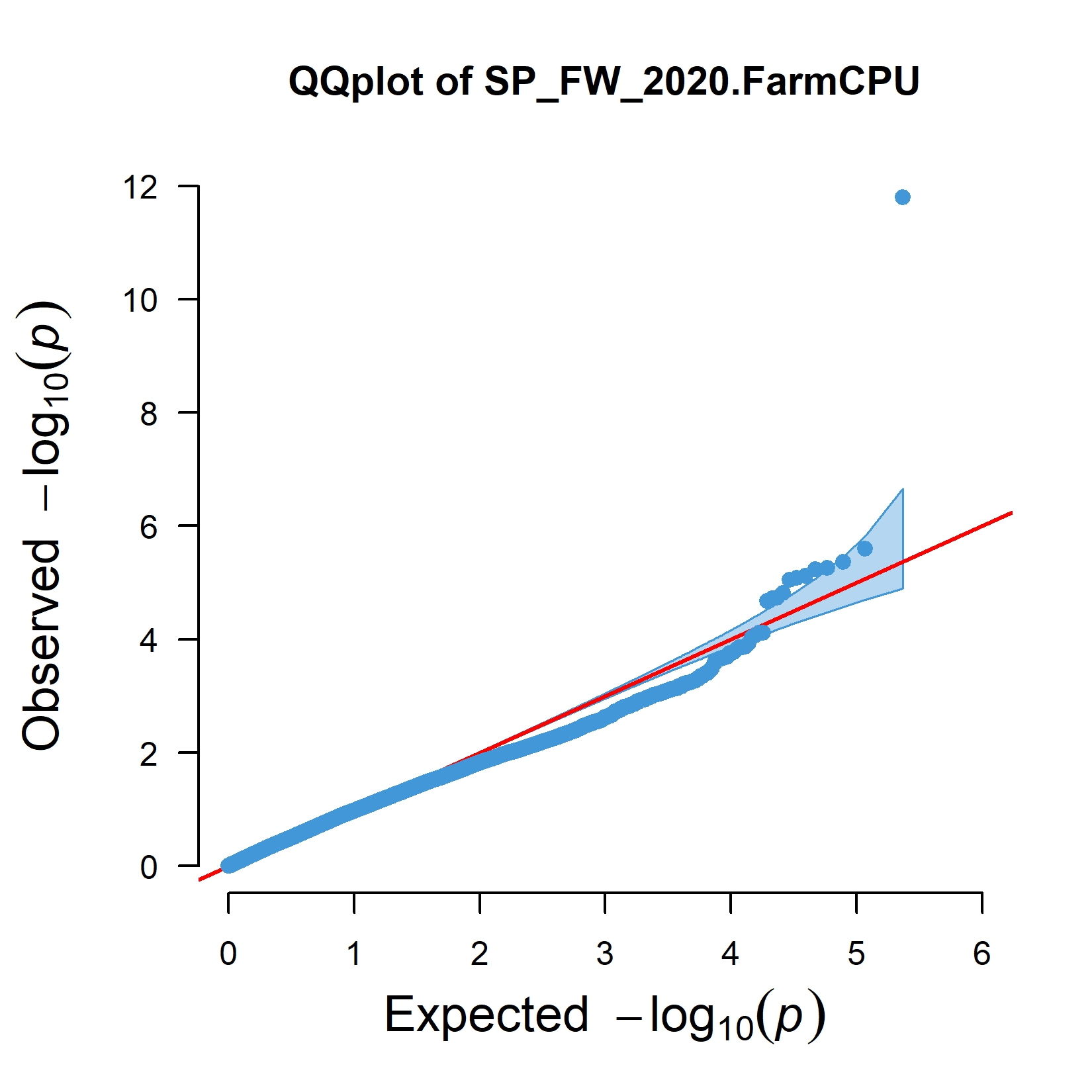 | 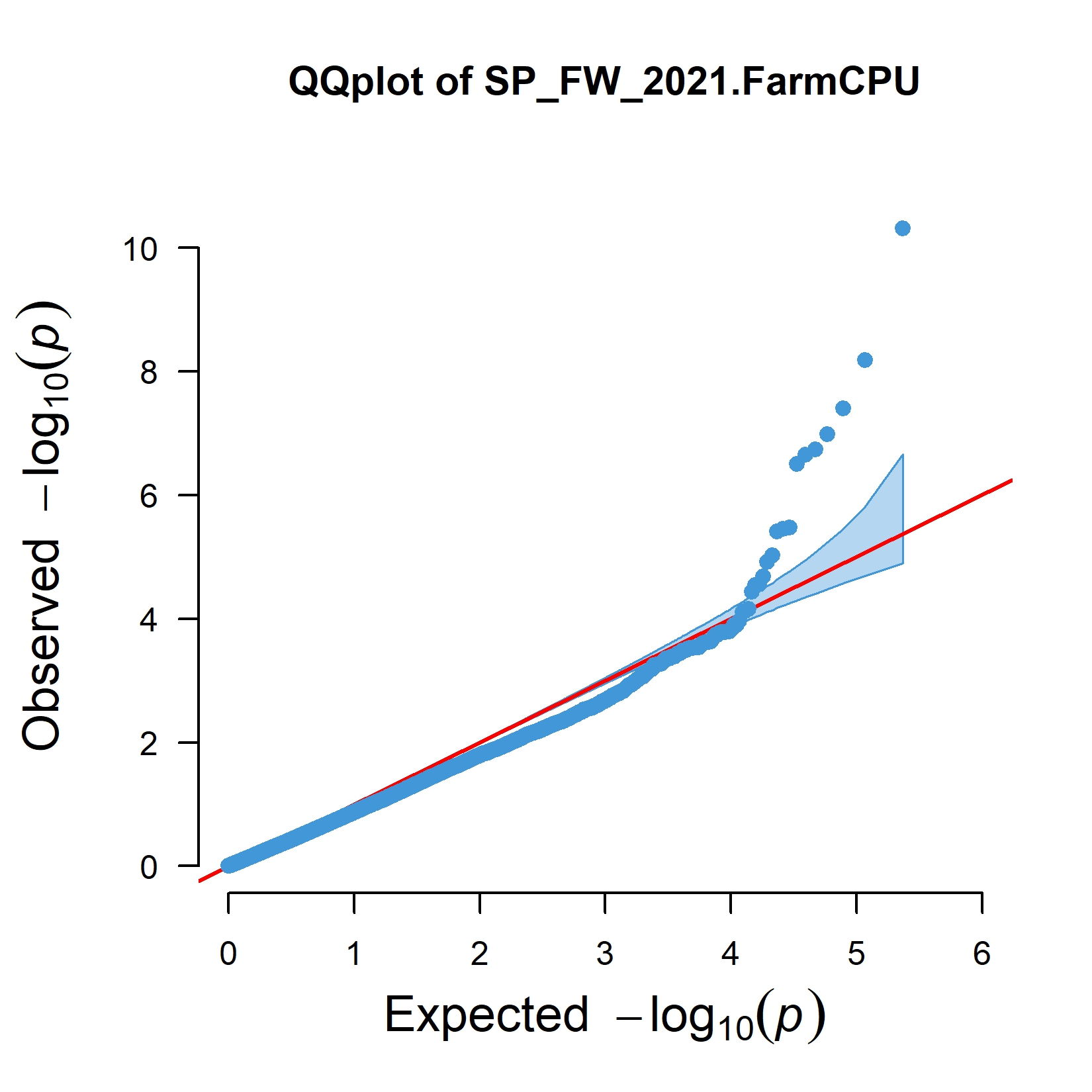 |
| --- | --- | --- |
| 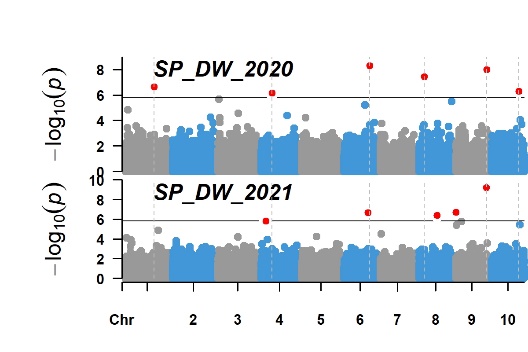 | 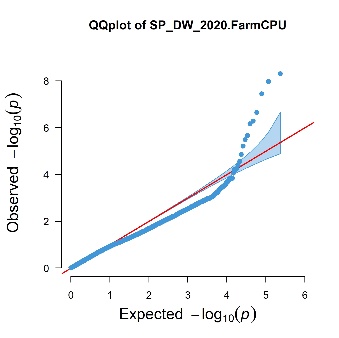 | 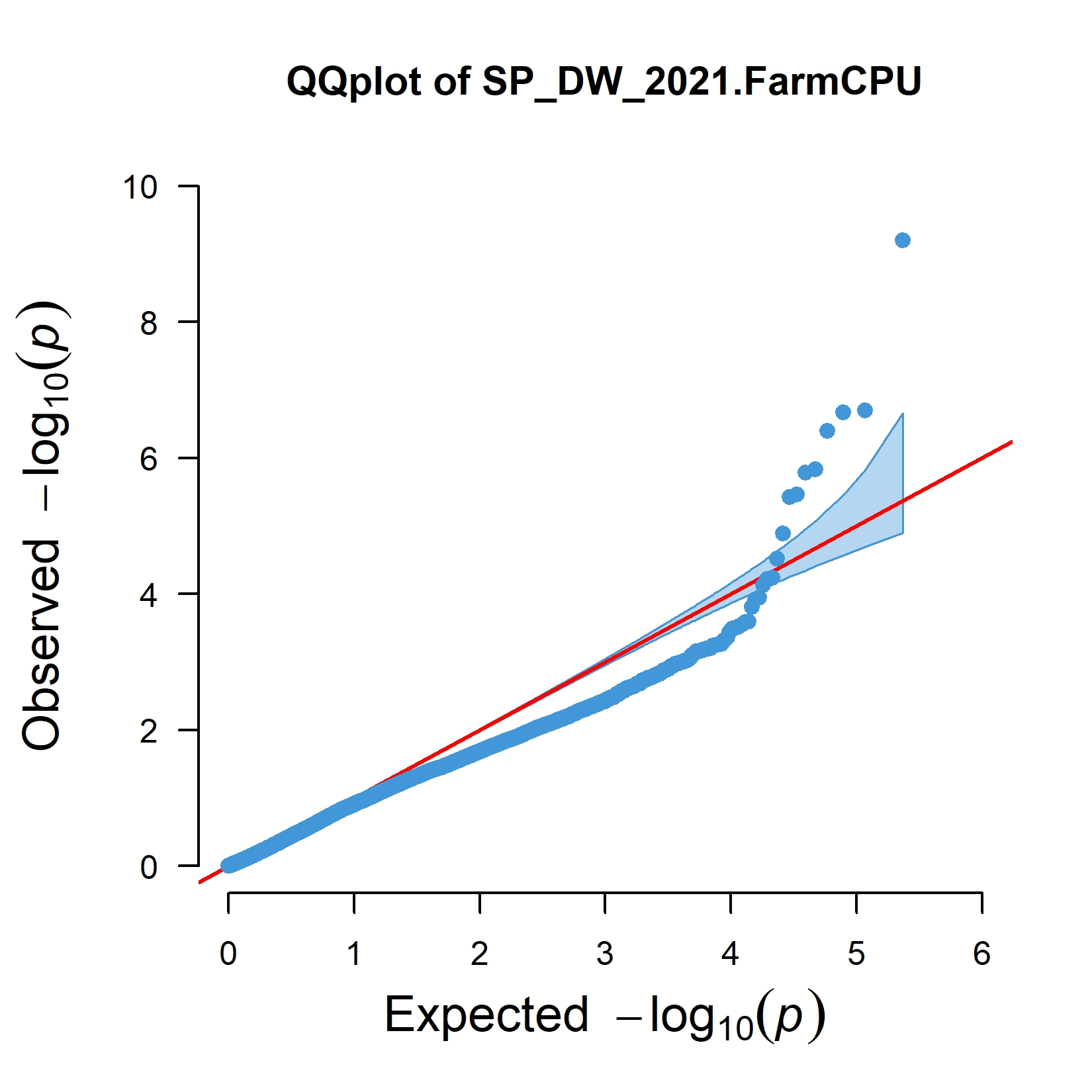 |
| 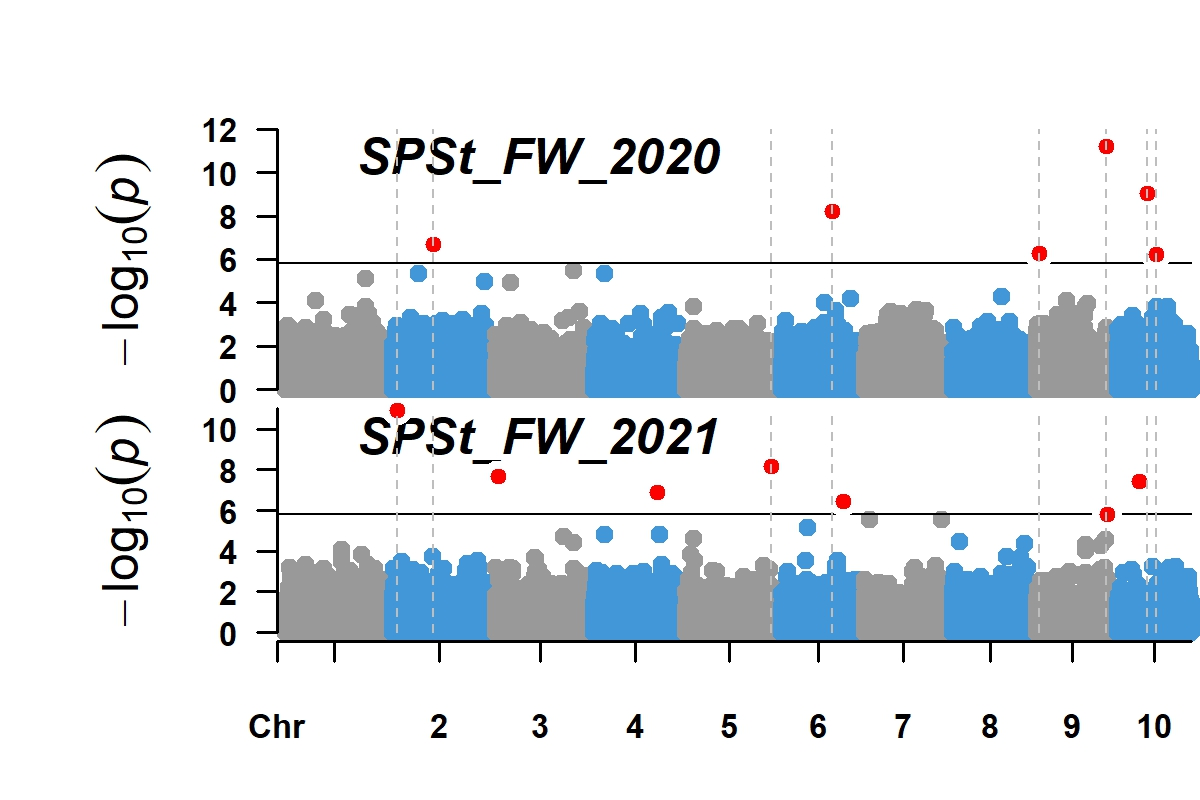 | 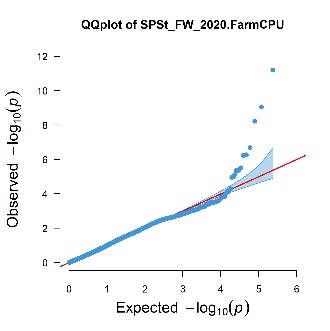 | 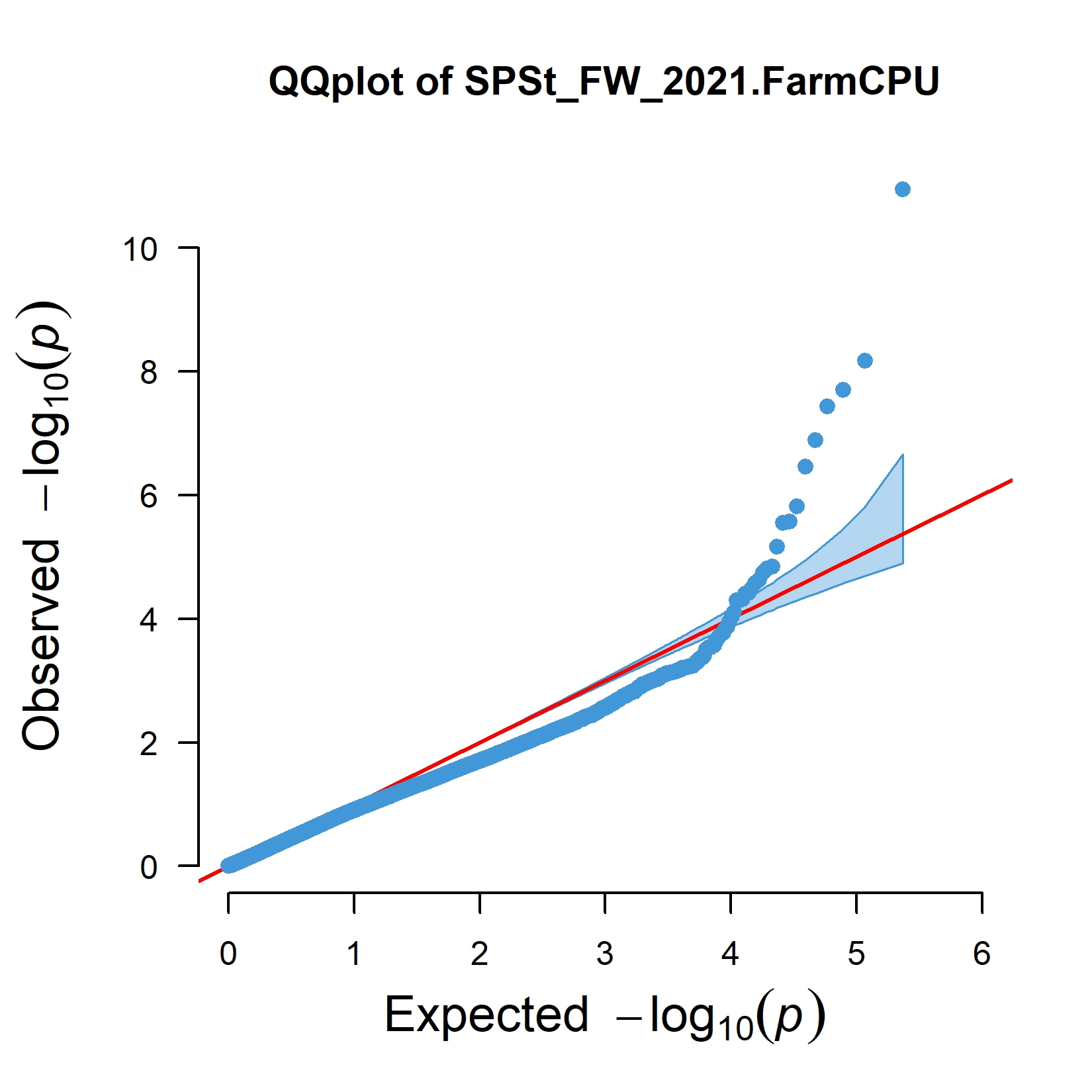 |
| 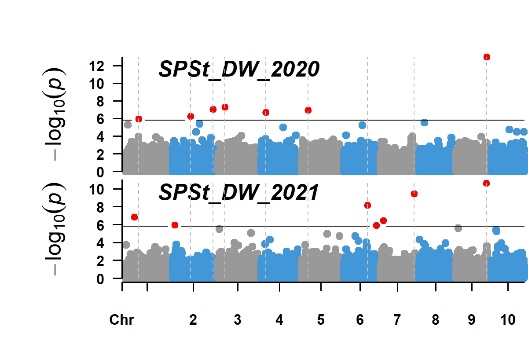 | 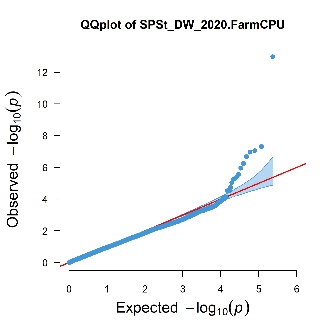 | 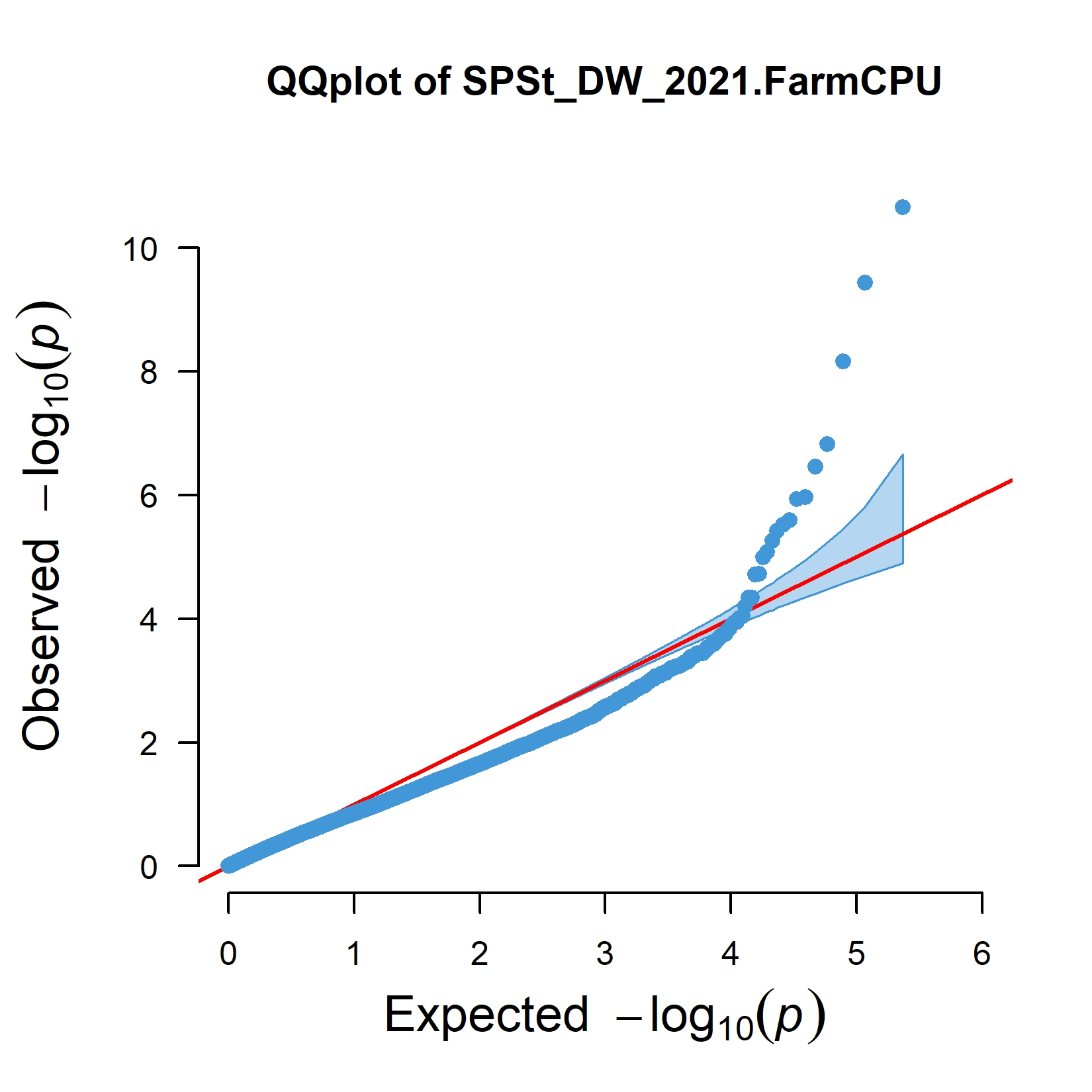 |
| 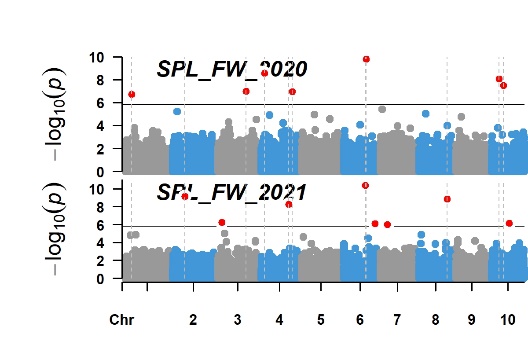 | 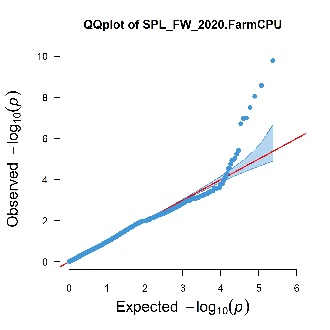 | 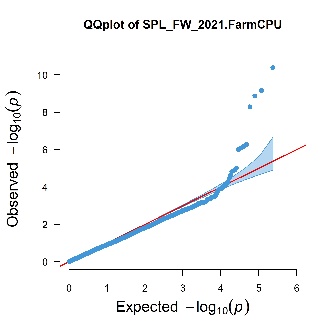 |
| 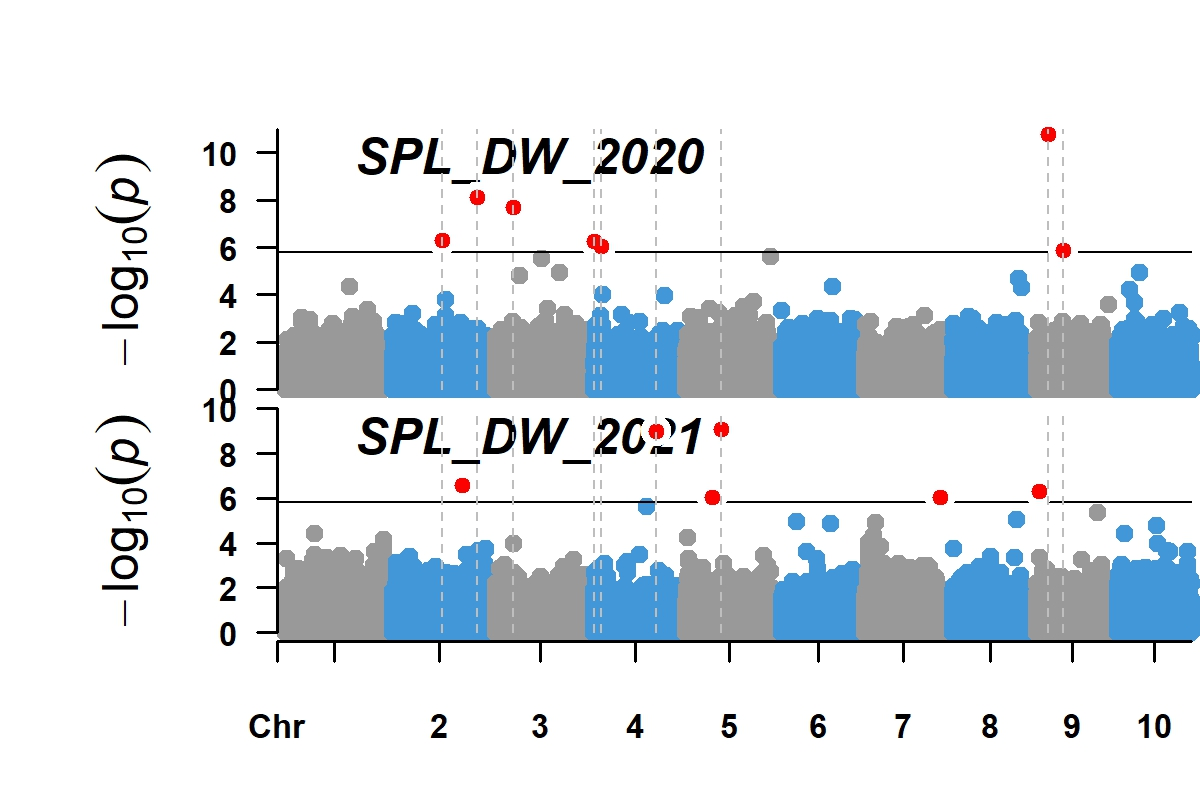 | 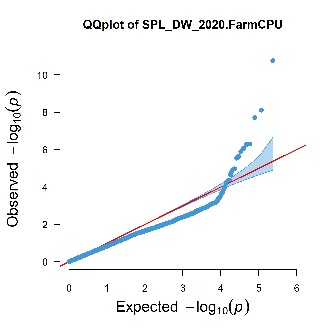 | 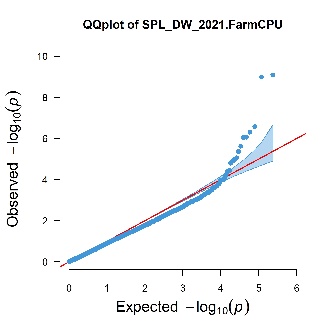 |
| 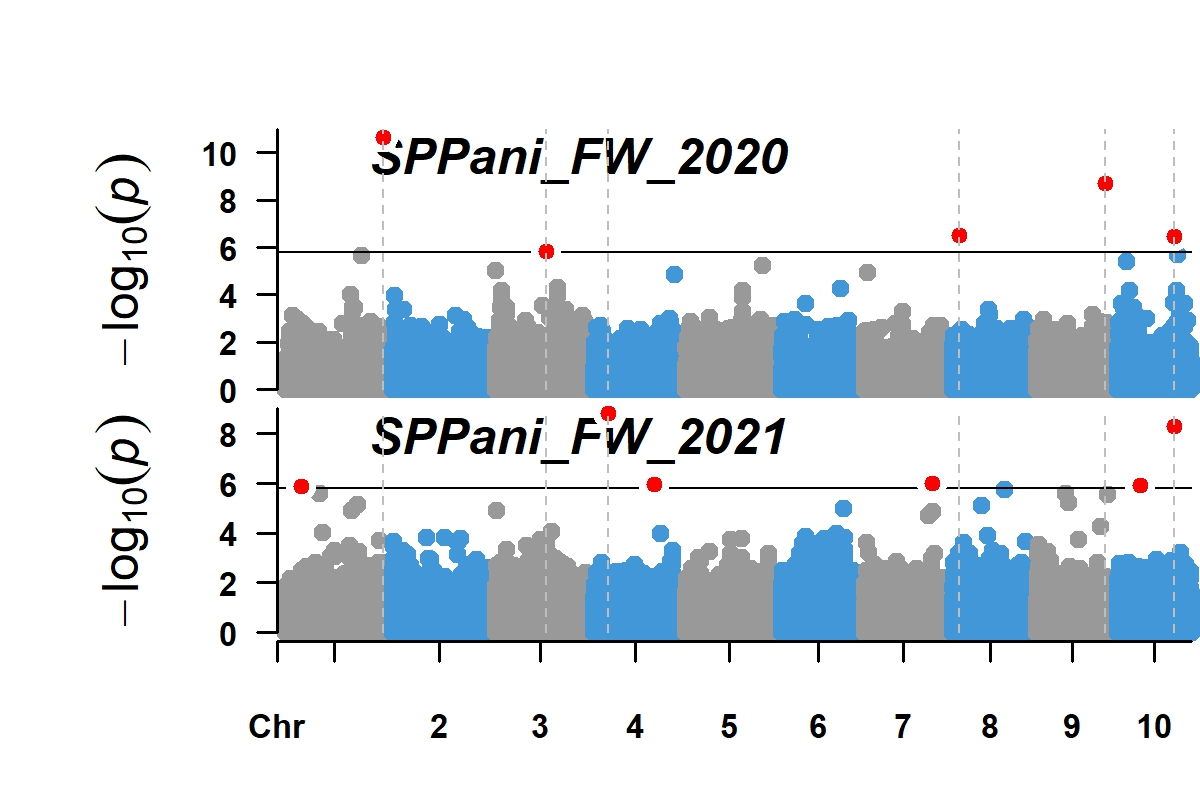 | 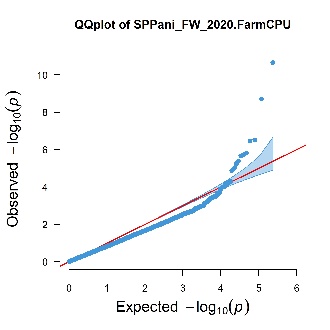 | 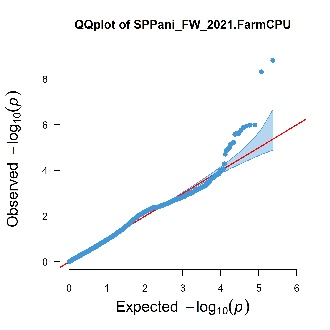 |
| 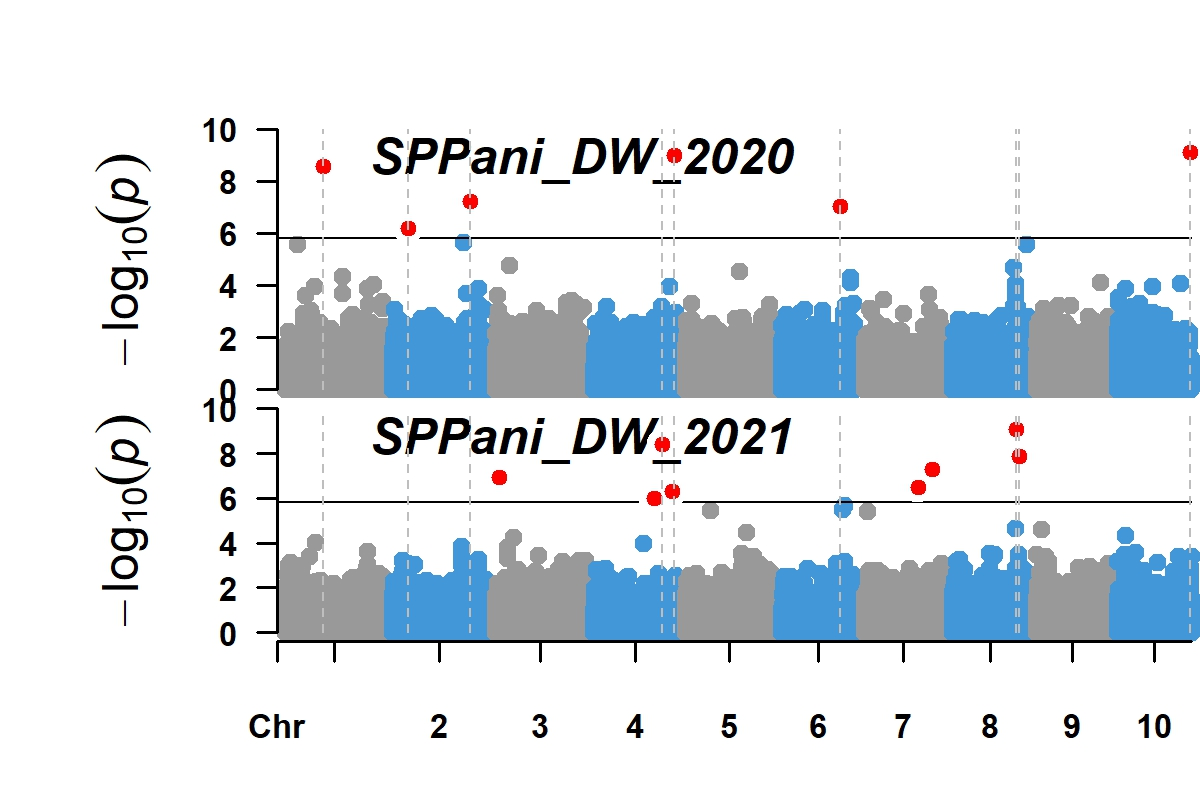 | 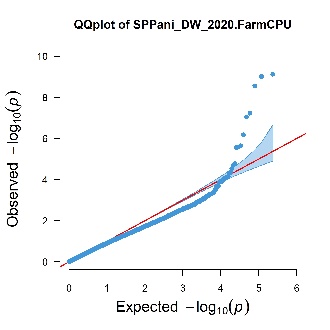 | 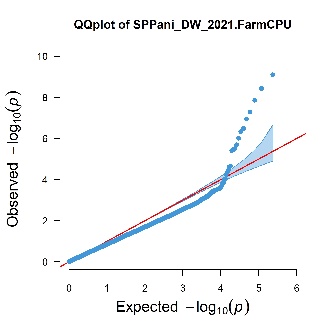 |
| 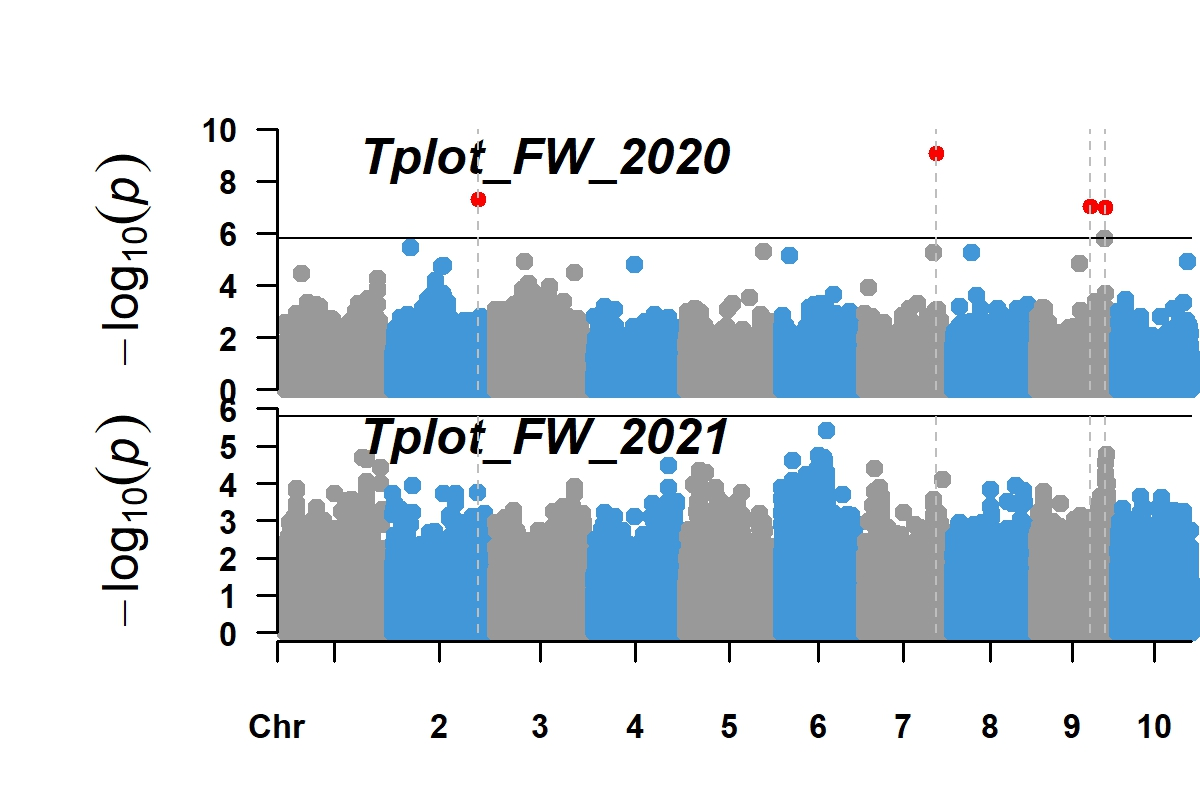 | 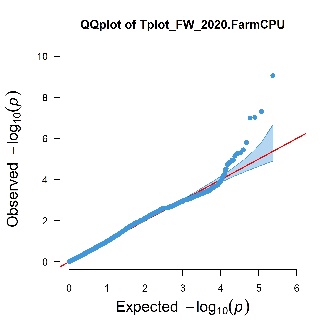 | 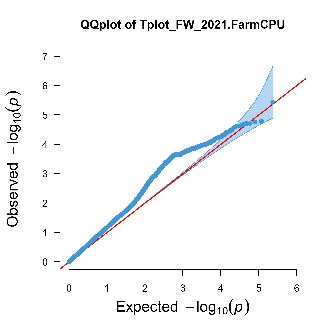 |
| 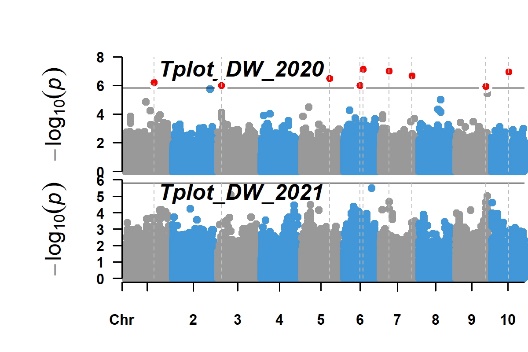 | 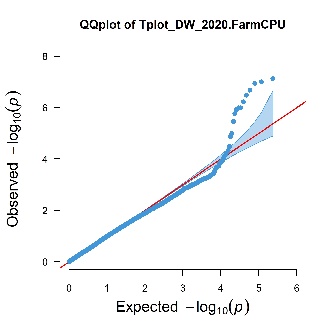 | 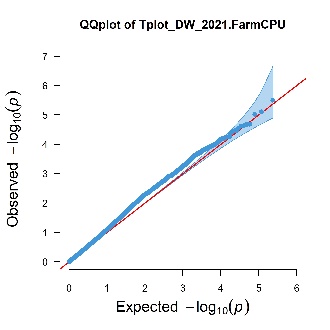 |
| 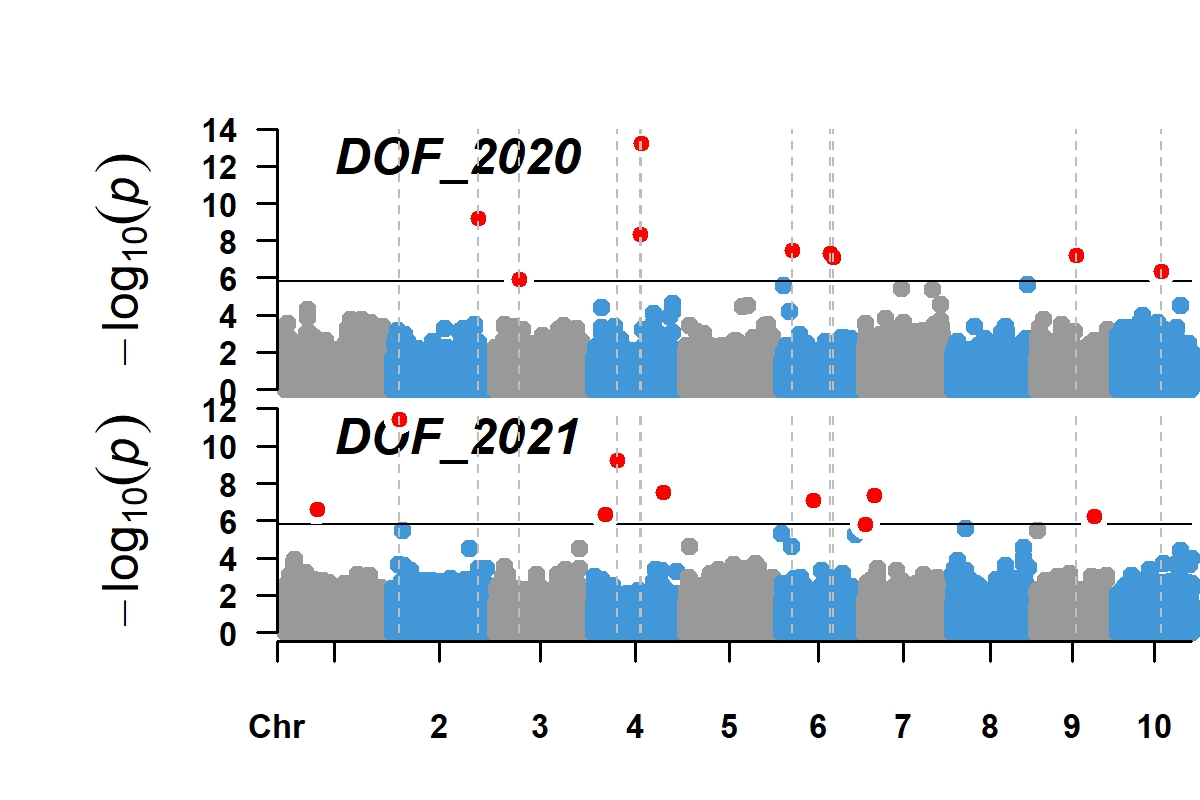 | 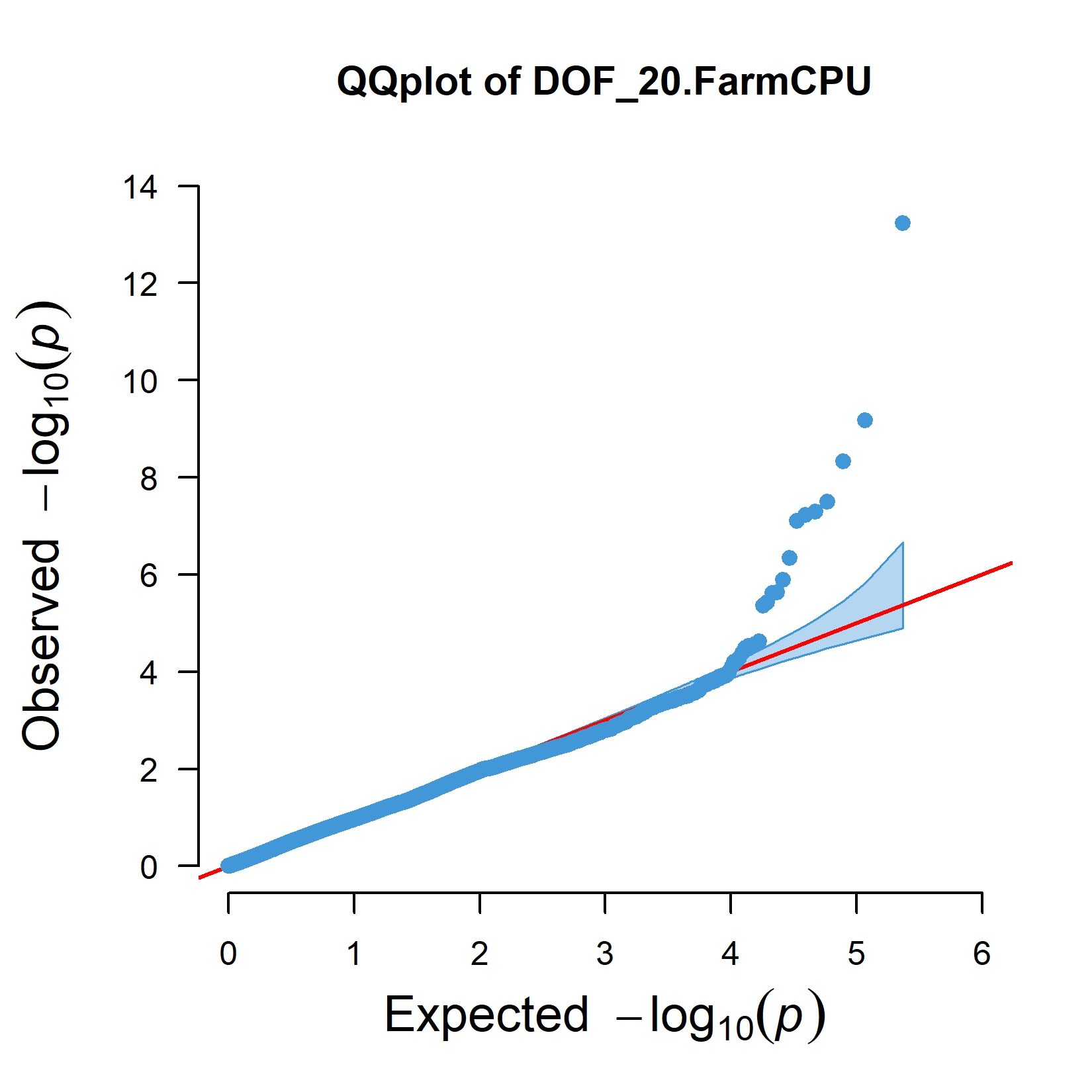 | 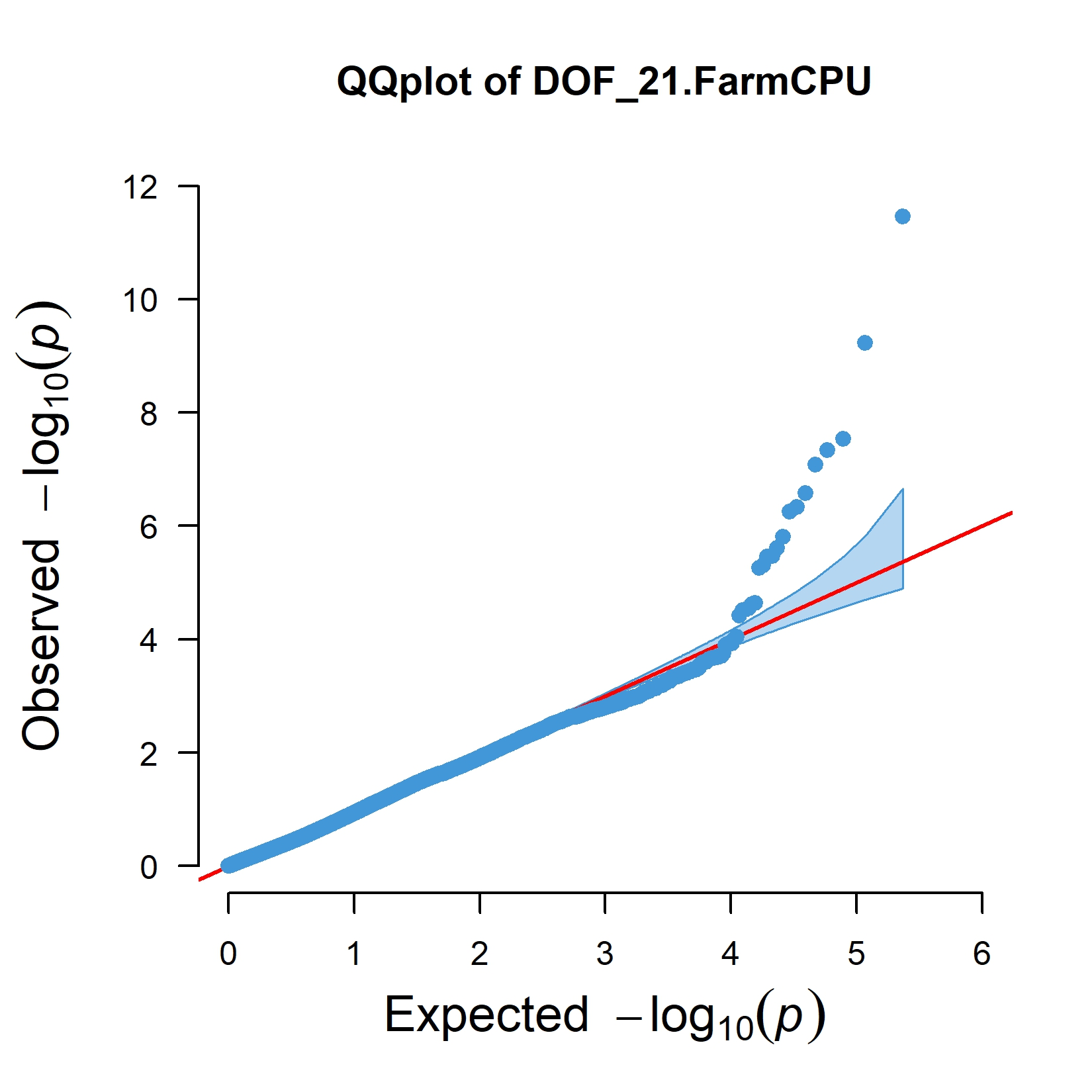 |
| 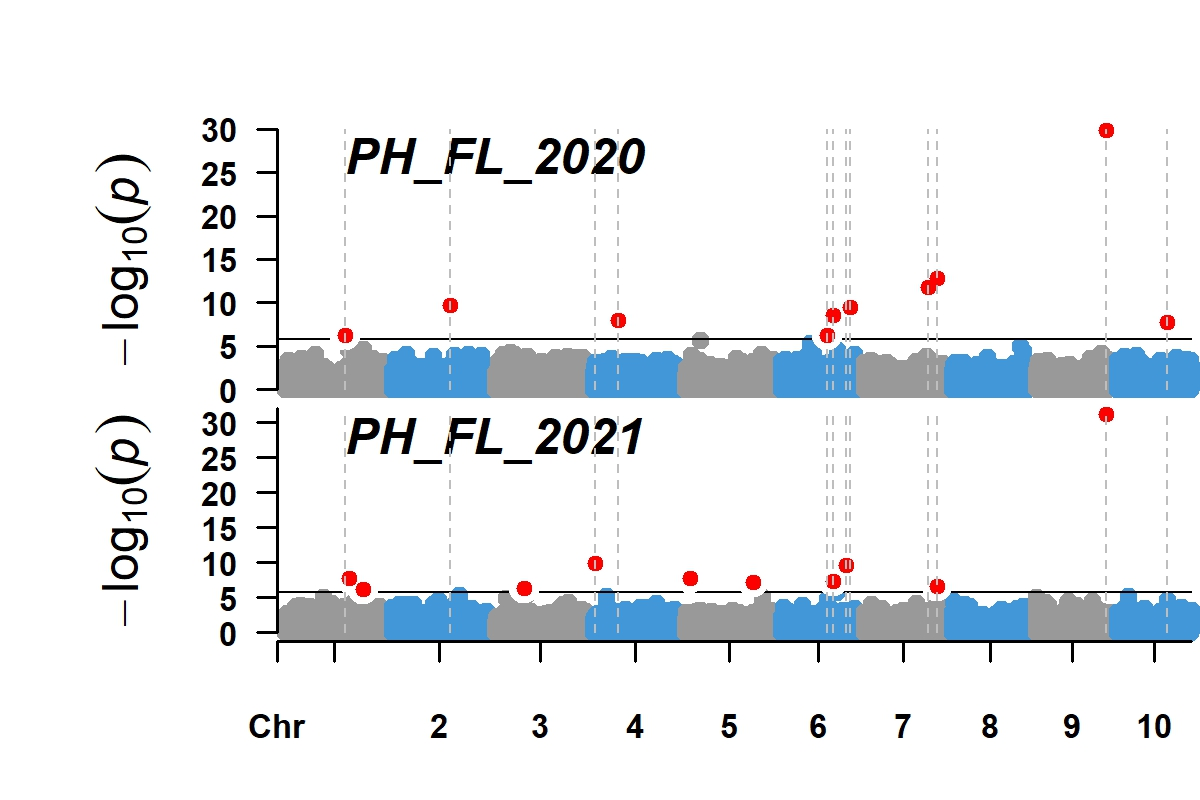 | 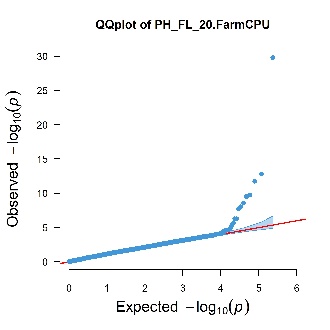 | 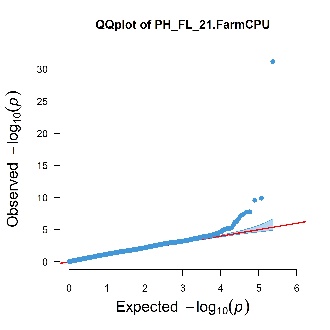 |
| 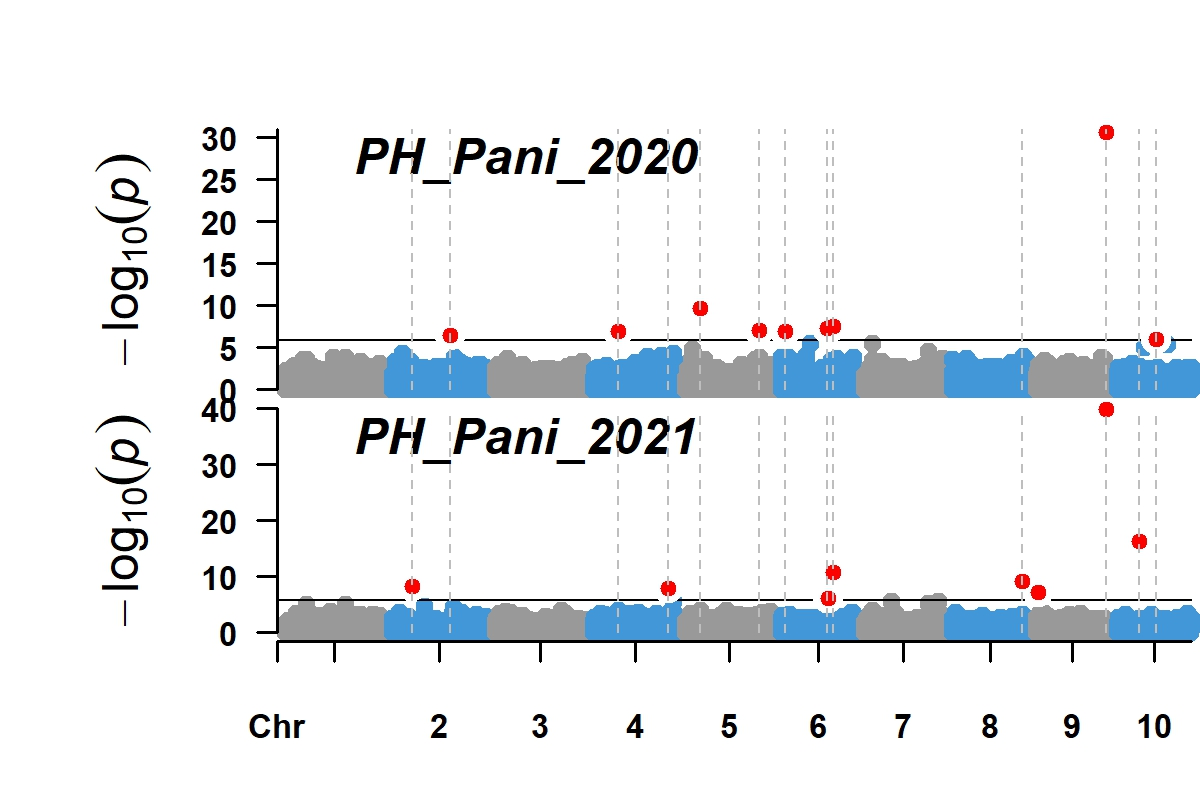 | 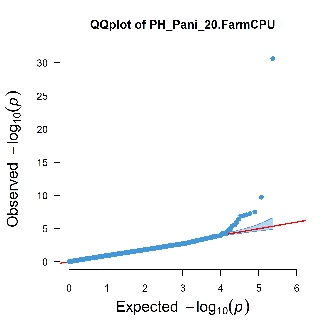 | 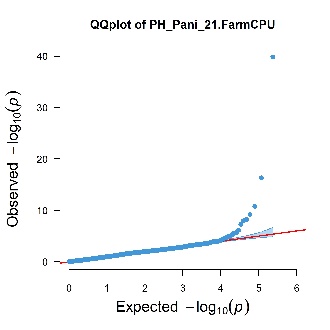 |
| 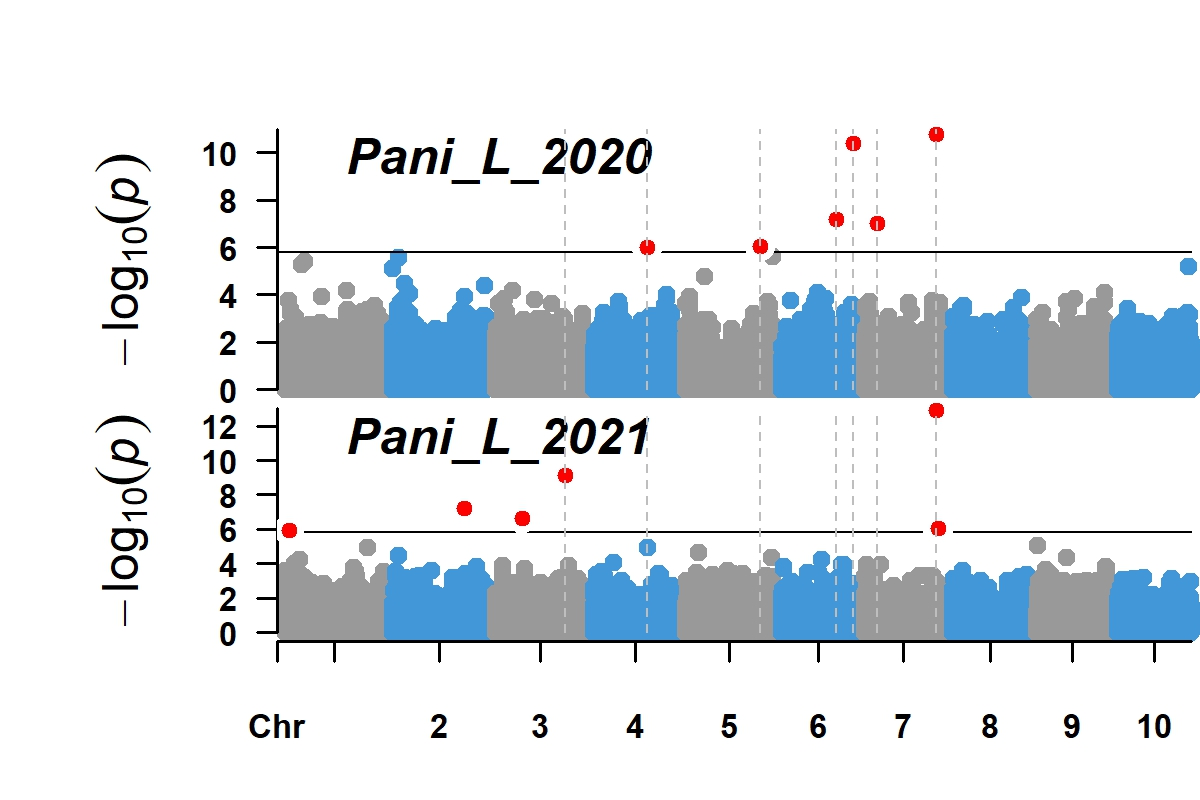 | 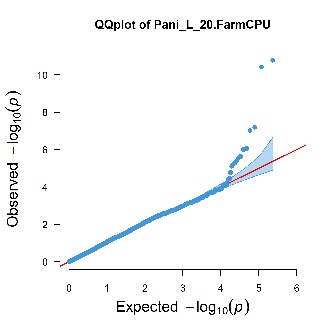 | 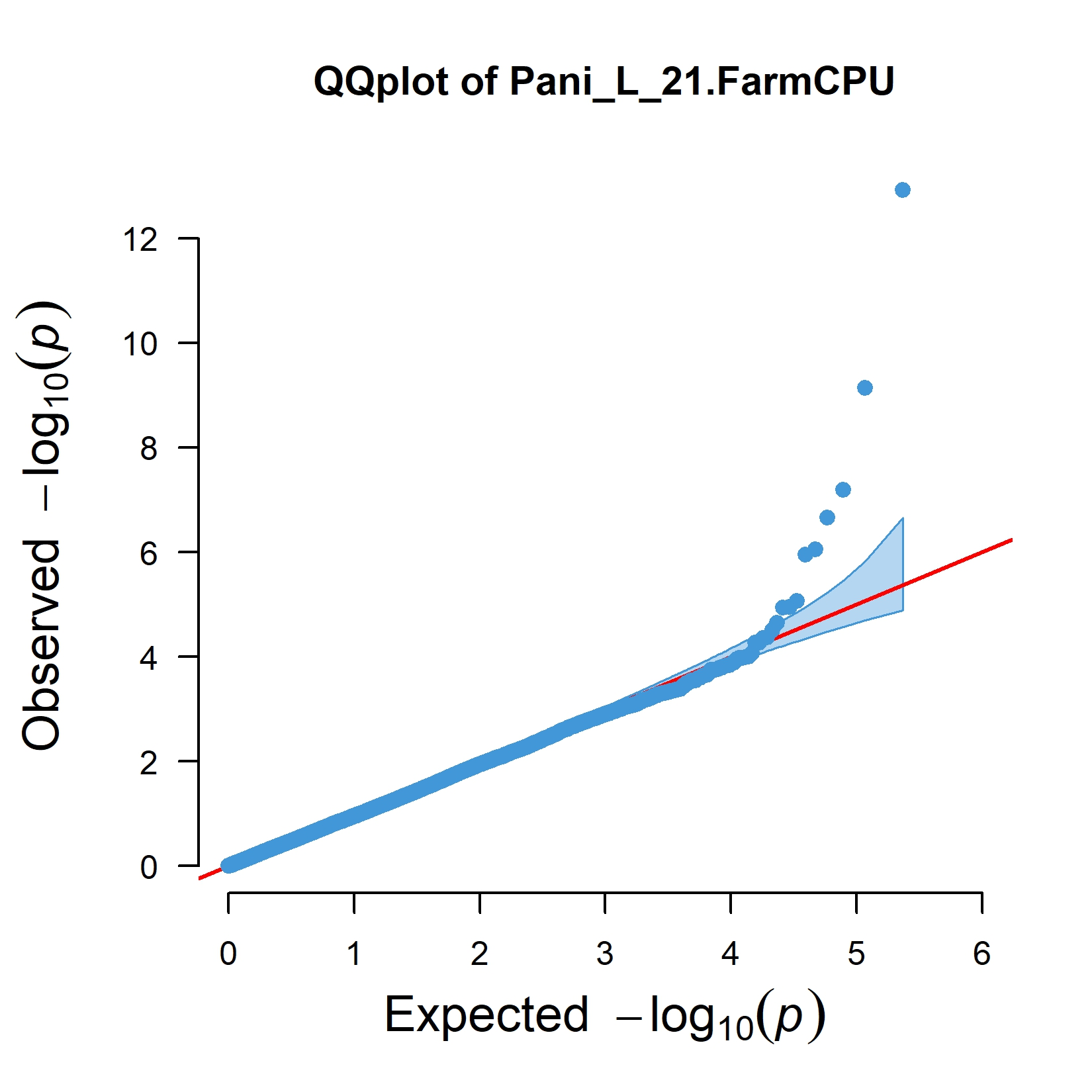 |
| 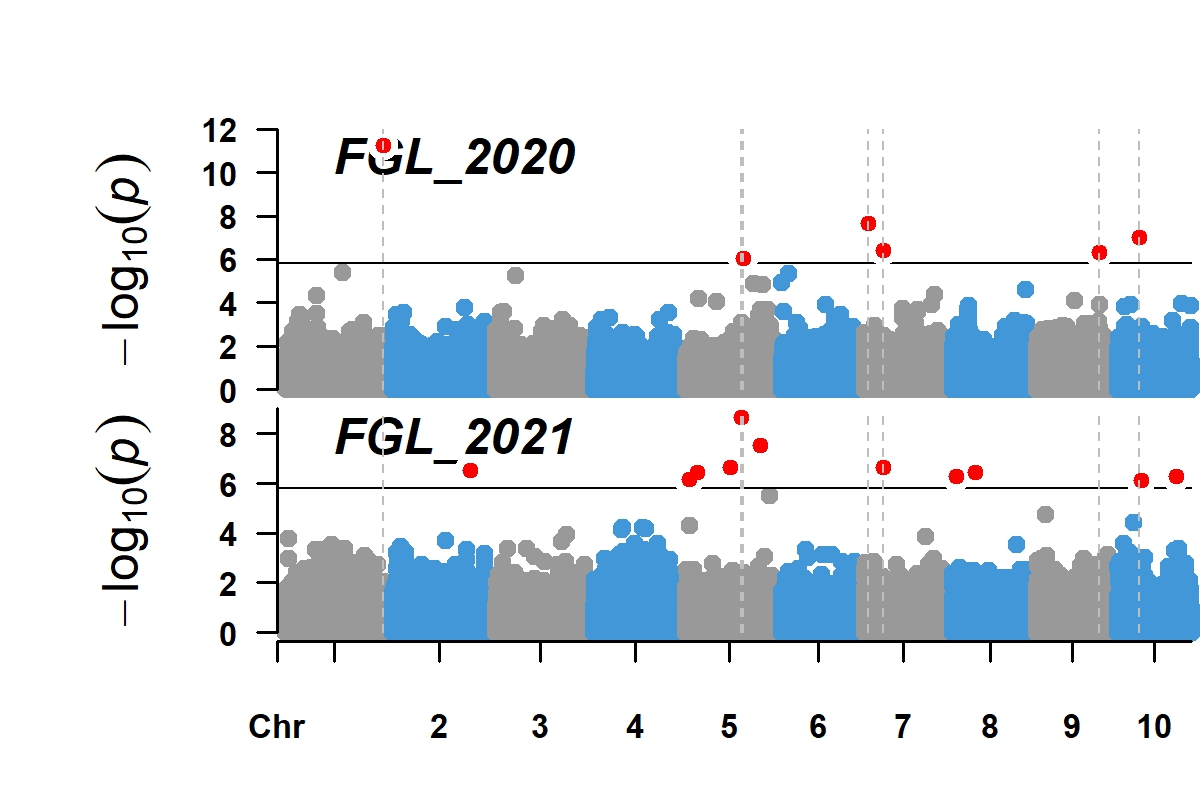 | 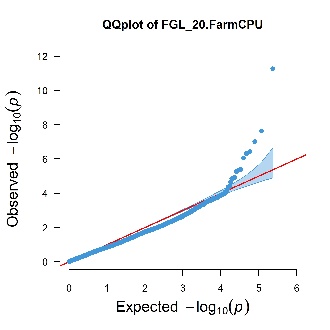 | 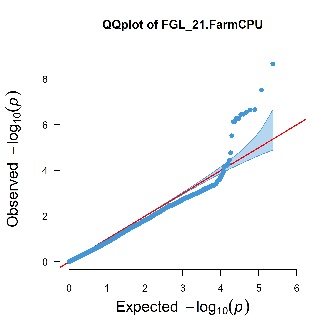 |
| 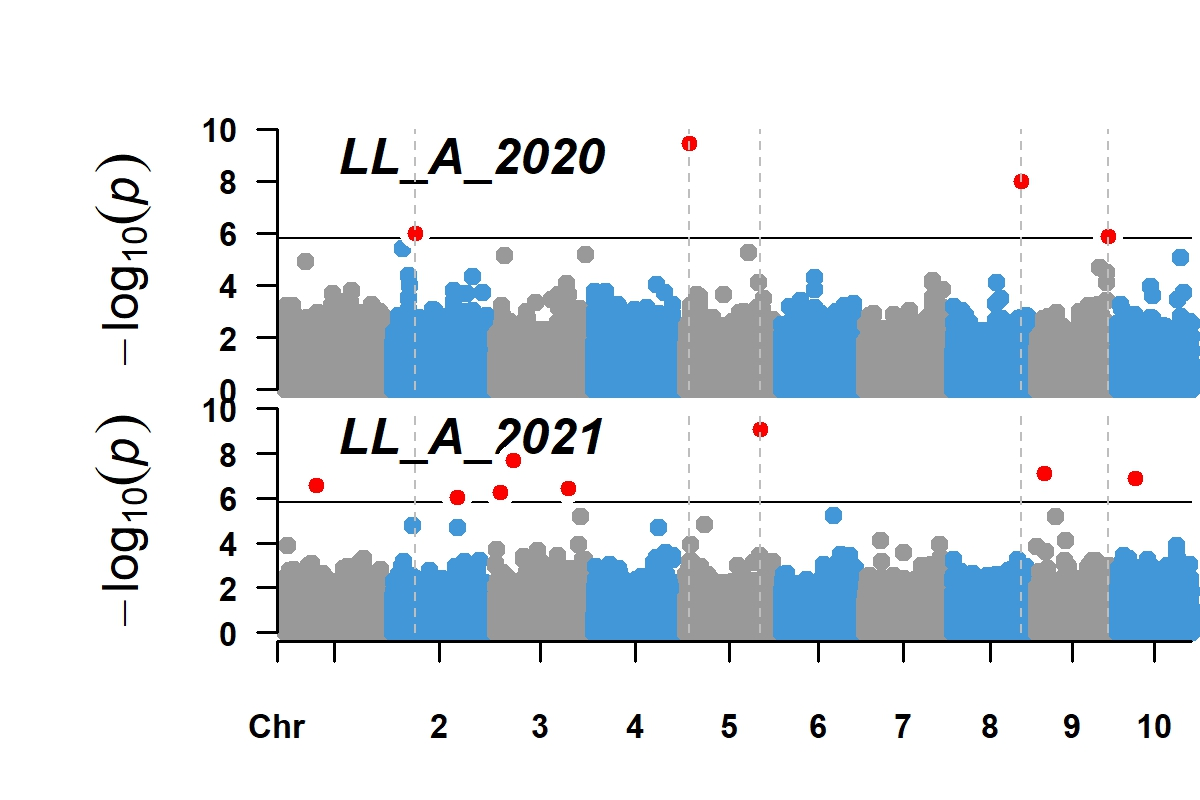 | 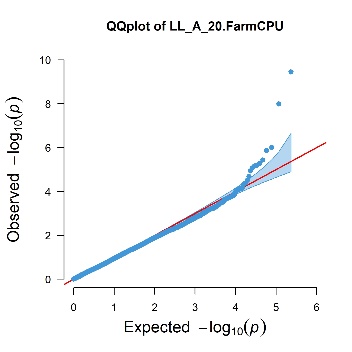 | 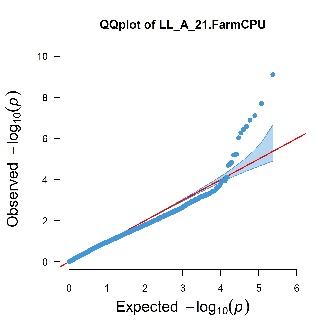 |
| 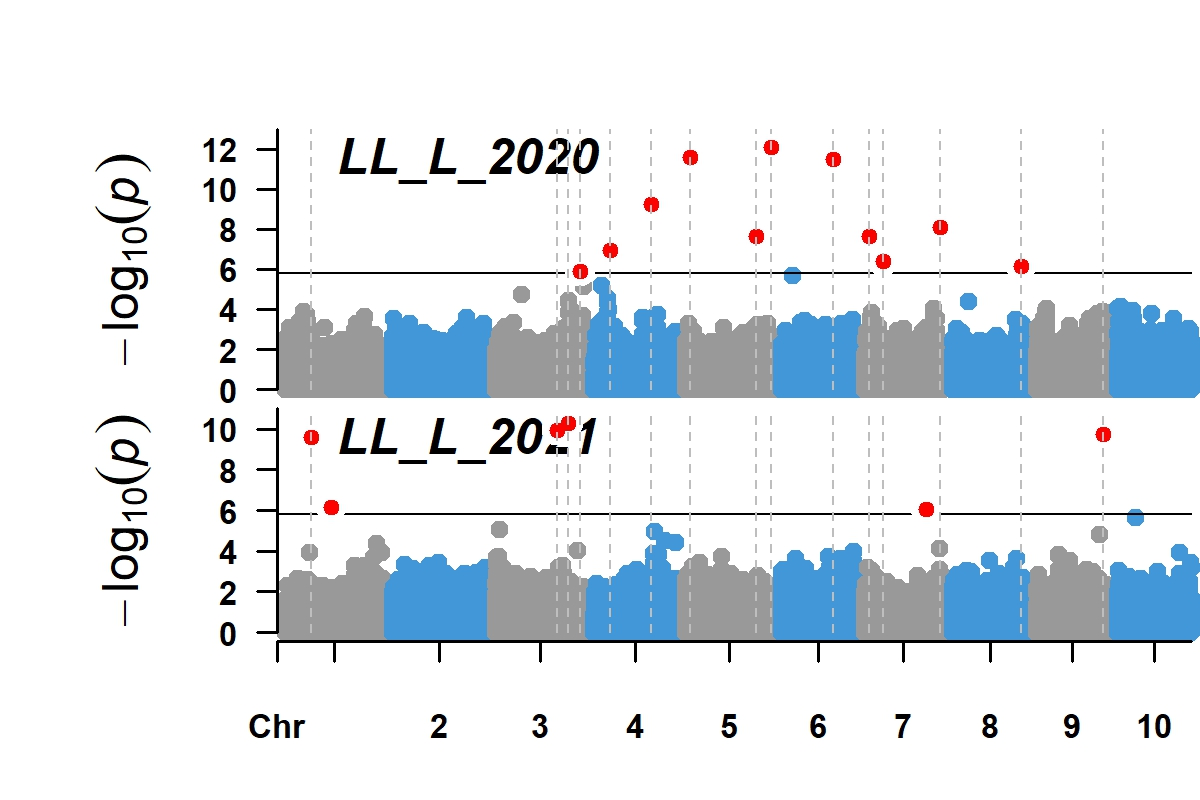 | 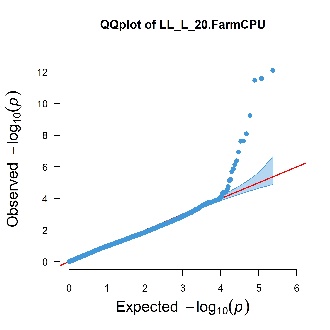 | 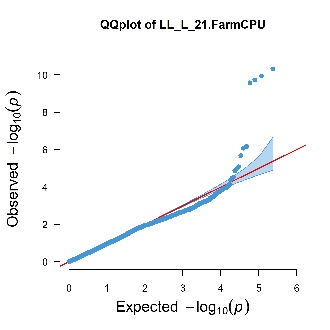 |
| 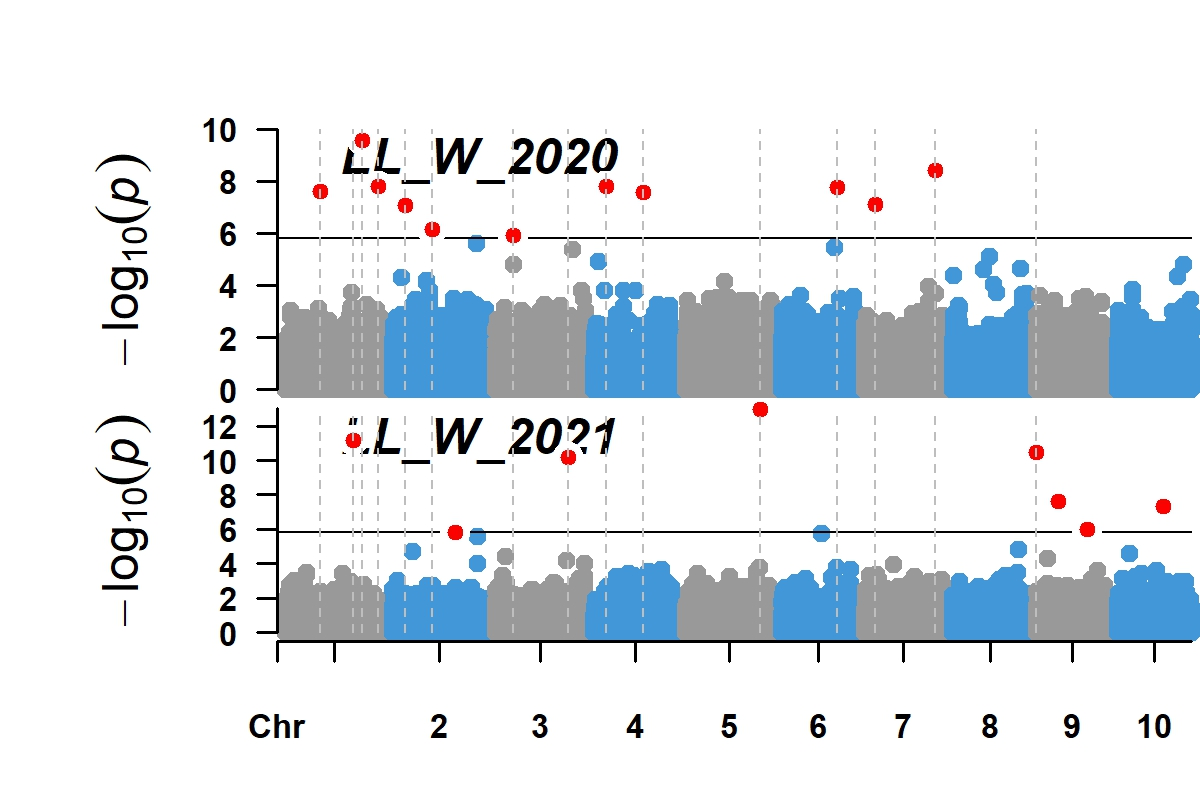 | 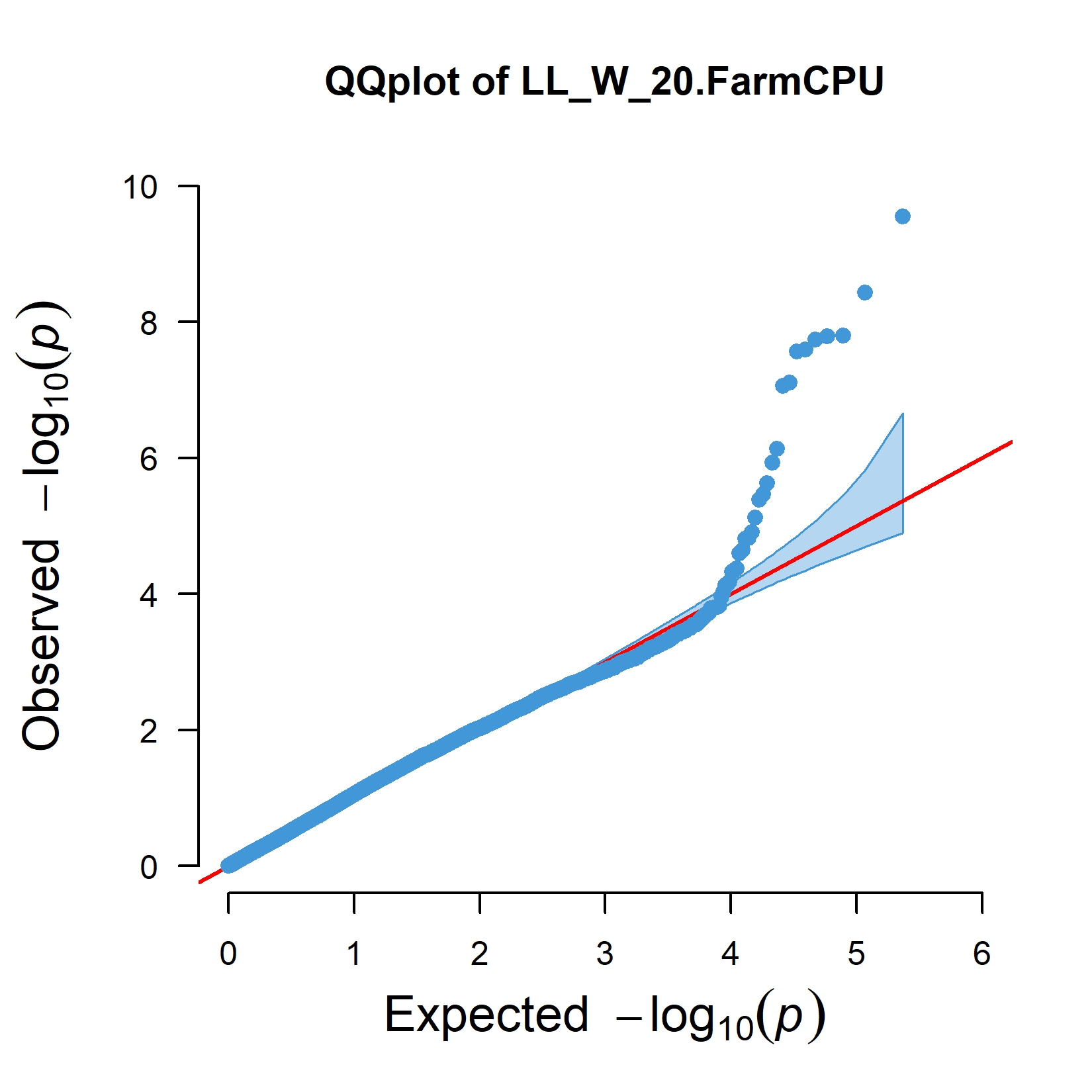 | 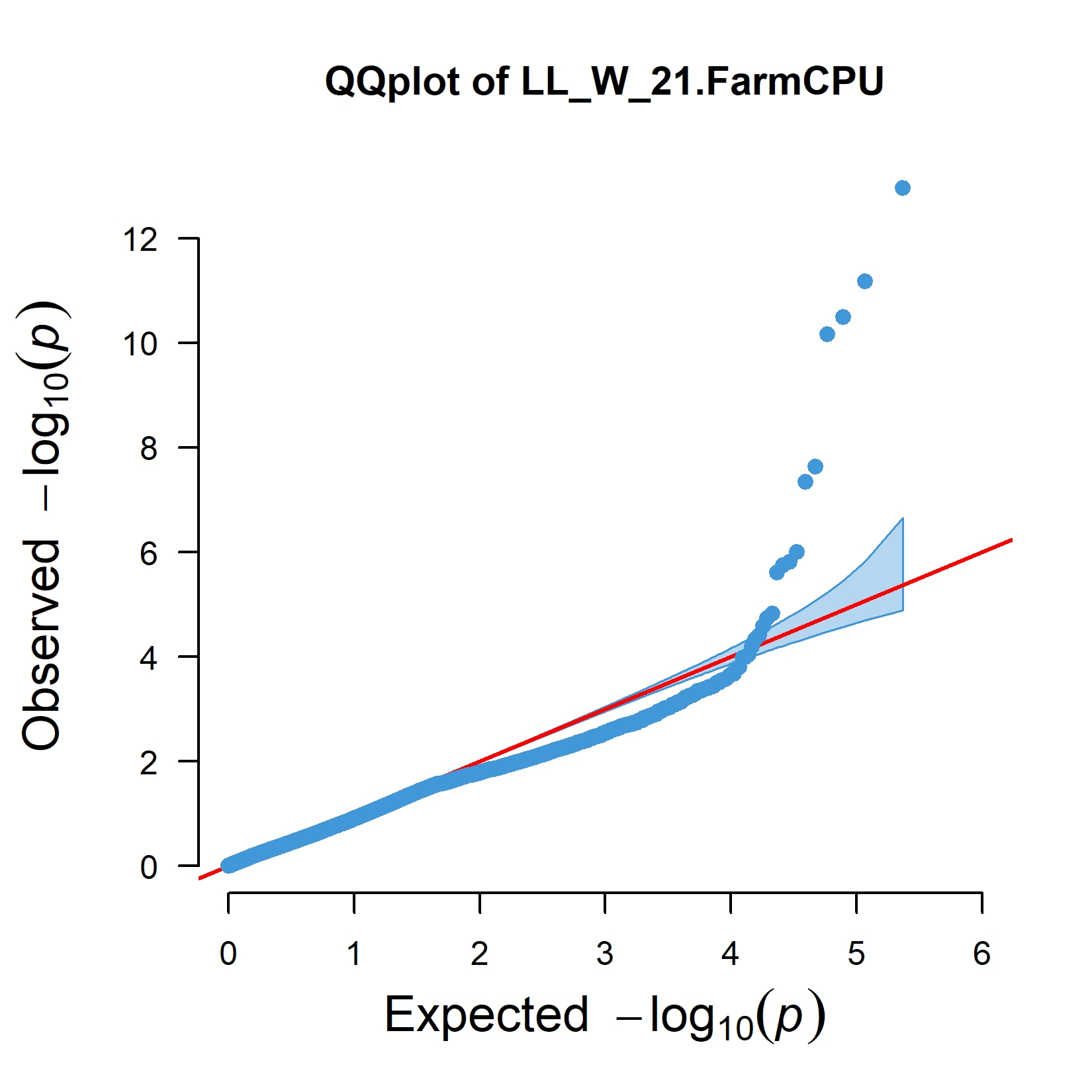 |
| 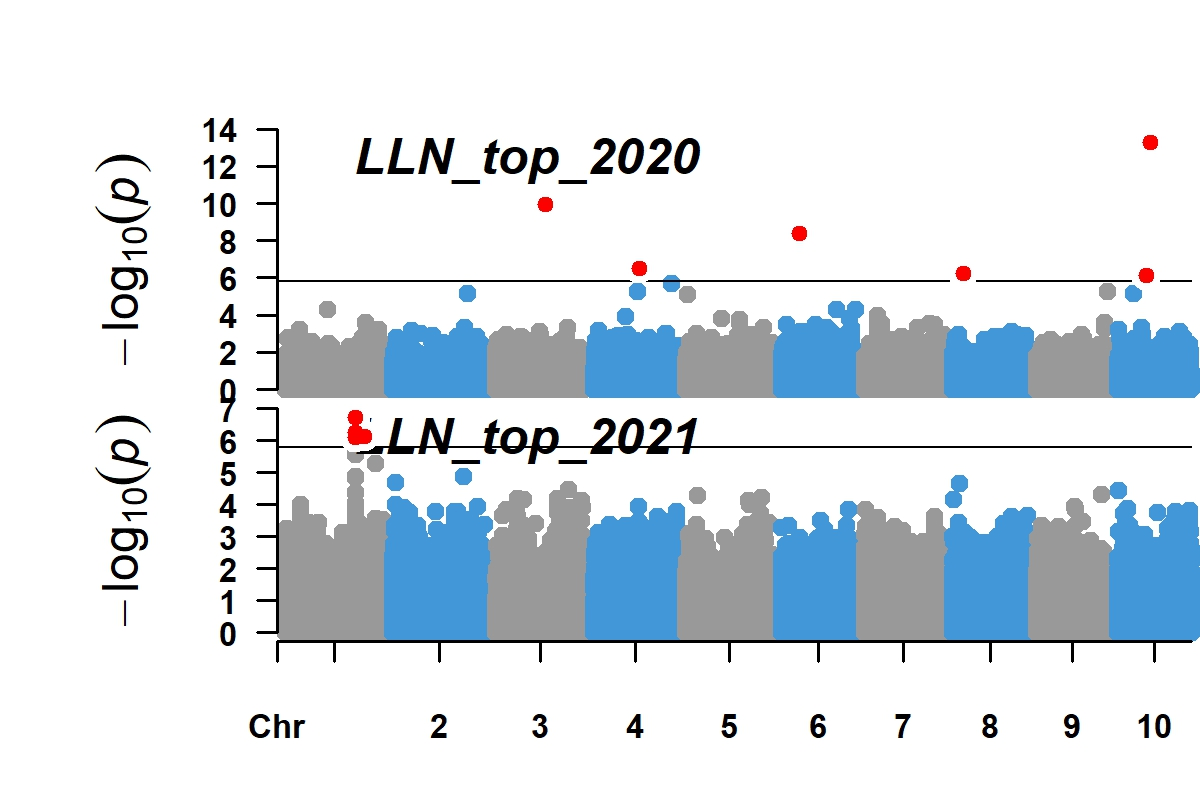 | 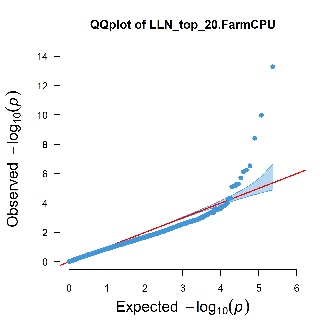 | 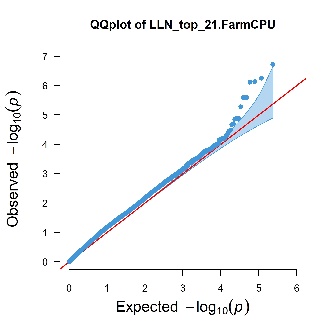 |
| 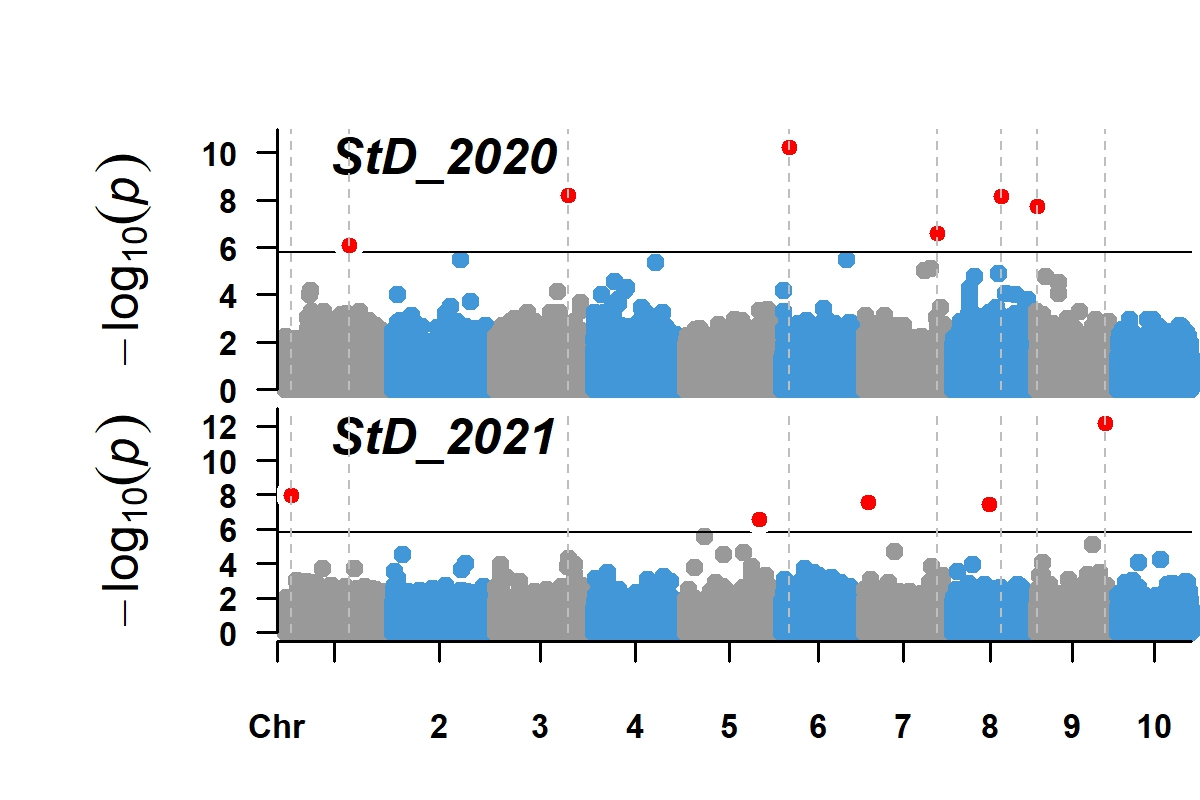 | 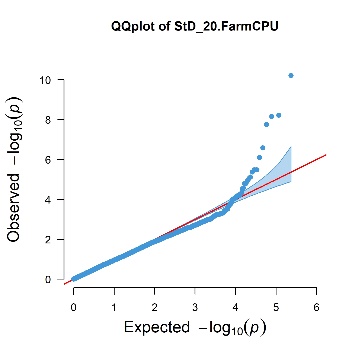 | 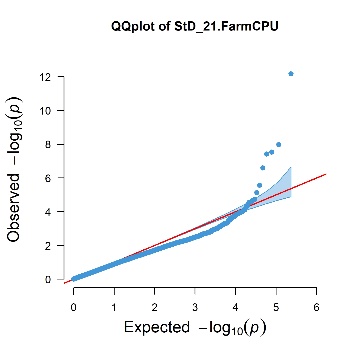 |
| 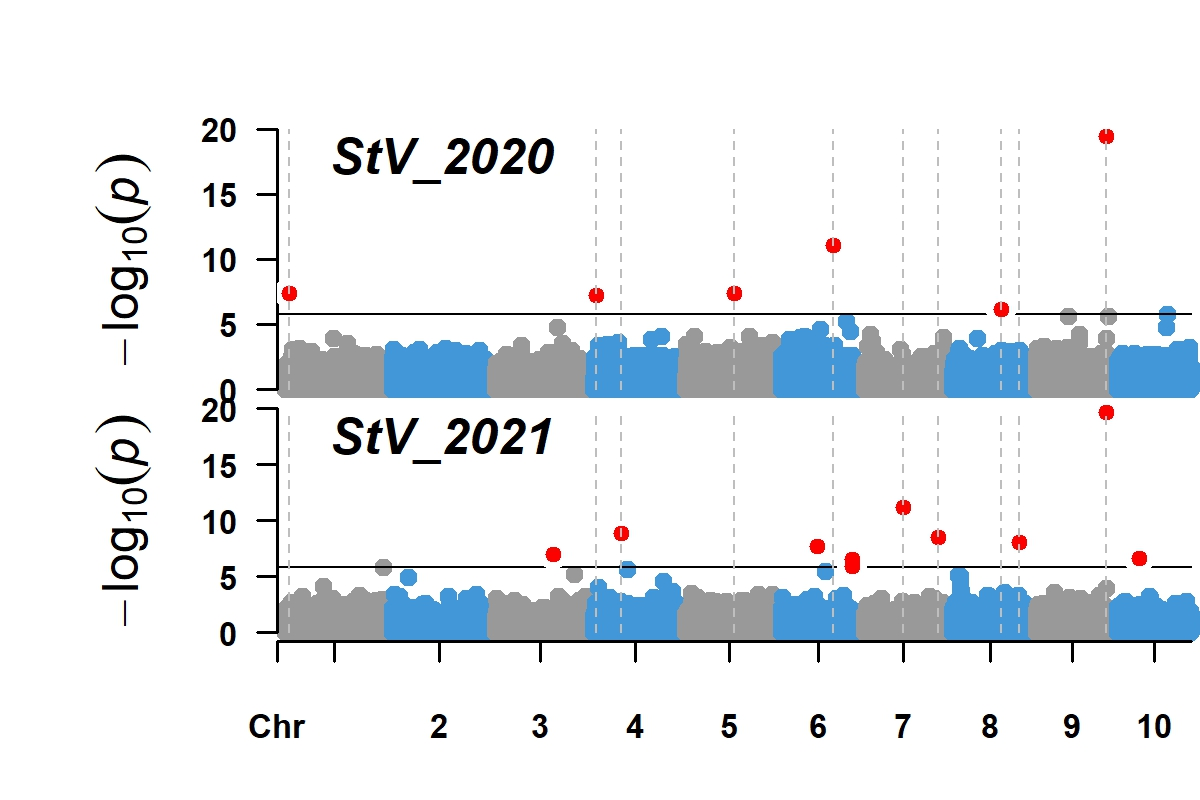 | 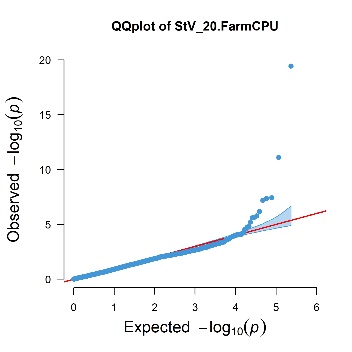 | 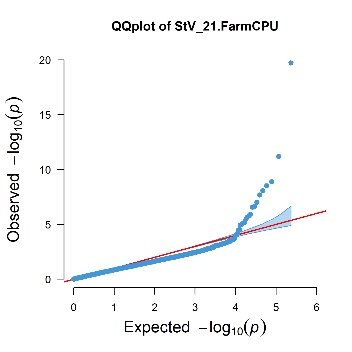 |
| 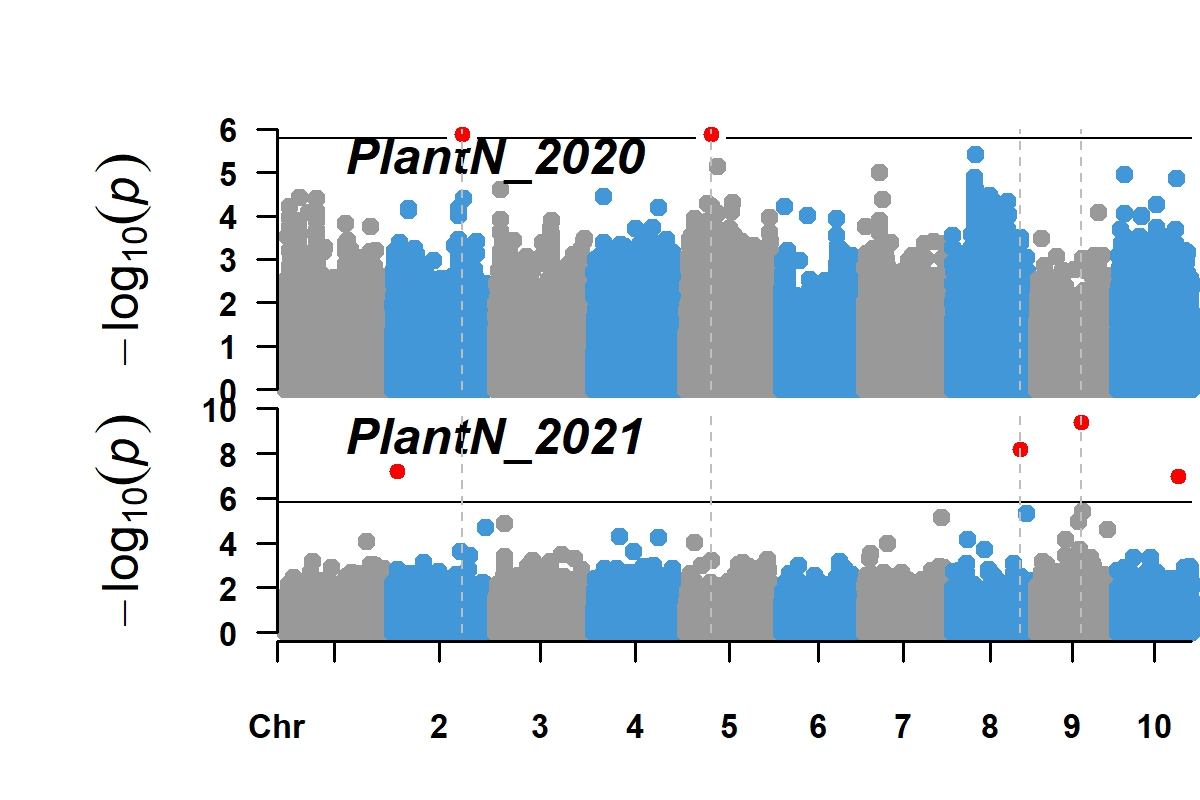 | 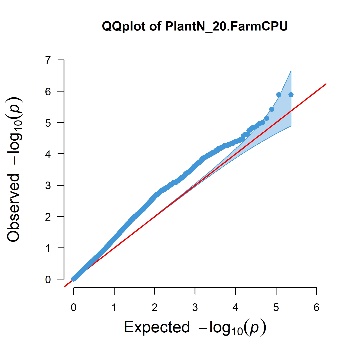 | 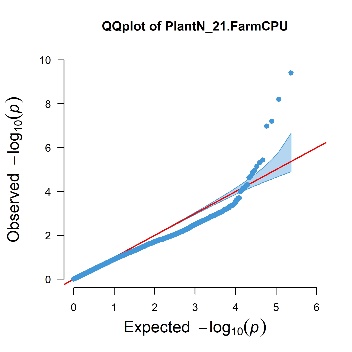 |
| 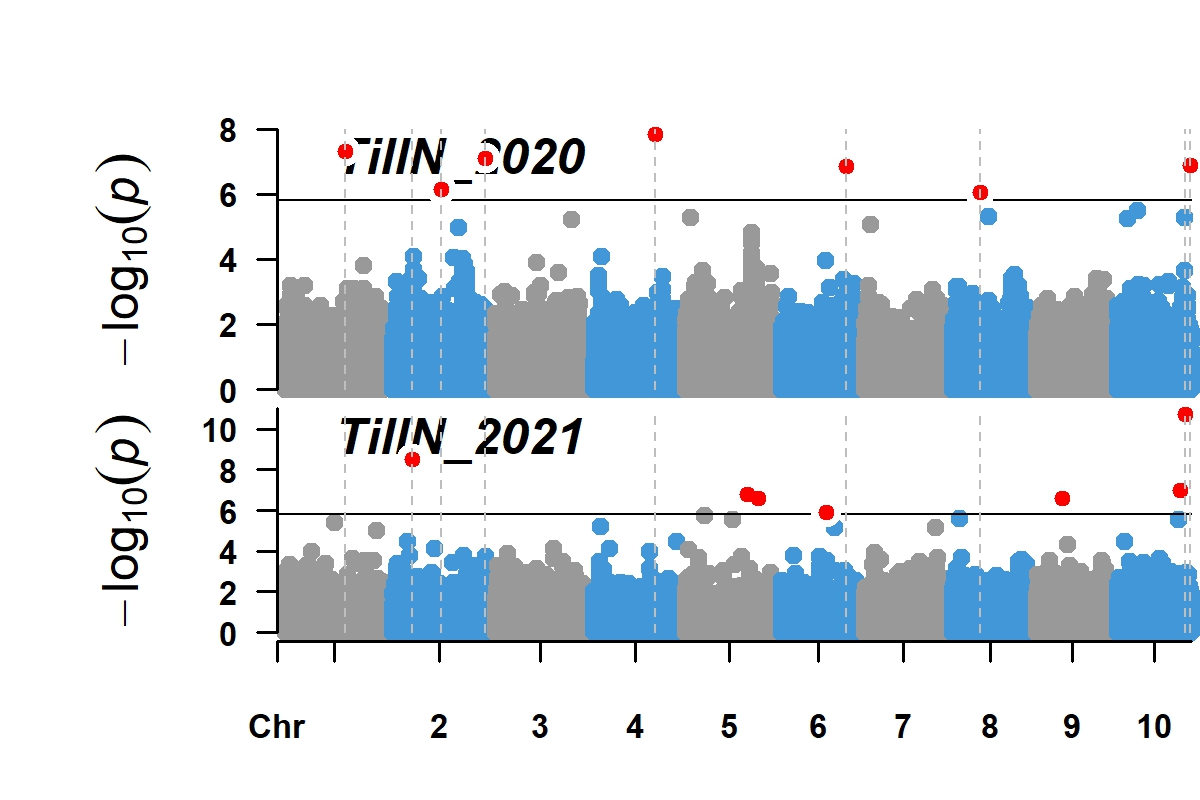 | 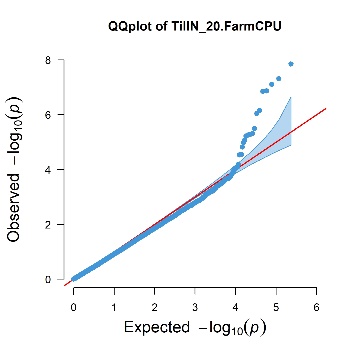 | 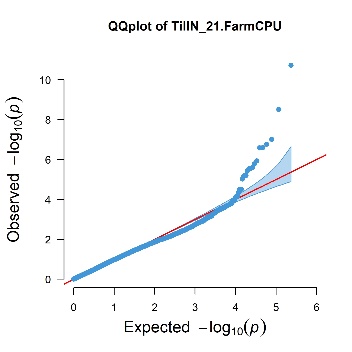 |
| 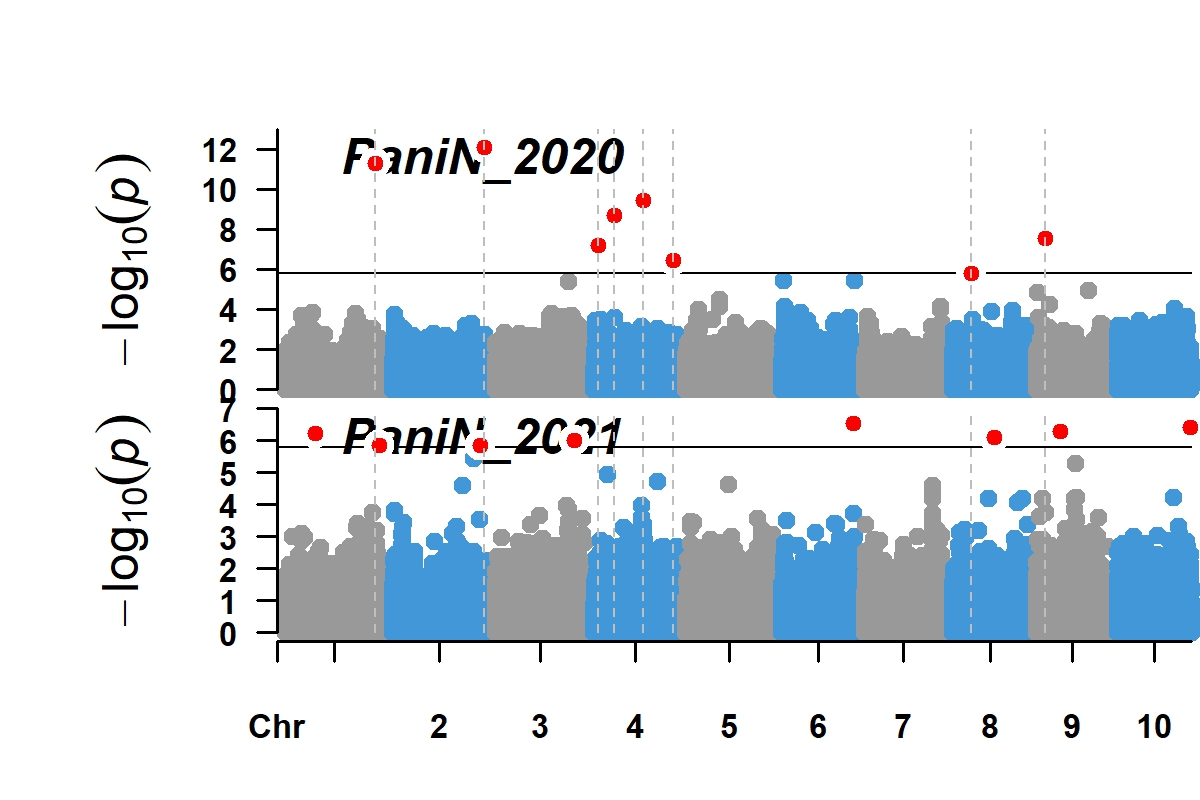 | 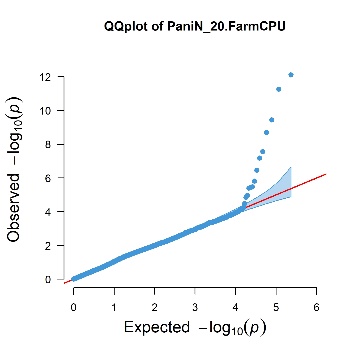 | 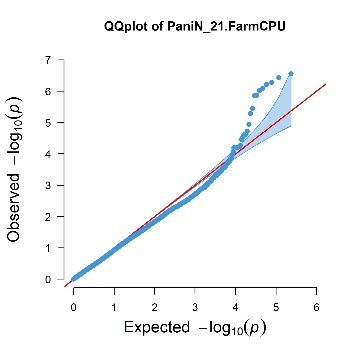 |

**Figure S3. Genome-wide association study (GWAS) for multiple architectural and biomass yield traits using sorghum association panel (403 accessions) over two growing seasons.** Significant SNPs were visualized through Manhattan plot (left) and the quantile-quantile (Q-Q) plots (right). The horizontal grey line in the Manhattan plot indicates a significant threshold (P ≤ 2.4 × 10^-6^).

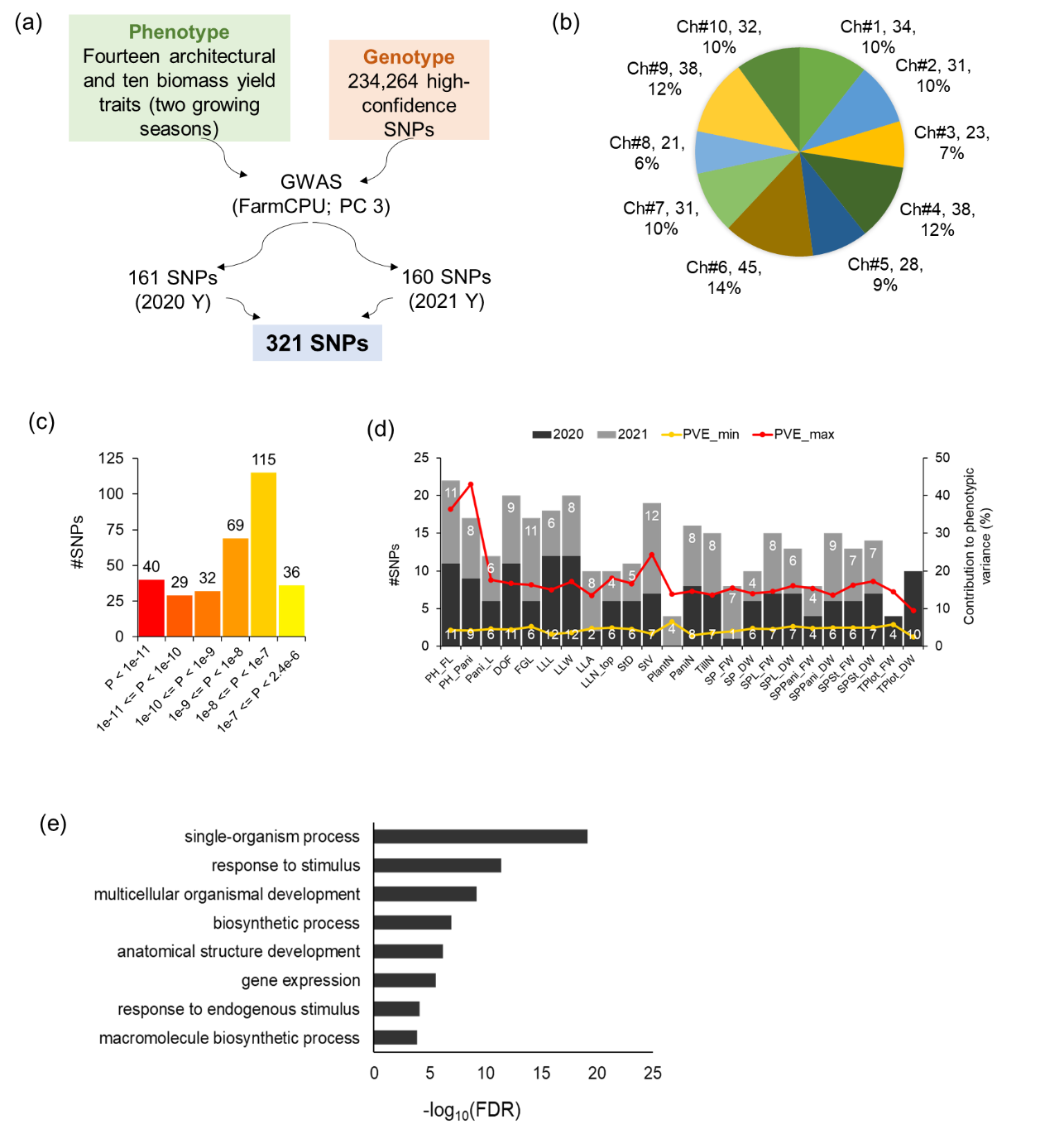

**Figure S4. Summary of SNPs identified from Genome-wide association studies (GWAS) for multiple plant architectural and biomass yield traits in sorghum over two growing seasons.**

(a) Total number of identified SNP through FarmCPU-based GWAS method, **(b)** Chromosome-wise SNP distribution, and **(c)** SNP partitioning based on –log_10_(P-values).
(b) Traits-wise SNPs distribution over two growing seasons. The plot also showed the percentage of phenotypic variance minimum (PVP_min) and maximum (PVP_max) explained by small- and large effect SNPs for each trait.
(c) Gene ontology enrichment analysis of co-localized genes around 321 SNPs.

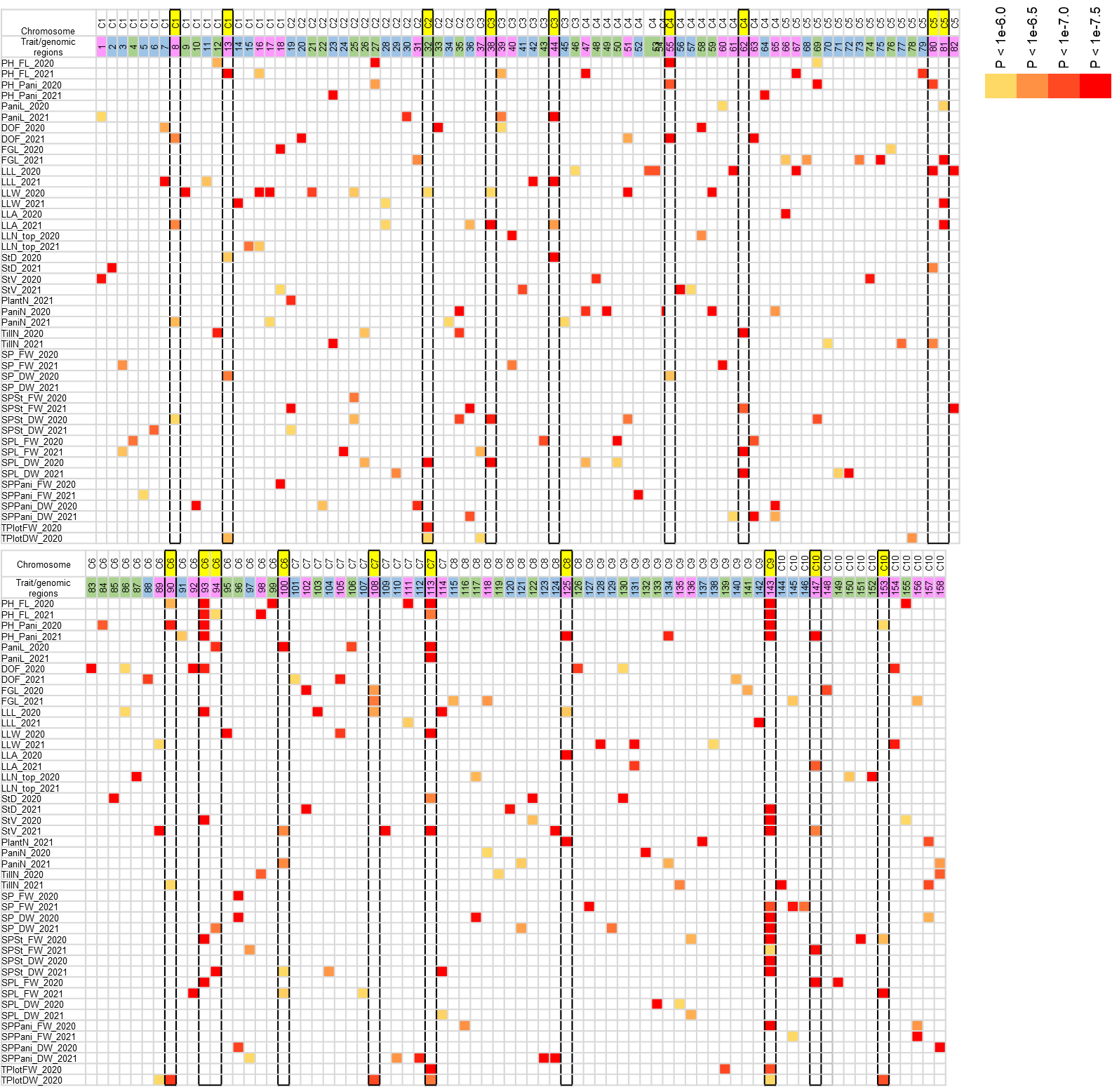

**Figure S5. Distribution of significant SNPs in various genomic regions.** These genomic regions were marked as 1 to 158, distributed across all the 10 sorghum chromosomes. The significance of each SNP was color-coded based on P-value of the association, with white boxes representing no significant association. Genomic regions highlighted in light green and light blue color represent significant SNPs detected in 2020 and 2021, respectively. Pink-colored genomic regions indicate shared significant region between the phenotypic traits of both years. Solid boxes with yellow highlighting around specific genomic regions indicate hotspot areas associated with no fewer than four traits, including plant architecture, biomass, or both. Further detail of these genomic regions was mentioned in Table S3.

| Chromosome 1 | Chromosome 2 |
| --- | --- |
| 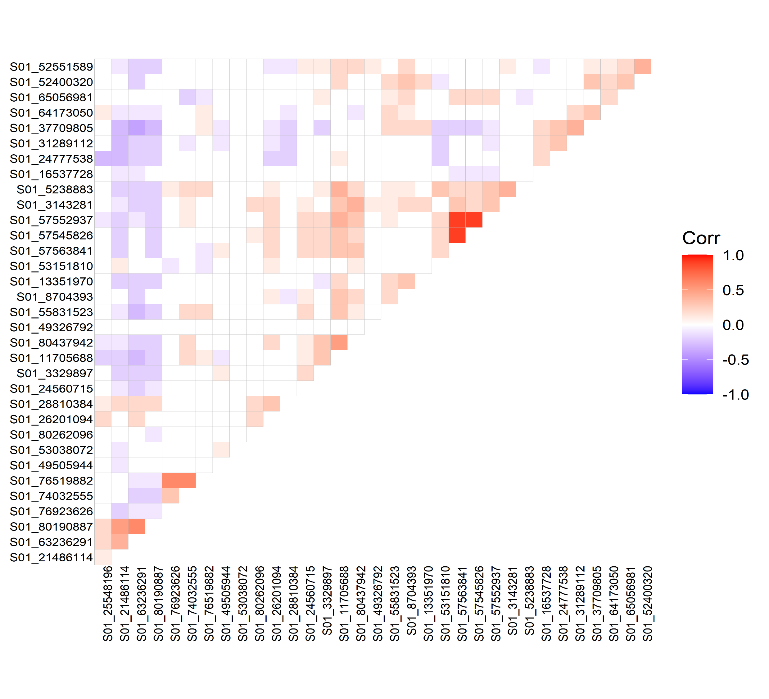 | 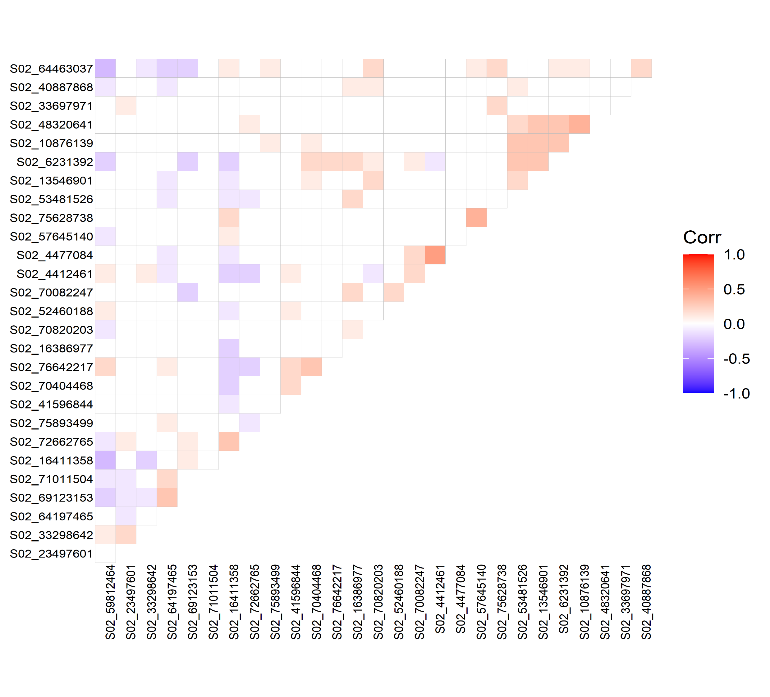 |
| Chromosome 3 | Chromosome 4 |
| 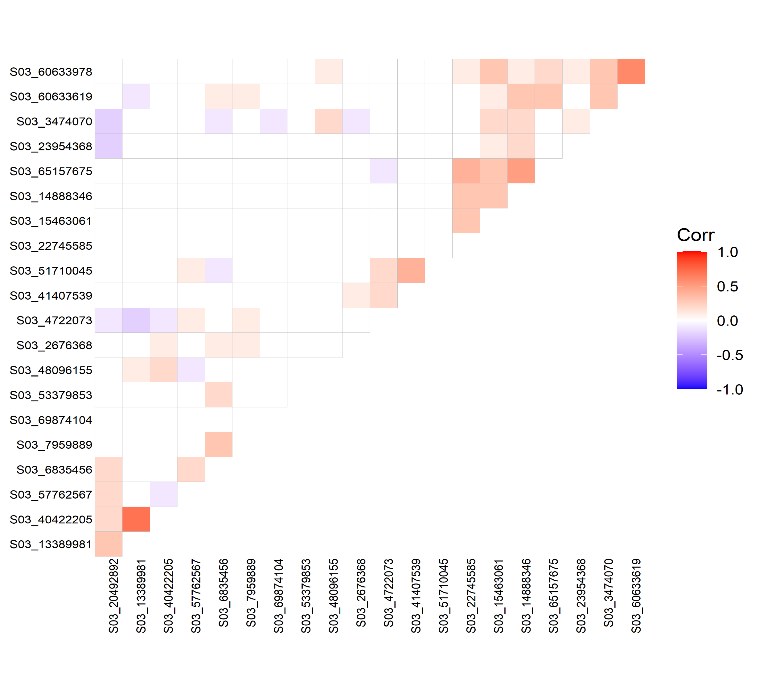 | 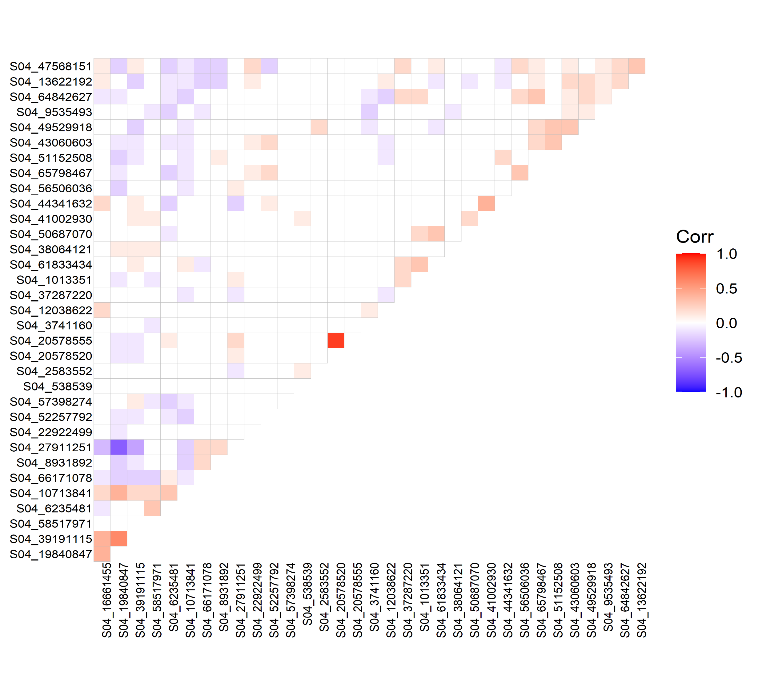 |
| Chromosome 5 | Chromosome 6 |
| 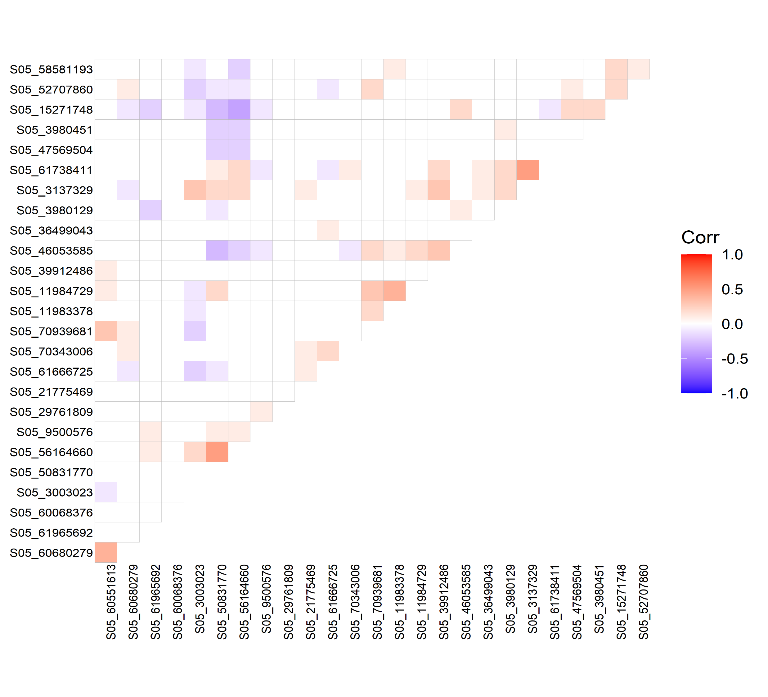 | 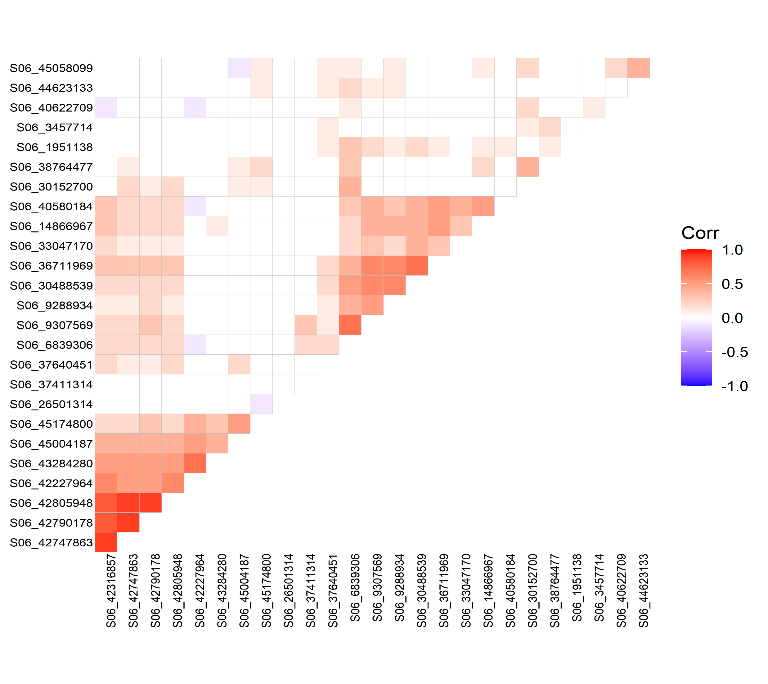 |
| Chromosome 7 | Chromosome 8 |
| 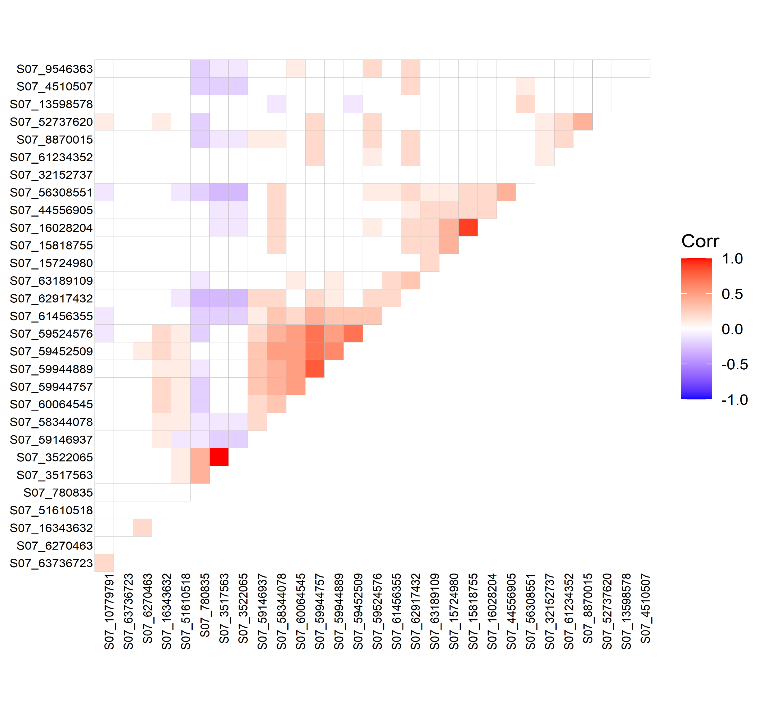 | 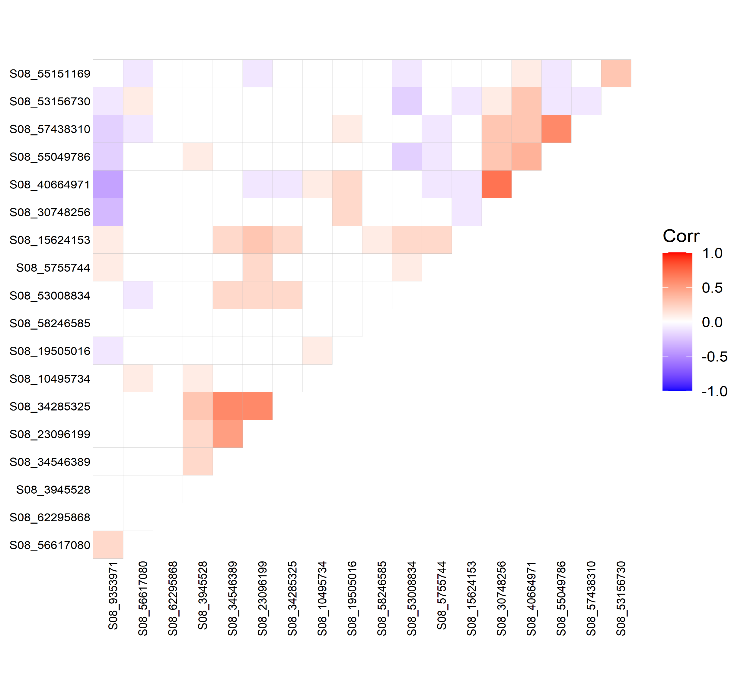 |
| Chromosome 9 | Chromosome 10 |
| 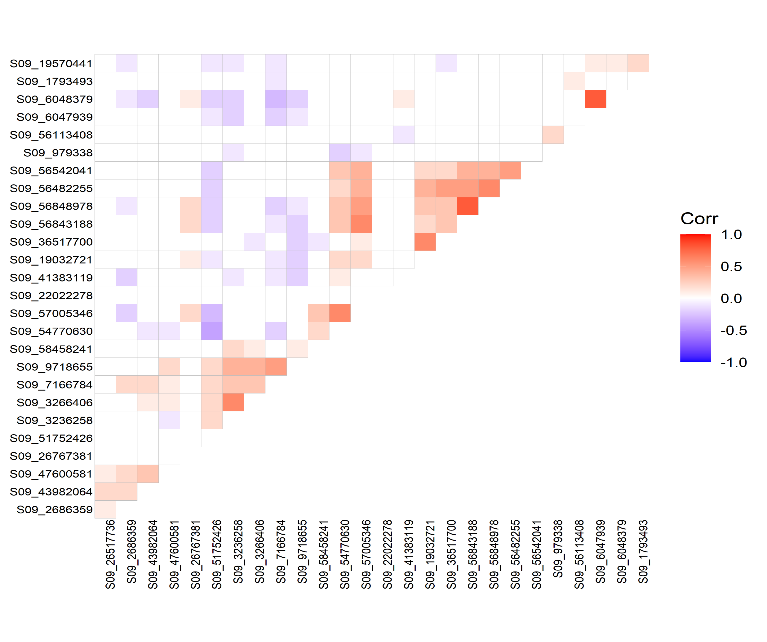 | 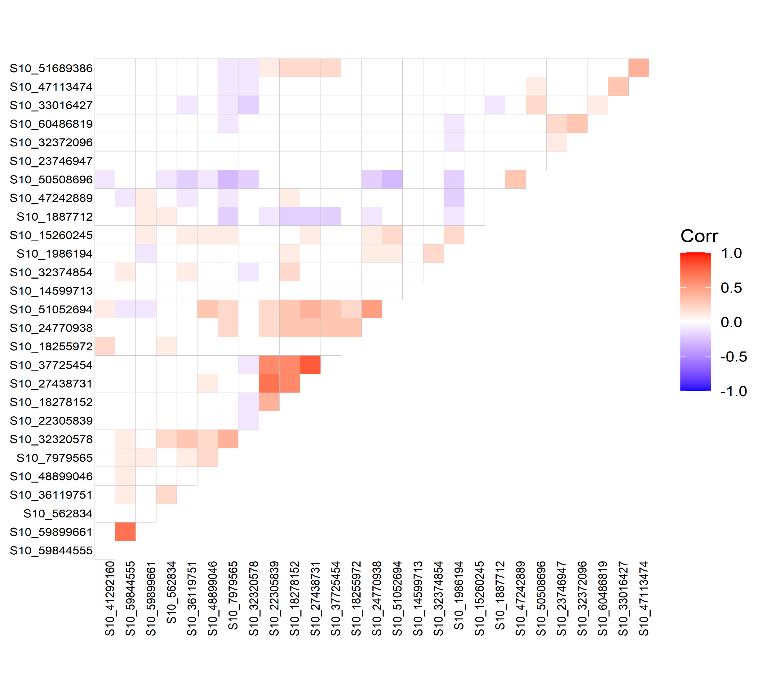 |

**Figure S6. Chromosome-wise correlation analysis among SNPs.** Positive correlations were indicated in red, while negative correlations were shown in blue. All correlations, whether positive or negative, are considered significant at P < 0.05, while non-significant correlations denoted as ‘white’. Purple boxes showed the SNP blocks with positive associations.

**Table S1. One way analysis (ANOVA) of Plant Architectural- and biomass yield traits. This was modelled using R code.**

res.aov2 <- aov(Trait of interest ~ Year + Accession)

Signif. codes: 0 ‘***’ 0.001 ‘**’ 0.01 ‘*’ 0.05 ‘.’ 0.1 ‘ ’ 1

| **Trait** |  | **Df**  **(n-1)** | **Sum Sq**  **(SS)** | **Mean Sq (SS/df)** | **F value** | **Pr (>F)** |
| --- | --- | --- | --- | --- | --- | --- |
| PH_FL | Year | 1 | 333.3 | 333.3 | 2.5 | 1.14E-01 |
|  | Accession | 403 | 1610419.7 | 3996.1 | 30.1 | 4.00E-175 *** |
|  | Residuals | 379 | 50380.7 | 132.9 |  |  |
| PH_Pani | Year | 1 | 18445.2 | 18445.2 | 125.5 | 2.37E-25 *** |
|  | Accession | 403 | 1781863.7 | 4421.5 | 30.1 | 3.60E-175 *** |
|  | Residuals | 379 | 55711.2 | 147.0 |  |  |
| Pani_L | Year | 1 | 13819.5 | 13819.5 | 450.6 | 1.88E-66 *** |
|  | Accession | 403 | 85542.3 | 212.3 | 6.9 | 1.47E-70 *** |
|  | Residuals | 379 | 11624.6 | 30.7 |  |  |
| DOF | Year | 1 | 33251.4 | 33251.4 | 1964.6 | 5.13E-152 *** |
|  | Accession | 403 | 42657.0 | 105.8 | 6.3 | 2.32E-64 *** |
|  | Residuals | 379 | 6414.8 | 16.9 |  |  |
| FGL | Year | 1 | 272.9 | 272.9 | 167.1 | 6.50E-32 *** |
|  | Accession | 403 | 2227.4 | 5.5 | 3.4 | 1.50E-31 *** |
|  | Residuals | 379 | 619.0 | 1.6 |  |  |
| LLL | Year | 1 | 2287.7 | 2287.7 | 72.3 | 4.26E-16 *** |
|  | Accession | 403 | 62498.2 | 155.1 | 4.9 | 2.64E-50 *** |
|  | Residuals | 379 | 11987.7 | 31.6 |  |  |
| LLW | Year | 1 | 203.2 | 203.2 | 304.3 | 1.88E-50 *** |
|  | Accession | 403 | 1081.6 | 2.7 | 4.0 | 8.02E-40 *** |
|  | Residuals | 379 | 253.0 | 0.7 |  |  |
| LLA | Year | 1 | 1024300.6 | 1024300.6 | 258.4 | 1.06E-44 *** |
|  | Accession | 403 | 6597901.1 | 16372.0 | 4.1 | 3.43E-41 *** |
|  | Residuals | 379 | 1502412.2 | 3964.1 |  |  |
| LLN_top | Year | 1 | 107.6 | 107.6 | 125.6 | 2.25E-25 *** |
|  | Accession | 403 | 788.4 | 2.0 | 2.3 | 5.64E-16 *** |
|  | Residuals | 379 | 324.6 | 0.9 |  |  |
| StD | Year | 1 | 3351.4 | 3351.4 | 733.2 | 1.27E-90 *** |
|  | Accession | 403 | 8052.2 | 20.0 | 4.4 | 3.98E-44 *** |
|  | Residuals | 379 | 1732.4 | 4.6 |  |  |
| StV | Year | 1 | 3527644.7 | 3527644.7 | 445.7 | 5.72E-66 *** |
|  | Accession | 403 | 16715486.5 | 41477.6 | 5.2 | 5.00E-54 *** |
|  | Residuals | 379 | 2999615.5 | 7914.6 |  |  |
| PlantN | Year | 1 | 1454.5 | 1454.5 | 280.1 | 1.83E-47 *** |
|  | Accession | 403 | 2865.9 | 7.1 | 1.4 | 9.96E-04 *** |
|  | Residuals | 379 | 1968.4 | 5.2 |  |  |
| PaniN | Year | 1 | 2024.0 | 2024.0 | 96.0 | 2.38E-20 *** |
|  | Accession | 403 | 28109.0 | 69.7 | 3.3 | 1.61E-30 *** |
|  | Residuals | 379 | 7991.6 | 21.1 |  |  |
| TillN | Year | 1 | 26.5 | 26.5 | 34.0 | 1.17E-08 *** |
|  | Accession | 403 | 767.1 | 1.9 | 2.4 | 2.41E-18 *** |
|  | Residuals | 379 | 294.8 | 0.8 |  |  |
| **Plant biomass traits** | | | | | | |
| SP_FW | Year | 1 | 2002022.1 | 2002022.1 | 262.5 | 3.09E-45 *** |
|  | Accession | 403 | 12440614.6 | 30870.0 | 4.0 | 3.58E-40 *** |
|  | Residuals | 379 | 2890217.3 | 7625.9 |  |  |
| SP_DW | Year | 1 | 2720.9 | 2720.9 | 3.0 | 8.60E-02 |
|  | Accession | 403 | 1629141.4 | 4042.5 | 4.4 | 1.69E-44 *** |
|  | Residuals | 379 | 348027.8 | 918.3 |  |  |
| SPSt_FW | Year | 1 | 510859.5 | 510859.5 | 252.4 | 6.32E-44 *** |
|  | Accession | 403 | 3856095.7 | 9568.5 | 4.7 | 2.58E-48 *** |
|  | Residuals | 379 | 767023.0 | 2023.8 |  |  |
| SPSt_DW | Year | 1 | 12852.0 | 12852.0 | 57.9 | 2.22E-13 *** |
|  | Accession | 403 | 546434.6 | 1355.9 | 6.1 | 6.08E-63 *** |
|  | Residuals | 379 | 84139.1 | 222.0 |  |  |
| SPL_FW | Year | 1 | 118180.4 | 118180.4 | 358.7 | 9.09E-57 *** |
|  | Accession | 403 | 343865.3 | 853.3 | 2.6 | 2.05E-20 *** |
|  | Residuals | 379 | 124875.3 | 329.5 |  |  |
| SPL_DW | Year | 1 | 3875.9 | 3875.9 | 107.1 | 2.90E-22 *** |
|  | Accession | 403 | 40453.8 | 100.4 | 2.8 | 4.81E-23 *** |
|  | Residuals | 379 | 13721.8 | 36.2 |  |  |
| SPPani_FW | Year | 1 | 127032.1 | 127032.1 | 44.3 | 1.01E-10 *** |
|  | Accession | 403 | 2979276.2 | 7392.7 | 2.6 | 3.28E-20 *** |
|  | Residuals | 379 | 1087830.6 | 2870.3 |  |  |
| SPPani_DW | Year | 1 | 15242.9 | 15242.9 | 35.5 | 5.94E-09 *** |
|  | Accession | 403 | 421632.7 | 1046.2 | 2.4 | 3.73E-18 *** |
|  | Residuals | 379 | 162915.2 | 429.9 |  |  |
| TPlot_FW | Year | 1 | 120.5 | 120.5 | 7.4 | 6.88E-03 ** |
|  | Accession | 403 | 30325.4 | 75.2 | 4.6 | 5.68E-47 *** |
|  | Residuals | 379 | 6183.9 | 16.3 |  |  |
| TPlot_DW | Year | 1 | 3.7 | 3.7 | 1.7 | 1.90E-01 |
|  | Accession | 403 | 3141.9 | 7.8 | 3.6 | 6.19E-35 *** |
|  | Residuals | 379 | 812.1 | 2.1 |  |  |

**Table S2. List of genes with SNPs present in the genomic regions.**

| **Trait** | **SNP** | **REF/ALT allele** | **Colocalize gene** | **Gene start** | **Gene end** | **Annotation** |
| --- | --- | --- | --- | --- | --- | --- |
| **DOF** | S02_6231392 | G/A | Sobic.002G063900 | 6225486 | 6232403 | Zinc ion binding protein |
|  | S04_57398274 | G/A | Sobic.004G223300 | 57381899 | 57399265 | ARF GTPase-activating domain-containing protein |
|  | S06_1951138 | T/G | Sobic.006G014000 | 1950491 | 1953090 | Erythronate-4-phosphate dehydrogenase |
|  | S07_8870015 | C/G | Sobic.007G076600 | 8865541 | 8870462 | RNA-dependent RNA polymerase |
|  | S08_62295868 | A/C | Sobic.008G188600 | 62295656 | 62302002 | Glyoxalase family protein |
|  | S09_3236258 | C/T | Sobic.009G035300 | 3236236 | 3237381 | NA |
| **Leaf related traits** | | | | | | |
| LLN_top | S01_57563841 | T/G | Sobic.001G296400 | 57563837 | 57580610 | Palmitoyl-protein thioesterase 1 precursor |
| LLL | S01_21486114 | C/A | Sobic.001G224300 | 21485582 | 21489706 | CESA7 - cellulose synthase |
| LLW | S01_55831523 | C/G | Sobic.001G285100 | 55829290 | 55832731 | NA |
| LLW | S01_76519882 | C/T | Sobic.001G495600 | 76517358 | 76520284 | Plant-specific domain TIGR01615 family protein |
| LLA | S02_53481526 | G/A | Sobic.002G169400 | 53476672 | 53486427 | Leucine Rich Repeat family protein |
| LLA | S03_4722073 | C/G | Sobic.003G052300 | 4721513 | 4725477 | Glycosyltransferase |
| LLA/LLA | S03_60633978 | G/T | Sobic.003G269600 | 60633944 | 60636136 | Cytochrome P450 |
| SPL_FW/DW | S04_6235481 | C/T | Sobic.004G076400 | 6234594 | 6237205 | Zinc-binding protein |
| SPL_FW | S04_58517971 | T/C | Sobic.004G237300 | 58513994 | 58520353 | TCP-domain protein |
| LLW | S06_46372143 | C/G | Sobic.006G093600 | 46371335 | 46373767 | NA |
| FGL | S05_61666725 | T/C | Sobic.005G148500 | 61659987 | 61667430 | OsFBX402 |
| LLW | S07_58344078 | G/T | Sobic.007G151400 | 58341777 | 58345411 | Cytokinin dehydrogenase precursor |
| FGL | S10_18278152 | G/T | Sobic.010G132100 | 18274379 | 18282651 | Thioredoxin domain-containing protein 17 |
| FGL | S10_1986194 | A/G | Sobic.010G023900 | 1984187 | 1986475 | S10/S20 domain containing ribosomal protein |
| **Plant height and stem related trait** | | | | | | |
| StV | S01_3143281 | T/C | Sobic.001G042200 | 3142618 | 3149897 | ATP/GTP/Ca++ binding protein |
| SP_FW | S01_8704393 | G/T | Sobic.001G111500 | 8708287 | 8721152 | phytochrome A |
| PH_FL | S04_1013351 | G/A | Sobic.004G012300 | 1019939 | 1026858 | transposon protein |
| PH_FL | S04_1013351 | G/A | Sobic.004G012250 | 1013937 | 1016535 | NA |
| StV | S04_2583552 | A/G | Sobic.004G031900 | 2581116 | 2583897 | phosphatidylinositol transfer |
| PH_Pani | S05_60680279 | C/G | Sobic.005G143200 | 60675665 | 60681004 | transposon protein |
| PH_Pani | S05_60680279 | C/G | Sobic.005G143300 | 60680201 | 60688234 | NA |
| PH_Pani | S06_38764477 | G/A | Sobic.006G053601 | 38765522 | 38769204 |  |
| PH_FL/PH_Pani | S06_42790178 | T/C | Sobic.006G067600 | 42785485 | 42802516 | histone deacetylase |
| StV | S06_42805948 | T/C | Sobic.006G067700 | 42803037 | 42807134 | Protein kinase |
| PH_FL | S06_45058099 | T/C | Sobic.006G082100 | 45062982 | 45068690 | dehydration-responsive element-binding protein |
| SP_DW | S06_45174800 | G/A | Sobic.006G083300 | 45201363 | 45228930 | DEAD/DEAH box helicase domain containing protein |
| SP_FW/DW | S06_48856570 | C/G | Sobic.006G122400 | 48852708 | 48856765 | uncharacterized glycosyltransferase |
| PH_FL | S07_59944757 | T/C | Sobic.007G164400 | 59944610 | 59946235 | inorganic phosphate transporter |
| SPSt_DW | S07_63736723 | A/C | Sobic.007G208300 | 63740858 | 63745071 | peptidase |
| PH_Pani | S08_58246585 | A/G | Sobic.008G149200 | 58248671 | 58251403 | vegetative storage protein |
| SP_DW | S09_2686359 | G/A | Sobic.009G030200 | 2690764 | 2697614 | homeobox protein knotted-1-like 6 |
| SP_DW/SPSt_DW | S09_56542041 | G/A | Sobic.009G222800 | 56542104 | 56544279 | NA |
| SP_FW/ SPSt_FW | S09_58458241 | C/G | Sobic.009G249400 | 58459026 | 58461413 | protein kinase family protein |
| SP_FW/ SPSt_FW | S09_58458241 | C/G | Sobic.009G249500 | 58461666 | 58466785 | mitochondrial carrier protein |
| SP_FW | S09_979338 | G/C | Sobic.009G010600 | 980050 | 992263 | retinoblastoma-related protein-like |
| SP_FW | S10_1887712 | C/A | Sobic.010G023300 | 1888726 | 1890513 |  |
| PH_FL/StV | S10_41292160 | A/C | Sobic.010G145800 | 41279629 | 41293580 | KH domain containing protein |
| SP_FW | S10_7979565 | C/T | Sobic.010G091400 | 7989858 | 8021963 | AAA-type ATPase family protein |
| **Panicle and tiller related trait** | | | | | | |
| SPPani_FW | S01_13351970 | G/T | Sobic.001G162200 | 13355784 | 13364699 | NA |
| SPPani_DW | S02_13546901 | A/C | Sobic.002G111400 | 13545427 | 13547706 | Heavy metal-associated domain |
| TillN | S02_16386977 | A/G | Sobic.002G123000 | 16395494 | 16406892 | NA |
| Tplot_FW | S02_70820203 | G/T | Sobic.002G341400 | 70819427 | 70821008 | NA |
| TillN | S02_76642217 | A/G | Sobic.002G418600 | 76633446 | 76644644 | SNF2 family N-terminal domain |
| Pani_L | S03_57762567 | T/C | Sobic.003G238000 | 57764407 | 57768648 | Ras-related protein |
| Tplot_DW | S03_6835456 | C/G | Sobic.003G079700 | 6833746 | 6837627 | Ser/Thr protein kinase |
| PaniN | S04_3741160 | C/G | Sobic.004G045566 | 3740988 | 3744736 | Ras-related protein |
| SPPani_DW | S04_64842627 | T/C | Sobic.004G311750 | 64843889 | 64846480 | Transposon protein |
| PaniN | S04_65798467 | A/T | Sobic.004G323200 | 65797909 | 65798517 | SCP-like extracellular protein |
| Pani_L | S06_45004187 | A/T | Sobic.006G081200 | 45003932 | 45007305 | Phospholipase C |
| SPPani_DW | S06_48853118 | A/G | Sobic.006G122400 | 48852708 | 48856765 | Uncharacterized glycosyltransferase |
| TillN | S06_53273900 | C/A | Sobic.006G178300 | 53272301 | 53274151 | GDSL-like lipase/acylhydrolase |
| PaniN | S06_59255525 | G/T | Sobic.006G255300 | 59259860 | 59266545 | auxin response factor |
| SPPani_DW | S08_55151169 | C/T | Sobic.008G126700 | 55154011 | 55158809 | calmodulin binding protein |
| PlantN | S08_56617080 | G/A | Sobic.008G137200 | 56618112 | 56620952 | Mur ligase family protein |
| PlantN | S08_56617080 | G/A | Sobic.008G137100 | 56617223 | 56617624 | RALFL42 |
| SPPani_FW | S10_1887712 | C/A | Sobic.010G023300 | 1888726 | 1890513 |  |
| SPPani_FW | S10_47242889 | G/A | Sobic.010G160700 | 47244445 | 47253575 | phosphoenolpyruvate carboxylase |
| PaniN | S10_59844555 | G/A | Sobic.010G261900 | 59847394 | 59851087 | methyltransferase |
| SPPani_DW | S10_60486819 | G/A | Sobic.010G270700 | 60480062 | 60488603 | GDSL-like lipase/acylhydrolase |

**Table S3. Summary of genomic regions with significant associations displaying multiple SNPs in linkage disequilibrium.** The physical position of the genomic region is based on the sorghum reference genome v3.1.1. The graphical representation of these SNPs in identified genomic regions was shown in Figure S5.

| Genomic region (GR)  (Start-end) | #Ch | GR ID |  | Genomic region (GR)  (Start-end) | #Ch | GR ID |
| --- | --- | --- | --- | --- | --- | --- |
| 3143281 - 3329897 | 1 | 1 |  | 48096155 - 48444155 | 3 | 41 |
| 5238883 - 5568883 | 1 | 2 |  | 51710045 - 52200045 | 3 | 42 |
| 8704393 - 9004393 | 1 | 3 |  | 53379853 - 53809853 | 3 | 43 |
| 11705688 - 12055688 | 1 | 4 |  | 57762567 - 60633978 | 3 | 44 |
| 13351970 - 13706970 | 1 | 5 |  | 65157675 - 65478675 | 3 | 45 |
| 16537728 - 16897728 | 1 | 6 |  | 69874104 - 70195104 | 3 | 46 |
| 21486114 - 21861114 | 1 | 7 |  | 538539 - 1013351 | 4 | 47 |
| 24560715 - 25201094 | 1 | 8 |  | 2583552 - 2803552 | 4 | 48 |
| 28810384 - 29470384 | 1 | 9 |  | 3741160 - 4046160 | 4 | 49 |
| 31289112 - 31841112 | 1 | 10 |  | 6235481 - 6560481 | 4 | 50 |
| 37709805 - 38135805 | 1 | 11 |  | 8931892 - 10713841 | 4 | 51 |
| 49326792 - 49505944 | 1 | 12 |  | 12038622 - 12318622 | 4 | 52 |
| 52400320 - 53151810 | 1 | 13 |  | 13622192 - 13932192 | 4 | 53 |
| 55831523 - 56261523 | 1 | 14 |  | 16661455 - 16966455 | 4 | 54 |
| 57545826 - 57563841 | 1 | 15 |  | 19840847 - 20578555 | 4 | 55 |
| 63236291 - 65056981 | 1 | 16 |  | 22922499 - 23247499 | 4 | 56 |
| 74032555 - 76923626 | 1 | 17 |  | 27911251 - 28256251 | 4 | 57 |
| 80190887 - 80437942 | 1 | 18 |  | 37287220 - 39191115 | 4 | 58 |
| 4412461 - 4477084 | 2 | 19 |  | 41002930 - 41316930 | 4 | 59 |
| 6231392 - 6601392 | 2 | 20 |  | 43060603 - 44341632 | 4 | 60 |
| 10876139 - 11241139 | 2 | 21 |  | 47568151 - 49529918 | 4 | 61 |
| 13546901 - 14036901 | 2 | 22 |  | 50687070 - 52257792 | 4 | 62 |
| 16386977 - 16411358 | 2 | 23 |  | 56506036 - 58517971 | 4 | 63 |
| 23497601 - 23796601 | 2 | 24 |  | 61833434 - 62153434 | 4 | 64 |
| 33298642 - 33697971 | 2 | 25 |  | 64842627 - 66171078 | 4 | 65 |
| 40887868 - 41596844 | 2 | 26 |  | 3003023 - 3137329 | 5 | 66 |
| 48320641 - 48635641 | 2 | 27 |  | 3980129 - 4169129 | 5 | 67 |
| 52460188 - 53481526 | 2 | 28 |  | 9500576 - 9731026 | 5 | 68 |
| 57645140 - 57951140 | 2 | 29 |  | 11983378 - 12195051 | 5 | 69 |
| 59812464 - 60133464 | 2 | 30 |  | 15271748 - 15583648 | 5 | 70 |
| 64197465 - 64463037 | 2 | 31 |  | 21775469 - 22008469 | 5 | 71 |
| 69123153 - 70820203 | 2 | 32 |  | 29761809 - 29960809 | 5 | 72 |
| 71011504 - 71344504 | 2 | 33 |  | 36499043 - 36800043 | 5 | 73 |
| 72662765 - 73007765 | 2 | 34 |  | 39912486 - 40237409 | 5 | 74 |
| 75628738 - 76642217 | 2 | 35 |  | 46053585 - 46289263 | 5 | 75 |
| 2676368 - 4722073 | 3 | 36 |  | 47569504 - 47881849 | 5 | 76 |
| 6835456 - 7959889 | 3 | 37 |  | 50831770 - 51140870 | 5 | 77 |
| 13389981 - 15463061 | 3 | 38 |  | 52707860 - 53008727 | 5 | 78 |
| 20492892 - 23954368 | 3 | 39 |  | 56164660 - 56420895 | 5 | 79 |
| 40422205 - 41407539 | 3 | 40 |  | 58581193 - 60680279 | 5 | 80 |

| Genomic region (GR)  (Start-end) | #Ch | GR ID |  | Genomic region (GR)  (Start-end) | #Ch | GR ID |
| --- | --- | --- | --- | --- | --- | --- |
| 61666725 - 61965692 | 5 | 81 |  | 34285325 - 34546389 | 8 | 121 |
| 70343006 - 70939681 | 5 | 82 |  | 40664971 - 40896861 | 8 | 122 |
| 1951138 - 2149703 | 6 | 83 |  | 53008834 - 53156730 | 8 | 123 |
| 3457714 - 3702714 | 6 | 84 |  | 55049786 - 55151169 | 8 | 124 |
| 6839306 - 7028980 | 6 | 85 |  | 56617080 - 58246585 | 8 | 125 |
| 9288934 - 9521934 | 6 | 86 |  | 62295868 - 62507542 | 8 | 126 |
| 14866967 - 15048640 | 6 | 87 |  | 979338 - 1184017 | 9 | 127 |
| 26501314 - 26690437 | 6 | 88 |  | 1793493 - 2028171 | 9 | 128 |
| 30152700 - 30357398 | 6 | 89 |  | 2686359 - 2897922 | 9 | 129 |
| 36711969 - 36946536 | 6 | 90 |  | 3236258 - 3503690 | 9 | 130 |
| 38764477 - 38962714 | 6 | 91 |  | 6047939 - 6270506 | 9 | 131 |
| 40580184 - 40797643 | 6 | 92 |  | 7166784 - 7412017 | 9 | 132 |
| 42227964 - 42444423 | 6 | 93 |  | 9718655 - 9920309 | 9 | 133 |
| 44623133 - 44869089 | 6 | 94 |  | 19032721 - 19570441 | 9 | 134 |
| 46372143 - 46539977 | 6 | 95 |  | 21628827 - 22022278 | 9 | 135 |
| 48853118 - 49099839 | 6 | 96 |  | 26517736 - 26767381 | 9 | 136 |
| 51334145 - 51858612 | 6 | 97 |  | 36517700 - 36712483 | 9 | 137 |
| 53273900 - 53620573 | 6 | 98 |  | 41383119 - 41570588 | 9 | 138 |
| 56719937 - 57087060 | 6 | 99 |  | 43982064 - 44207900 | 9 | 139 |
| 58127831 - 60840408 | 6 | 100 |  | 47600581 - 47958803 | 9 | 140 |
| 780835 - 949891 | 7 | 101 |  | 51752426 - 52119771 | 9 | 141 |
| 3517563 - 3701346 | 7 | 102 |  | 54770630 - 55175996 | 9 | 142 |
| 4510507 - 4721552 | 7 | 103 |  | 56113408 - 58458241 | 9 | 143 |
| 6270463 - 6479589 | 7 | 104 |  | 562834 - 741401 | 10 | 144 |
| 8870015 - 9546363 | 7 | 105 |  | 1887712 - 2155054 | 10 | 145 |
| 10779791 - 11000249 | 7 | 106 |  | 7979565 - 8216437 | 10 | 146 |
| 13598578 - 13810141 | 7 | 107 |  | 14599713 - 15260245 | 10 | 147 |
| 15724980 - 16343632 | 7 | 108 |  | 18255972 - 18585972 | 10 | 148 |
| 32152737 - 32409215 | 7 | 109 |  | 22305839 - 22705839 | 10 | 149 |
| 44556905 - 44793653 | 7 | 110 |  | 23746947 - 24201947 | 10 | 150 |
| 51610518 - 52737620 | 7 | 111 |  | 24770938 - 25190956 | 10 | 151 |
| 56308551 - 56553388 | 7 | 112 |  | 27438731 - 27728765 | 10 | 152 |
| 58344078 - 61456355 | 7 | 113 |  | 32320578 - 33016427 | 10 | 153 |
| 62917432 - 63736723 | 7 | 114 |  | 36119751 - 37725454 | 10 | 154 |
| 3945528 - 4157317 | 8 | 115 |  | 41292160 - 41592160 | 10 | 155 |
| 5755744 - 5989867 | 8 | 116 |  | 47113474 - 48899046 | 10 | 156 |
| 9353971 - 10495734 | 8 | 117 |  | 50508696 - 51689386 | 10 | 157 |
| 15624153 - 19505016 | 8 | 118 |  | 59844555 - 60486819 | 10 | 158 |
| 23096199 - 23295722 | 8 | 119 |  |  |  |  |
| 30748256 - 30959954 | 8 | 120 |  |  |  |  |
